# A gentle palette of plasma membrane dyes

**DOI:** 10.1101/2024.05.04.592408

**Authors:** Jing Ling, Yitong Liu, Yunzhe Fu, Shuzhang Liu, Ling Ding, Lulu Huang, Peng Xi, Zhixing Chen

## Abstract

Plasma membrane stains are one of the most important organelle markers for unambiguous assignments of individual cells and monitoring membrane morphology and dynamics. The state-of-the-art PM stains are bright, specific, fluorogenic, and compatible with super-resolution imaging. However, when recording membrane dynamics, particularly under light-intensive microscopes, PM is prone to photodynamic damages due to its phospholipid bilayer nature. Here we developed PK Mem dyes tailored for time-lapse fluorescence imaging. By integrating triplet-state quenchers into the MemBright dyes featuring cyanine chromophores and amphiphilic zwitterion anchors, PK Mem dyes exhibited a three-fold reduction in phototoxicity and a more than four-fold improvement in photostability in imaging experiments. These dyes enable 2D and 3D imaging of live or fixed cancer cell lines and a wide range of primary cells, at the same time pair well with various fluorescent markers. PK Mem dyes can be applied to neuronal imaging in brain slices and *in vivo* two-photon imaging. The gentle nature of PK Mem palette enables ultralong-term recording of cell migration and cardiomyocyte beating. Notably, PK Mem dyes are optically compatible with STED/SIM imaging, which can handily upgrade the routine of time-lapse neuronal imaging, such as growth cone tracking and mitochondrial transportations, into nanoscopic resolutions.

## Introduction

The plasma membrane (PM) is a dynamic structure separating the cell from the extracellular environment^1^ and orchestrating substance transport^2^, signal transduction^3, 4^ and cell recognition^5^. Fluorescence imaging is one of the primary ways to study PM, such as in cell division, visualization of cell morphology, tracking membrane dynamics as well as monitoring intracellular and extracellular vesicles^6^. Therefore, strategies for fluorescently highlighting plasma membranes are instrumental in the visualization of membrane dynamics.

While it is possible to target genetically encoded fluorescent proteins to PM as imaging tags, chemical dyes offer more convenient and brighter staining of the PM, leading to their wider range of applications^6-8^. Achieving high specificity is the main quest of cell membrane imaging, and various strategies have been developed toward this goal. Exploiting antigen-antibody interaction between probes and surface antigen is an effective approach, giving a family of widely-used PM stains based on wheat germ agglutinin (WGA)-dye conjugate^9, 10^. However, due to the high molecular weight, WGA-iFluor™ 488 exhibits significant variability in intercellular staining and is prone to cellular endocytosis and dye internalization^11, 12^. In addition, it’s reported that WGA-iFluor™ 488 can partially hinder the dynamics of the cell membrane^11^. Alternatively, chemical probes, bearing lower molecular weight and minimal perturbations to membranes, are developed and widely applied. These PM dyes are devised with membrane anchors and fluorophores, giving high labeling contrast through noncovalent interactions with phospholipids^13-18^. Generations of PM dyes, such as DiD, FM4-64, and Cell Mask, have been developed for cell membrane imaging^19-21^. However, each of them has certain inherent limitations. DiD and Cell Mask family dyes rely on hydrophobic interaction between long alkyl chains and cell membrane, resulting in probe internalization, high staining concentration and low solubility in working solution^18^. FM4-64 tends to translocate to the nuclear membrane at physiological temperatures due to its affinity for negatively charged lipids^22, 23^. Therefore, several strategies have been developed to address these issues. Among the emerging methods, the Klymchenko group reported the conjugation of amphiphilic zwitterionic anchors with various fluorophores, such as Nile Red^24, 25^, squaraine^26^, cyanine^12^, and BODIPY^27^ to create PM dyes capable of selectively and persistently staining cell membranes. One notable combination among them are MemBright dyes^12^ (commercial name: MemGlow), consisting of cyanine and two amphiphilic zwitterionic anchors, which have found widespread applications with confocal microscopy, two-photon microscopy, and stochastic optical reconstruction microscopy.

On the flip side of the constantly evolving PM labeling strategies is the photophysical chemistry of the core fluorophores. In the past decade, the field has witnessed the transition from widefield and confocal imaging towards super-resolution and time-lapse imaging, which can monitor cell membrane morphology and dynamics at unprecedented spatial and temporal resolutions. However, the strong excitation schemes during imaging give rise to phototoxicity and photobleaching, leading to cell death and fluorescent signal disappearance during live imaging^28-30^. The reactive oxygen species (ROS) stemmed from the photosensitized dyes and molecular oxygen is believed to account for phototoxicity and photobleaching^31-34^. The PM, bearing unsaturated lipid with vulnerable olefin groups, exhibits greater sensitivity to ROS compared to other subcellular structures such as cytoskeleton, nucleus, or lipid droplet^35, 36^. Imaging-induced phototoxicity can impair the structure and function of the cell membrane, leading to membrane blebbing, membrane rupture, and eventual cell death^37^. Consequently, gentle fluorophores with low photodamage are becoming instrumental for future live-cell recordings. Among the few chemical strategies that can alleviate phototoxicity and photobleaching^38-41^, intramolecular triplet-state quenching has stood out as one of the most promising strategies. Originally proposed and demonstrated with cyclooctatetraene (COT)-conjugated cyanine dyes and single-molecule applications^33, 39^, TSQ-dye molecules have been recently employed for live cell imaging applications including mitochondrial cristae imaging^34, 42^, voltage imaging^43, 44^, and chemigenetic protein imaging^33, 45^. COT-conjugation has generally alleviated phototoxicity during time-lapse imaging at these selected cellular targets.

Herein, we reported PK Mem dyes, a palette of gentle PM dyes (Fig. 1A). Cyanine chromophore, triplet state quenchers, and membrane anchors are elaborately integrated to PK Mem molecules, giving excellent labeling efficiency of PM, reduced phototoxicity, and enhanced photostability. PK Mem dyes can effectively stain cancer cell lines, primary cells, and brain slices with high contrast. We demonstrate advanced applications including *in vivo* two-photon imaging in a mouse brain, stimulated emission depletion (STED) imaging of dendritic spines, time-lapse tracking of migrasomes, and SIM recording of mitochondrial movement in axons. Practically, the phototoxicity of PK Mem dyes is less than one-third that of other cyanine-based PM stains, offering the monitoring of cellular processes over remarkably extended durations.

**Figure 1.**
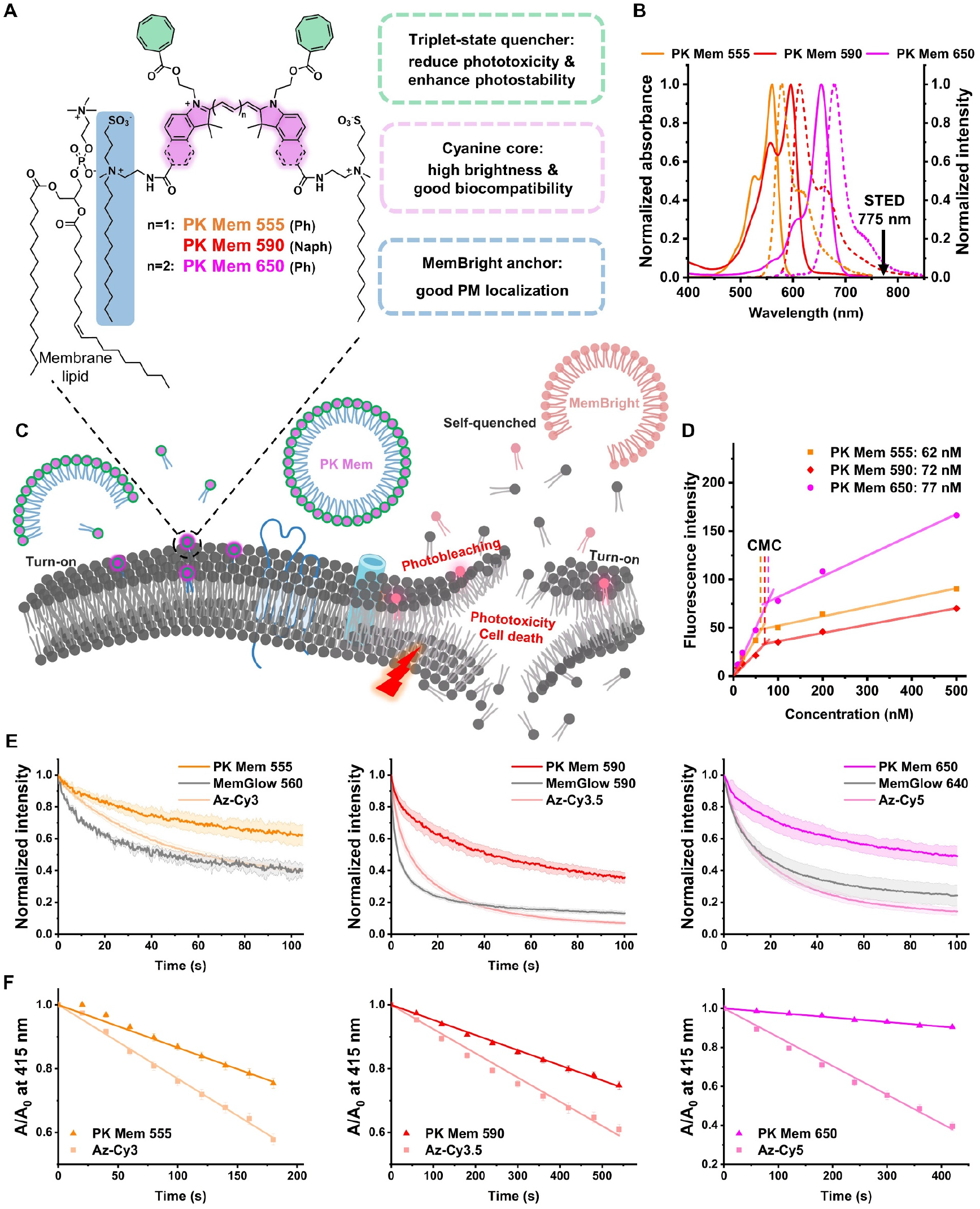
PK Mem dyes employ cyclooctatetraenes as triplet-state quenchers to optically strengthen and photodynamically tame the MemBright dyes. (A) Chemical structures of PK Mem dyes featuring the modular integration of cyclooctatetraenes and amphiphilic anchors to the cyanine palette. (B) Absorption (solid lines) and emission (dashed lines) spectra of PK Mem dyes in MeOH. (C) Schemes for the mechanisms of turn-on, photostability, and phototoxicity of PK Mem dyes. (D) Plot of the maximum fluorescence intensity against the concentration of PK Mem dyes showing their critical micelle concentration (CMC). (E) Photostability comparison among PK Mem dyes, MemGlow, and AZ-Cy labelled on fixed HeLa cells. (F) In vitro detection of reactive oxygen species (ROS) through 1,3-diphenylisobenzofuran decay assay.

## Results

### Design, synthesis, and characterizations of PK Mem dyes

The design of gentle PM dyes with good specificity and low phototoxicity hinges on the streamlined assembly of triplet-state quenchers and membrane anchors onto selected fluorophores with practical synthetic chemistry. The most established TSQ for intracellular imaging, acyl-cyclooctatetraene, and its optimal two-carbon linker to the cyanine fluorophores, were inherited from previous works^33, 34, 42^. Considering that the valency and position of membrane anchors can influence the performance of PM dyes^26, 27^, we strategically planned two anchors at the benzene or naphthalene rings to obtain symmetrical probes.

The molecular design of PK Mem dyes is substantiated with practical synthetic chemistry within 7 steps from commercial materials. Take PK Mem 590 as an example (Scheme 1), the substituted naphthylhydrazine **2**, produced from commercial material **1**, was converted into intermediate **3** through Fischer indole synthesis. The key intermediate, fluorophore **5**, was synthesized from **3** through alkylation with iodoethanol followed by condensation with diphenylformamidine. The membrane anchor motif was prepared via alkylation and deprotection reactions on precursor **6**, giving a primary amine, **7. 7** was grafted onto fluorophore **5** by amide bonds to give intermediate **8**. Subsequently, compound **8** was alkylated with 1,3-propanesultone to yield the amphiphilic intermediate **9**, thereby completing the installations of two membrane anchors. Finally, cyclooctatetraenecarboxylic acid (COT-COOH) was coupled to the fluorophore, giving PK Mem 590. Likewise, PK Mem 555, 650, and their non-COT counterparts, Az-Cy dyes, were prepared through similar routes (Fig. S1 and Supplementary scheme 1-6).

With the PK Mem dyes in hand, their absorption and emission spectra in different solvents were recorded (Fig. 1B and Table S1). Resembling MemBright dyes, the fluorescence quantum yields of PK Mem dyes experience a drastic decrease in PBS and a marked increase in DOPC liposomes (Table S1), indicating that the amphiphilic PK Mem dyes could form micelles in aqueous solutions, leading to strong self-quenching in fluorescence. The aggregates of PK Mem dyes undergo disassembly within lipid membranes (Fig. 1C), giving a fluorogenic response to liposomes and cells. The critical micelles concentrations of PK Mem dyes were determined to be approximately 70 nM (Fig. 1D). Compared to micelles in PBS, the fluorescence quantum yields of PK Mem 555, 590, and 650 in DOPC liposomes increased 24-fold, 41-fold, and 180-fold, respectively, demonstrating high fluorogenicity and the potential for wash-free imaging (Table S1).

To investigate the impact of the triplet-state quencher COT on phototoxicity and photostability, PK Mem dyes and MemGlow dyes, bearing the same fluorophores and anchors but with or without COTs were compared in a head-to-head manner. Az-Cy dyes, featuring different connectivity between chromophores and anchors, were also compared. First, photobleaching curves were obtained from imaging fixed HeLa cells to avoid membrane motion artifacts. The bleaching half-life (τ_1/2_) of PK Mem dyes were 4.0-fold, 14.3-fold, and 5.6-fold longer than that of MemGlow dyes, and 3.2-fold, 5.4-fold, and 6.7-fold longer than that of Az-Cy3/3.5/5, respectively (Fig. 1E and Table S2). Furthermore, their reactive oxygen species (ROS) generations were measured using the 1,3-diphenylisobenzofuran decay assay^46^. Compared to Az dyes, the singlet oxygen quantum yields of PK Mem dyes decreased by 22-31% after conjugation with COT (Fig. 1F and Table S2). These data suggested that COT conjugation to membrane dyes exhibited alleviated phototoxicity and photobleaching.

### Imaging of cancer and primary cells

Having the gentle dyes characterized in vitro, we conducted cell labeling experiments to test the specificity of PK Mem dyes on various cells (Fig. 2A). After stained with PK Mem dyes (20 nM) for 5 min, HeLa cells were imaged under confocal microscopy without washing. PK Mem dyes exhibited distinct and intense localization on cell membranes, and the staining process was remarkably rapid (Fig. 2B). In a comparative co-staining experiment, PK Mem dyes achieved specific and bright membrane localization comparable to that of WGA-iFluor™ 488 within 5 min (Fig. S2). PK Mem probes exhibit minimal internalization within 120 min at 37? (Fig. S3). In confluent HeLa cells co-stained with PK Mem 650 and WGA-iFluor™ 488, the ratio of PK Mem 650 fluorescence intensity to WGA-488 fluorescence intensity was quantified at plasma membrane, cell-cell contacts, and tunneling nanotubes. At cell-cell contacts, PK Mem 650 exhibits approximately two-fold stronger labeling compared to WGA-488 (Fig. S4). By principle, WGA-488 labels the cell membrane through surface polysaccharides, resulting in variations between individual cells. Yet, WGA staining is more sensitive to accessibility due to its bulkier size, giving weaker signal at contact sites. At tunneling nanotubes, PK Mem 650 exhibited over two-fold higher staining selectivity than WGA-488 (Fig. S4). The impact of the probes on the viability of HeLa cells was assessed via CCK-8 assay. PK Mem dyes did not exhibit significant cytotoxicity at a concentration as high as 1 μM which is far above their staining concentrations (Fig. S5). Finally, compatibility with fixation was tested. HeLa cells were either stained after fixation (Fig. S6A) in 4% formaldehyde in PBS or fixed subsequent to staining (Fig. S6B). In both cases specific PM staining can be achieved.

**Figure 2.**
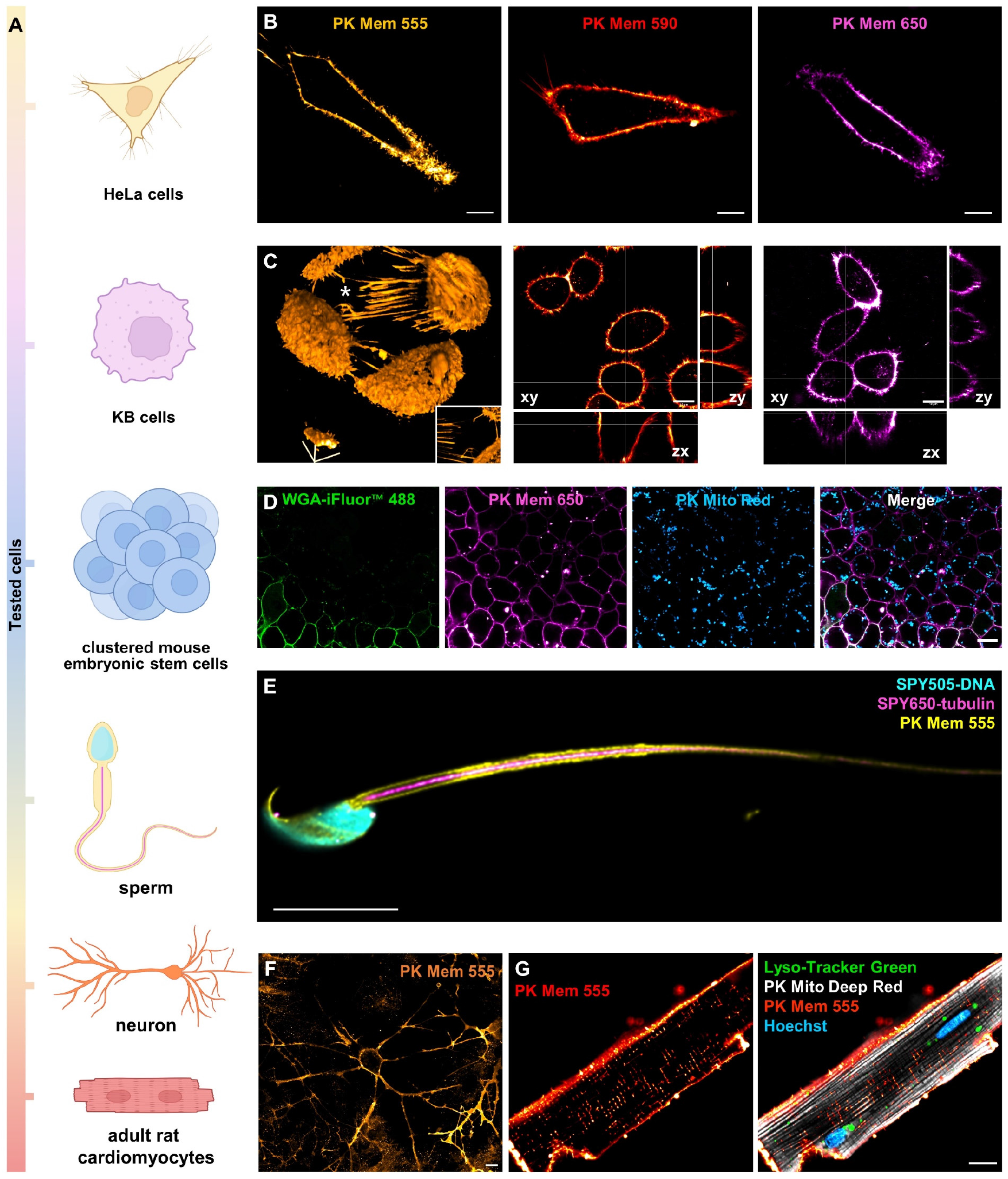
2D and 3D confocal imaging of various cells using PK Mem dyes. (A) Cancer and primary cells imaged in this study. (B) Laser scanning confocal microscopy (LSCM) images of live HeLa cells treated with PK Mem dyes (20 nM) for 5 min without washing. Scale bar = 10 μm. (C) Left, 3D reconstruction of live KB cells stained with PK Mem dyes. Inset in the left panel is the top view of intercellular nanotubes indicated by an asterisk. Right, orthogonal projection obtained from z stacks. Scale bar = 10 μm. (D) Multicolor images of live mouse embryonic stem cells (mESC) stained with WGA-488 and PK Mem 650. Mitochondrial was stained with PK Mito Red. Scale bar = 10 μm. (E) Multicolor image of a live rat sperm labeled with PK Mem 555 (200 nM), SPY505-DNA, and SPY650-tubulin. Scale bar = 10 μm. (F) LSCM of live hippocampal primary neurons stained with PK Mem 555 (20 nM, 10 min) without washing. Scale bar = 10 μm. (G) Multicolor image of a live adult rat cardiomyocyte labeled with PK Mem 555 (1 μM), Hoechst, PK Mito Deep Red, and Lyso-Tracker Green. Scale bar = 10 μm.

Next, KB cells were treated with PK Mem dyes (20 nM) for 5 min, resulting in a clean and bright fluorescence staining of the cellular membrane (Fig. S7). The strong fluorescence signal of PK Mem dyes enables 3D imaging and z-stack image reconstruction of live KB cells using laser scanning confocal microscopy (Fig. 2C and Fig. S8). PK Mem 555 exhibited distinct staining of tunneling nanotubes involved in intercellular communication (Fig. 2C and Video S1). We next challenged PK Mem dyes with various primary cells and for multicolor imaging. With the prolonged culture time, mESCs exhibited clonogenic growth with tightly arranged cells and an overall spherical shape. When co-stained with WGA-488, PK Mem dyes still exhibited relatively uniform and highly penetrable staining effects, while WGA-488 demonstrated limited diffusion and staining efficiency in densely packed mESCs (Fig. 2D and Fig. S9). A live rat sperm was labeled with PK Mem 555 (200 nM), SPY505-DNA, and SPY650-tubulin for three-color confocal imaging (Fig. 2E). The cell membrane of the sperm wrapped tightly around the axoneme of the flagellum. The tail of the sperm gradually becomes thinner, possibly due to the disappearance of the fibrous sheath^47^. This staining scheme of sperm can withstand formaldehyde fixation (Fig. S10). Alternatively, post-staining labeling of these three dyes can also achieve specific labeling (Fig. S11).

PK Mem dyes can effectively stain isolated hippocampal neurons at a low concentration of 20 nM (Fig. 2F and Fig. S12). Due to their remarkable photostability and minimal phototoxicity, PM dyes enable ultralong-term monitoring of axon growth of hippocampal neurons on DIV1. By capturing one image every two minutes with 0.3% power of a 561 nm laser line, it was feasible to monitor neurons stained with PK Mem 555 (20 nM) for up to 20.5 hours (Video S2). Membrane imaging was also a routine practice on fixed and permeabilized samples with multi-color immunofluorescence labeling. To test PK Mem dyes on fixed neurons, PK Mem 555 and Phalloidin-AF488 were applied to formaldehyde-fixed and Triton-permeabilized hippocampal neurons, giving the intricate structure of dendritic spines after fixation (Fig. S13A). In another demonstration, PK Mem 555, and a VGluT1 monoclonal antibody targeting glutamatergic axon terminals^48, 49^, were applied to a fixed and permeabilized hippocampal neuron. After further incubation with a secondary antibody for VGluT1 (Goat Anti-Rabbit IgG AF647), synapses that release glutamate can be identified on the plasma membrane (Fig. S13B). By incubating adult mouse cardiomyocytes with PK Mem 555, Hoechst, PK Mito Deep Red and Lyso-Tracker Green, a four-color image under live cell confocal microscopy was recorded (Fig. 2G). The strong fluorescence signal of PK Mem 555 enables the visualization of the cell membrane of adult mouse cardiomyocytes and their distinct cross-striated structure. Additionally, such four-color images can be recorded with adult mouse cardiomyocytes that were stained after fixation or fixed after staining (Fig. S14) with comparable image quality. For neonatal rat cardiomyocytes, PK Mem dyes could also be paired with Hoechst and Lyso-Tracker Green for multiplexed staining (Fig. S15). Overall, PK Mem dyes are bright and reliable stains for various primary cells, are compatible with fixation protocols, and pair well with other fluorescent markers.

### Imaging neurons in brain slices and in vivo

Due to the high brightness and fast diffusion speed of PK Mem dyes, we conducted further evaluations of their capabilities in brain slices and *in vivo* two-photon imaging. PK Mem 555 and Hoechst counterstain were applied to mouse brain slices (Fig. 3A) before the slice was imaged using a confocal microscope. High-quality images of cortical neurons and hippocampal neurons in mouse brain slices can be recorded (Fig. 3B and Video S3). Furthermore, by zooming in on the hippocampal region of the brain slices, the cell bodies and axons of neurons were clearly visible (Fig. 3B).

**Figure 3.**
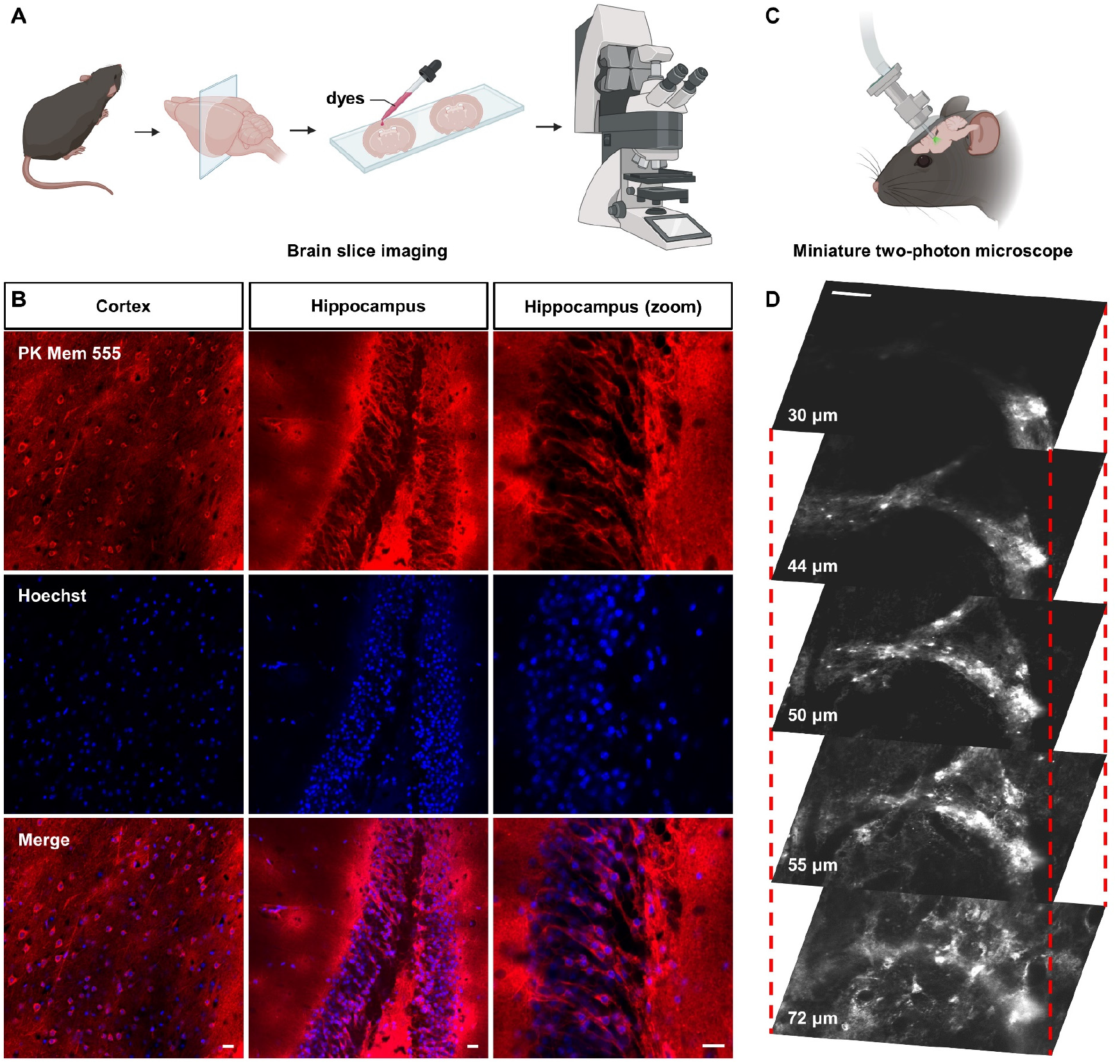
PK Mem 555 stains brain slices and a live mouse brain. (A) Scheme of neuronal imaging of mouse brain slices stained with dyes. (B) Confocal images of brain slices labeled with PK Mem 555 and Hoechst. Scale bar = 20 μm. (C) Scheme of the intravital two-photon imaging setup. (D) Two-photon images of PK Mem 555-labeled neurons in a mouse brain, see also Video. S3. Scale bar = 50 μm.

We further evaluated the potential of PK Mem dyes for *in vivo* imaging in mice using miniature two-photon microscopy with a single 920 nm fixed-wavelength fs-pulsed laser (Fig. 3C). The optical window provided an opportunity for two-photon imaging of awake mice, and the high signal-to-noise ratio of PK Mem 555 facilitated clear visualization of neuronal cell membranes (Fig. 3D). Moreover, blood vessels can be clearly identified in the superficial region of the mouse brain (Fig. S16). Both the brain slice and *in vivo* two-photon images demonstrated that PK Mem 555 preferentially highlights neurons for general fluorescence imaging.

### Ultralong-term dynamic imaging of cell membrane morphology, migration and beating with reduced phototoxicity

Phototoxicity-induced cell membrane blebbing and subsequent cell death are common issues during long-term recordings of live cells. We rigorously tested the phototoxicity of PK Mem dyes with imaging-based assays using head-to-head comparisons. HeLa cells labeled with PK Mem 555 or MemGlow 560 were imaged under time-lapse confocal microscopy with identical parameters. A representative movie demonstrated that HeLa cells labeled with PK Mem 555 began to bubble at frame 50, while those labeled with MemGlow 560 bubbled at frame 20 (Fig. 4A). The average cell bubbling time of PK Mem 555 (n=12) labeled cells was twice as long as that of MemGlow 560 (n=10) (Fig. 4B). Similar trends were recorded with PK Mem 590 and MemGlow 590, as well as PK Mem 650 and MemGlow 640 (Fig. S17). Overall, PK Mem dyes generally bear more than three-fold reduced phototoxicity in practical confocal imaging experiments, enabling long-term monitoring of dynamic morphological changes in cell membranes. Notably, such phototoxicity reduction is more pronounced in the live-cell imaging assay than that in ROS generation assay (Fig. 1F), suggesting a rich yet overlooked photochemistry and photobiology.

**Figure 4.**
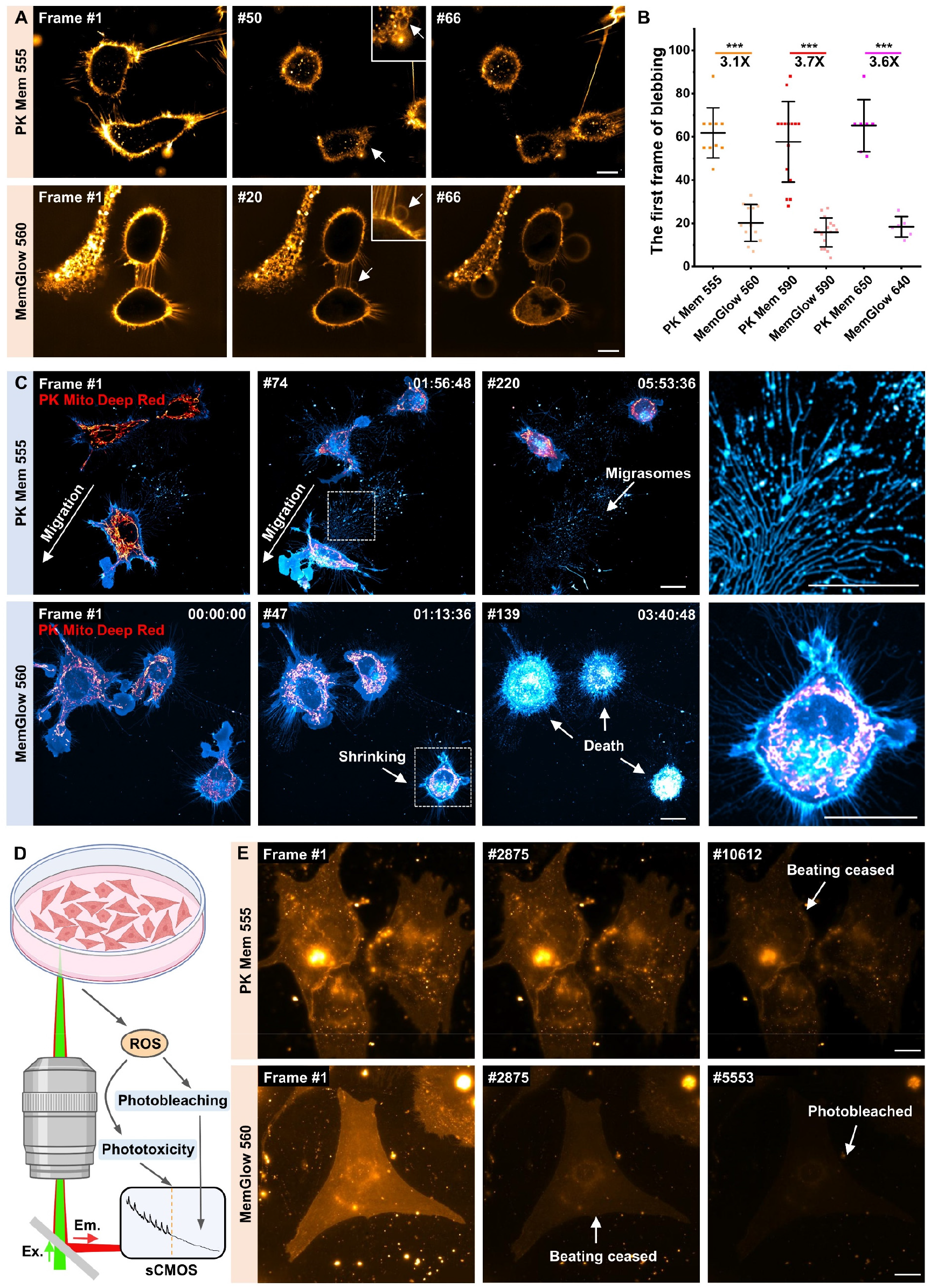
The low phototoxicity of PK Mem 555 enables long-term recording of cell migration and beating. (A) Long-term confocal recordings of HeLa cells labeled with MemGlow 560 and PK Mem 555. Blebbing events are highlighted with arrows, indicating photodamage. Scale bar = 10 μm. (B) Dot plots of the mean frame number in which the blebs emerge (N>5). Statistical significances were calculated with one-tailed Welch’s t-test (or two-tailed for symmetric stimulation), and p values were given for each comparison. (C) Long-term LSCM recordings of L929 cells labeled with MemGlow 560 and PK Mem 555. Mitochondria were stained with PK Mito Deep Red. The rightmost images show magnified views of the dashed white boxes in the second column, illustrating migrasomes and shrunken cells. Scale bar = 20 μm. (D) Schematic representation of long-term wide-field recordings of beating neonatal rat cardiomyocytes. Photobleaching diminishes fluorescence signal while phototoxicity can be characterized by the cease of beating. (E) Long-term wide-field recordings of neonatal rat cardiomyocytes labeled with MemGlow 560 and PK Mem 555. Scale bar = 10 μm.

Previous research has suggested that phototoxicity has a significant impact on cell migration speed^50^. In comparable video recordings, L929 cells labeled with PK Mem 555 and PK Mito Deep Red exhibited rapid migration, accompanied by the formation of retraction fibers and migrasomes throughout the nearly 6 h imaging duration (total 220 recorded frames). In contrast, cells labeled with MemGlow 560 and PK Mito Deep Red showed a compromised migration speed, accompanied by blebbing, mitochondrial deterioration, cell shrinkage, and death. (Fig. 4C and Video S4). The superior performance of PK Mem 555 labeling in time-lapse imaging, as evidenced by the maintenance of cell viability, highlights its privilege in time-lapse imaging and analysis.

We then switched to the recording of the beating process of cardiomyocytes which is orchestrated around plasma membranes. High-temporal-resolution imaging of neonatal rat cardiomyocytes is particularly challenging due to their increased sensitivity to phototoxicity^44^. Phototoxic damage can disrupt cardiomyocyte contractions (Fig. 4D). We employed wide-field microscopy to monitor the contractions of neonatal rat cardiomyocytes labeled with PK Mem 555 at high temporal resolution and high light intensity. Representative time-lapse videos revealed that neonatal rat cardiomyocytes, when labeled with PK Mem 555, ceased beating at frame 10612 while still retaining fluorescence on the membrane until frame 18001. In contrast, neonatal rat cardiomyocytes labeled with MemGlow 560 stopped beating around frame 2875 and completely lost fluorescence by frame 5553 (Fig. 4E and Video S5). These findings highlight the biocompatibility of PK Mem dyes for monitoring the physiological processes of sensitive and fragile primary cells. The live-cell imaging assays in this section establish PK Mem dyes as a gentle tool for investigating the intricate physiological dynamics of delicate primary cells.

### Super-resolution imaging of migrasomes and dendritic spines with STED microscopy

STED is a promising tool for studying cell membranes^51, 52^, particularly for imaging smaller membrane structures such as dendritic spines, retraction fibers, and membranous vesicles. Since STED microscopy typically requires higher light intensity than diffraction-limited approaches, phototoxicity and photobleaching can present a significant technical hurdle^53, 54^. Compared to Cy3, the presence of the naphthalene ring in Cy3.5 results in a more red-shifted emission spectrum without compromising photostability, making Cy3.5 an optically privileged chromophore for STED recording using a 775 nm depletion laser^42^. After conjugation with COT to enhance photostability and reduce phototoxicity, PK Mem 590 emerged as a promising probe for STED imaging of cell membrane structures, especially on live cells. Compared to conventional microscopy, STED imaging of the cell membrane of live L929 cells labeled with PK Mem 590 resulted in a clearly resolved membranous structure (Fig. S18).

We further studied migrasomes, the migration-dependent membrane-bound vesicular structures generated along retraction fibers in migrating cells, with diameters ranging from 500 nm to 3 μm^55^. By staining the cell membrane with PK Mem 590, the nucleus with Hoechst, the mitochondria with PK Mito Deep Red, and the lysosomes with Lyso-Tracker Green respectively, we obtained a four-color image (Fig. S19). PK Mem 590 effectively stained PM, along with the membranous extracellular vesicles such as migrasomes. Upon further magnification, the presence of an intriguing phenomenon of mitocytosis was uncovered, which is a migrasomes-mediated mitochondrial quality control process (Fig. S19)^56^. Following the proposed model where migrasomes are released along with the cell movements, migrasomes of myocardial fibroblasts can be imaged at the retracting end of the cell path. Using PK Mem 590 for STED imaging, migrasomes can be imaged at a sub-100 nm resolution, revealing their presence at the rear or crossroads of cell retraction fibers (Fig. 5A).

**Figure 5.**
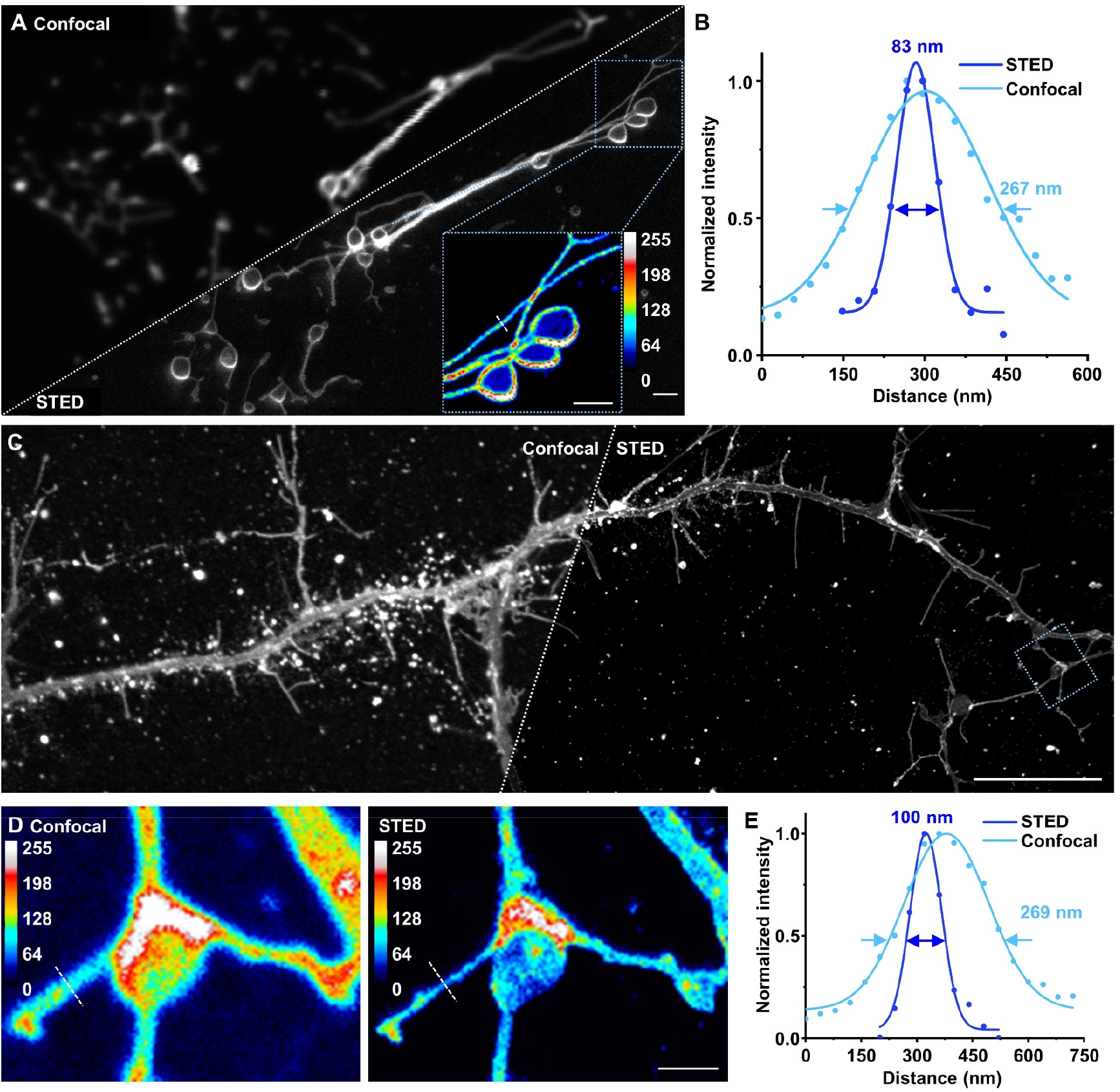
PK Mem 590 enables STED imaging on migrasomes and dendritic spines. (A) Confocal and STED images of migrasomes from live myocardial fibroblast. Scale bar = 1 μm. (B) Intensity profiles corresponding to the white dotted line of the STED and confocal images. (C) Confocal and STED images of axons and dendritic spines from a live hippocampal neuron. Scale bar = 10 μm. (D) Magnified view of the blue boxed area from Figure C. Scale bar = 1 μm. (E) Intensity profiles corresponding to the white dotted line of the STED and confocal images.

Moreover, we showcase the capability of PK Mem dyes to track endocytic vesicles. L1-CAM is a marker protein of endocytic vesicles that are related with neurite outgrowth, fasciculation, and migration^57, 58^. PK Mem 555 and L1-CAM monoclonal antibody were applied to live hippocampal neurons, followed by a fluorescent secondary antibody of L1-CAM. Imaging of live antibody uptake and internalization showcased that PK Mem 555 can track endocytic vesicles containing L1-CAM and monitor their internalization, highlighting its capability for monitoring endocytosis through internalization (Fig. S21).

Dendritic spines show activity-dependent and developmental regulation of their morphology. By employing STED microscopy, we could visualize axons and dendrites in primary hippocampal neurons labeled with PK Mem 590, particularly with fine details of dendritic spines (Fig. 5C). The implementation of STED microscopy yielded a remarkable enhancement in image resolution, surpassing approximately two-fold increase and providing a more detailed view of their morphology (Fig. 5D-E).

### Live-cell time-lapse super-resolution imaging of the growth cone and mitochondrial axonal transport

During neuronal development, neurons undergo polarization and swiftly extend their axons to form functional neural circuits. As highly dynamic structures located at the axon’s tip, growth cones direct axonal pathfinding toward their targets and facilitate axon elongation^59, 60^. Practically, PK Mem 590 enables time-lapse STED recordings of growth cones for ∼13 frames, which facilitated our understanding of the dynamic structural changes of the growth cone during neuronal migration processes (Fig. 6A and Video S6). Overall, the optical properties of PK Mem 590 render it a recommended probe for super-resolution imaging of PM using STED microscopy.

**Figure 6.**
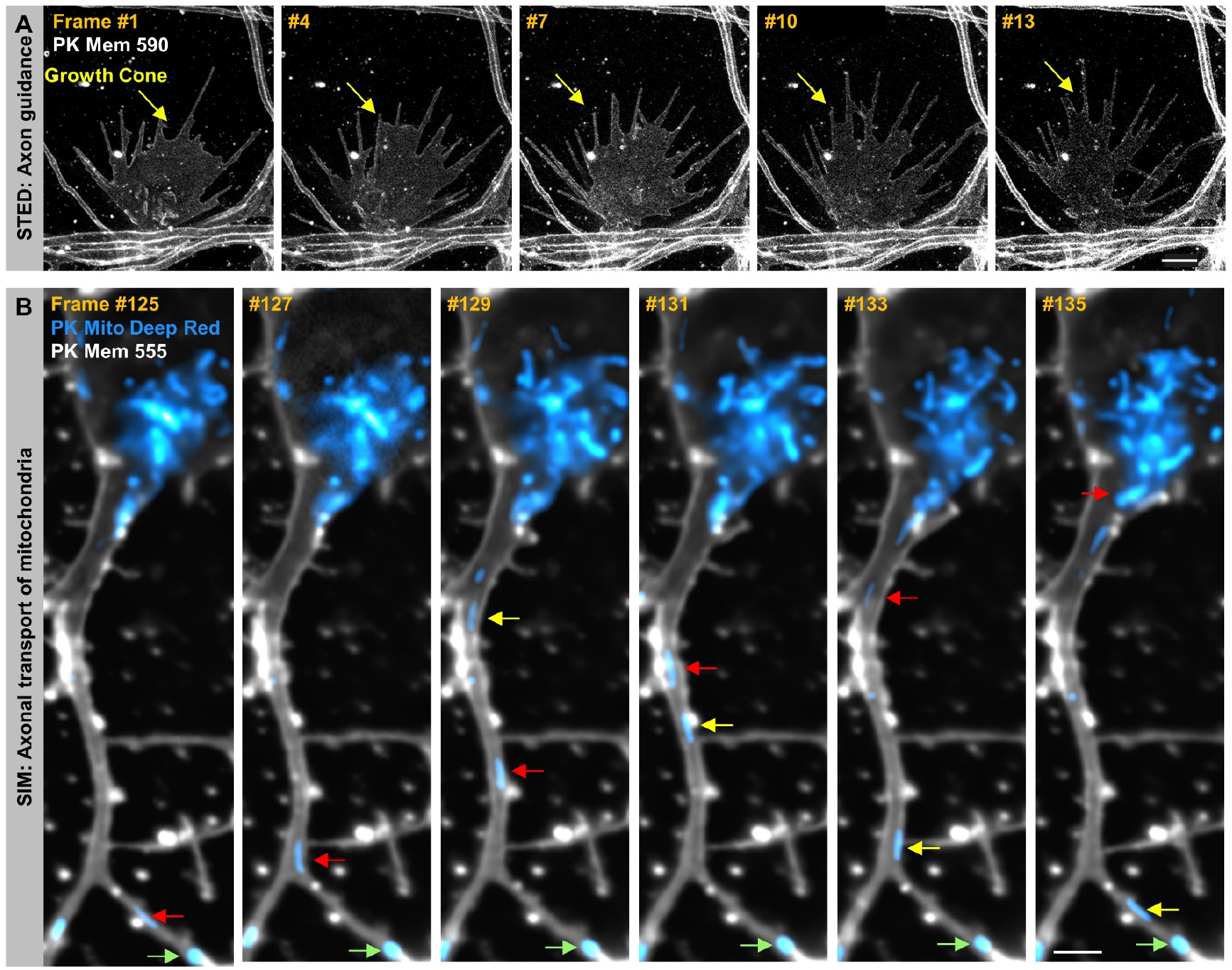
PK Mem dyes enable time-lapse super-resolution imaging of the growth cone and mitochondrial axonal transport. (A) Time-lapse STED images of the growth cone from the live hippocampal neuron stained with PK Mem 590. The arrow indicates the dynamics of the growth cone. Scale bar = 2 μm. (B) Time-lapse SIM images (0.5 Hz) of the axon from the live hippocampal neuron stained with PK Mem 555. Mitochondria were stained with PK Mito Deep Red. Anterograde transport (yellow arrow), retrograde transport (red arrow), and stationary/dynamic pause (green arrow) of mitochondria within the axon were indicated. Scale bar = 2 μm.

Axonal transport plays a vital role in maintaining neuronal health, in which mitochondria are important cargos. Anterograde transport ensures the transportation of healthy mitochondria from the soma to the axon terminals, while retrograde transport primarily facilitates the transfer of damaged mitochondria back to the soma for degradation and recycling^61, 62^. Structured illumination microscopy (SIM) is suitable for long-term, super-resolution imaging of live cells as it offers higher temporal resolution yet lower phototoxicity compared to STED microscopy^63-66^. During the time-lapse SIM imaging of neurons labeled with PK Mem 555 and PK Mito Deep Red, the mitochondrial bidirectional transports within neuronal axons were monitored for over 4 hours (7600 frames) (Fig. 6B, and Video S7). During the long-distance continuous transport of mitochondria, there were frequent occurrences of stationary/dynamic pauses and changes in direction. The elaborately tailored photophysical properties of PK Mem dyes, synergizing the enhanced resolution of SIM imaging, have allowed us to accurately distinguish the fluorescent signals of individual mitochondria within axons.

## Discussion

On the photophysical chemistry side of the fluorophores, conjugating with COT has been suggested to be a general strategy to reduce the phototoxicity and photobleaching of cyanine dyes. However, the effect of COT on photostability and phototoxicity seems to be largely influenced by the microenvironment. The *in vitro* singlet oxygen generation assay indicated that there was only approximately one-third reduction in the singlet oxygen quantum yields of PK Mem dyes compared with their non-COT counterparts (Table S2 & Fig. S3). However, PK Mem dyes showed more than a three-times decrease in phototoxicity in the live cell experiments of HeLa, L929 and cardiomyocyte (Fig. 4). The discrepancy between *in vitro* and cellular results may be due to differences in the microenvironment surrounding the dye. During the measurement of singlet oxygen quantum yield, the dye is present in a polar protic solvent, whereas in cellular experiments, the dye is surrounded by hydrophobic phospholipid molecules. We speculate that the excited-state energies of the dyes in different microenvironments are not the same. Moreover, in previous studies, the photostability of Cy3-COT conjugates appeared to be similar with the Cy3 counterpart without COT^34, 39^. However, COT conjugated to the membrane-localized Cy3 (as in PK Mem 555) enhanced the photostability by 42% (Table S2). These data collectively suggested that the photochemistry of excited-states is sensitive to the microenvironment, which should be further studied using both rigorous in vitro assays and validated on biological systems.

From a cell biology perspective, PK Mem dyes offer bright, specific, fluorogenic, yet gentle imaging of cell membranes. One limitation of PK Mem dyes is that, as amphiphilic organic molecules which have affinities to albumins, the compatibility of PK Mem with FBS is still limited. Adding FBS generally causes a loss of specific fluorescence on the cell membrane. For neurons whose maintenance does not depend on FBS, this issue would be less of a concern. Rather, the low concentration and minimal phototoxicity of PK Mem dyes allow for more physiological monitoring of neuronal activities. In this sense, PK Mem dyes and WGA-dye conjugates supplement and supplant each other as ideal PM stains for different applications.

In conclusion, we demonstrate PK Mem dyes as a robust toolkit for general imaging of plasma membranes. These dyes are compatible with various cancer cells or primary cells, enabling live-cell and fixed-cell imaging. Particularly, their reduced phototoxicity and high photostability make them ideal for long-term monitoring of physiological activities on the cell membrane, such as cell migration and cardiomyocyte contraction. PK Mem dyes offer bright neuronal membrane staining at concentrations as low as 20 nM and enable super-resolution imaging of dendritic spines morphology and long-term super-resolution imaging of growth cones extension and mitochondrial transport in axons. We anticipate that PK Mem dyes will become common reagents for membrane imaging in the time-lapse super-resolution era.

## Acknowledgments

This project was supported by funds from National Key R&D Program of China (2021YFF0502904 to Z.C.), and Beijing Municipal Science & Technology Commission (Project: Z221100003422013 to Z.C.). We thank Prof. Heping Cheng and Prof. Shiqiang Wang for the guidance on cardiomyocyte isolation, and the NMR facility of the National Center for Protein Sciences at Peking University for assistance with data acquisition. We also thank PKU-Nanjing Joint Institute of Translational Medicine, Nanjing 211800, China. -- Brain Observatory for providing miniature two-photon microscopy and the Nanjing Brain Observatory for its assistance in miniaturized two-photon surgery, data collection, and data analysis. Moreover, we thank Airy Technologies Co. Ltd., for providing with Airy Polar-SIM system for SIM imaging. Figure. 1C, 2A, 3A, 3C and 4D were created with BioRender.com.

## Author Contributions

J.L., and Y.L. performed all experiments and data analyses. Z.C. conceived and supervised the project. J.L., Y.L. and Z.C. wrote the paper with input from all authors. Y.F. and P.X. provided experiential guidance for SIM imaging experiments. S.L. provided guidance for the culture of neurons. L.D. provided mouse sperms. L.H. provided mouse embryonic stem cells.

---

**Scheme 1.**
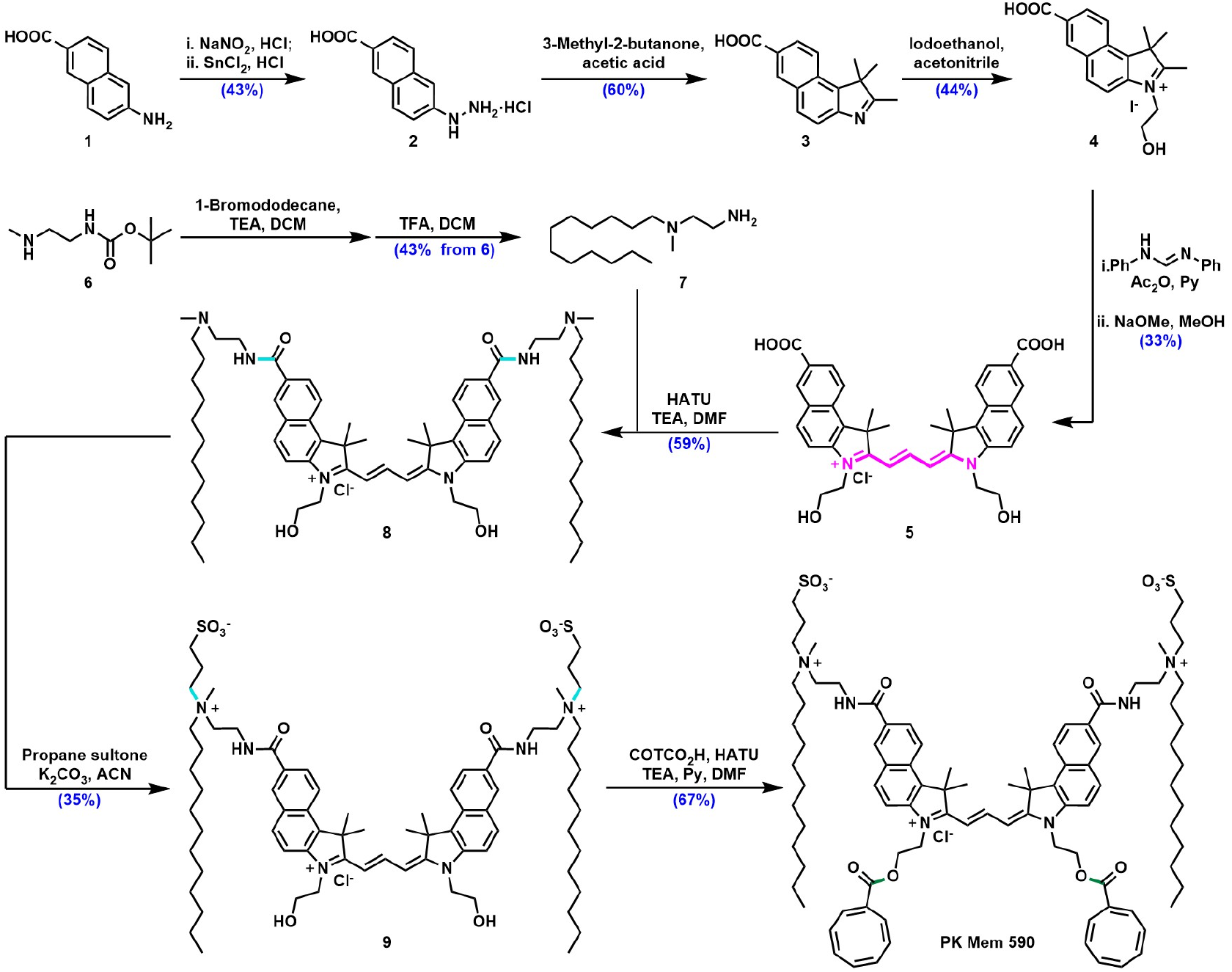
Synthetic route of PK Mem 590 via a modular derivatization of cyanine dyes with amphiphilic linkers and cyclooctatetraenes.

