## Supplemental Information for "A gentle palette of plasma membrane dyes"

**Supplementary Videos**

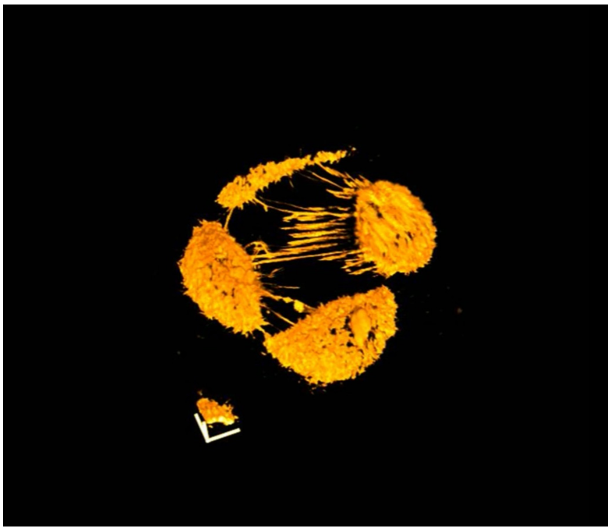

Video S1. 3D reconstruction of live KB cells stained with PK Mem 555.

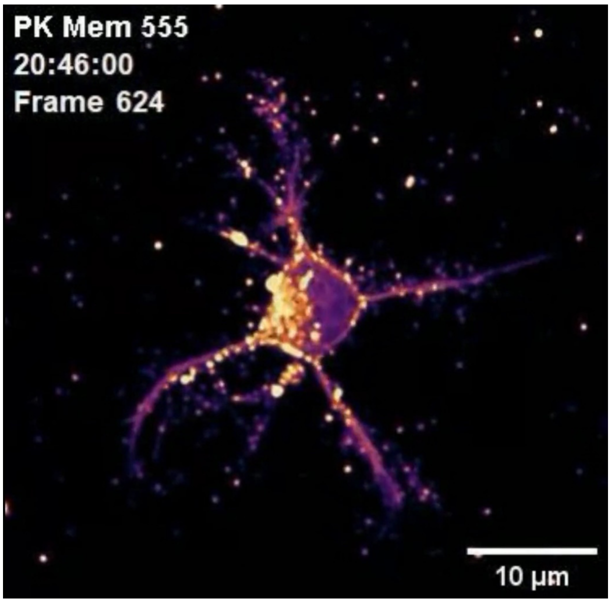

Video S2. Long-term confocal imaging of a neuron stained with PK Mem 555 (20.6 h, 624 frames). Scale bar = 10 µm.

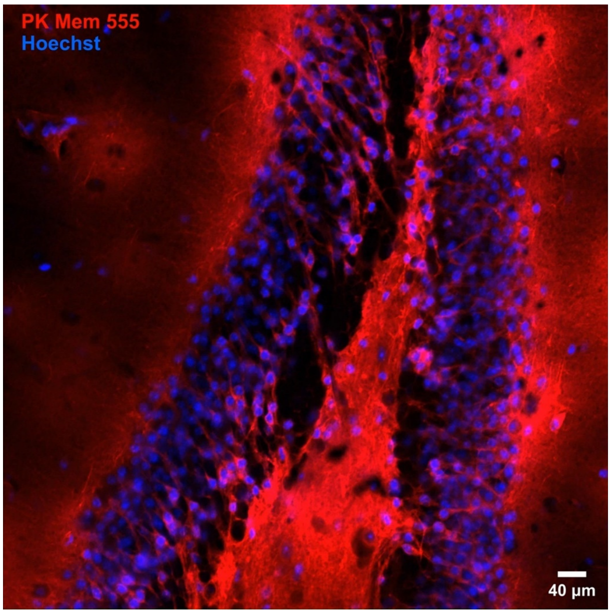

Video S3. Confocal z-stack of brain slices stained with PK Mem 555 and Hoechst. Scale bar = 40 µm. Related to Figure 3.

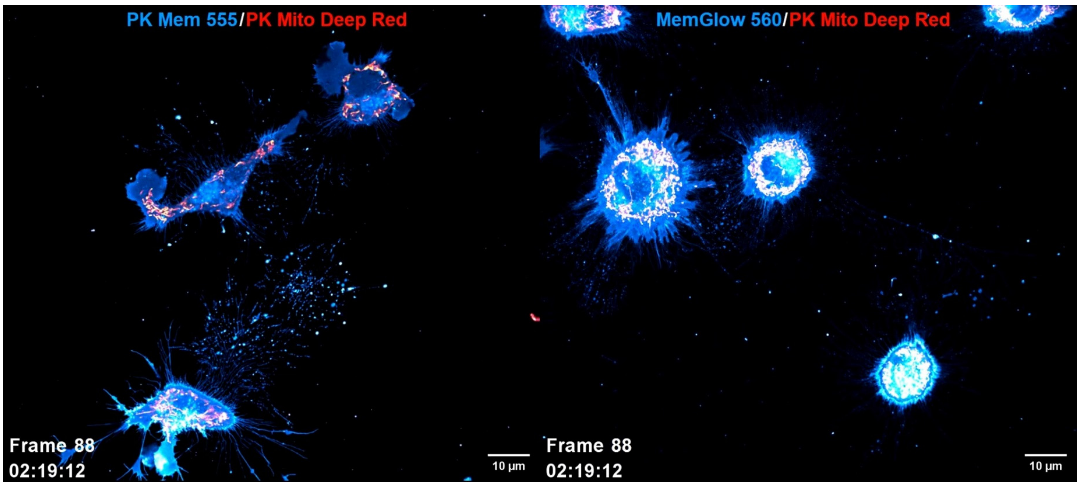

Video S4. Long-term confocal imaging of L929 cells migration over 6 h (220 frames). Cell membranes were stained with PK Mem 555 (left) or MemGlow 560 (right). Mitochondria were stained with PK Mito Deep Red. Scale bars = 10 µm. Related to Figure 4.

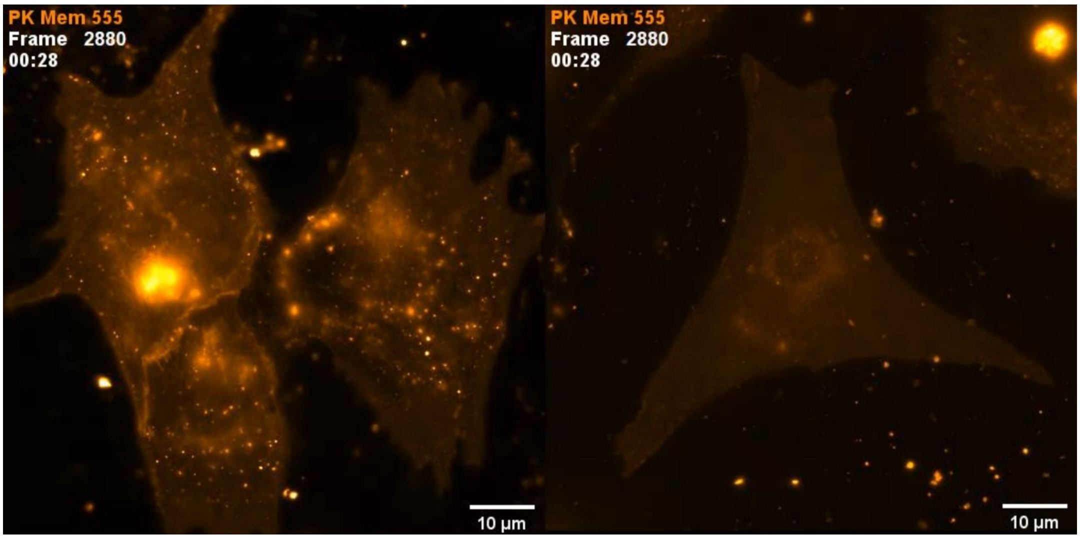

Video S5. Time-lapse wide-field imaging (3 min, 100 Hz, 18000 frames) of beating neonatal rat cardiomyocytes stained with PK Mem 555 (left) or MemGlow 560 (right). Scale bars = 10 µm.

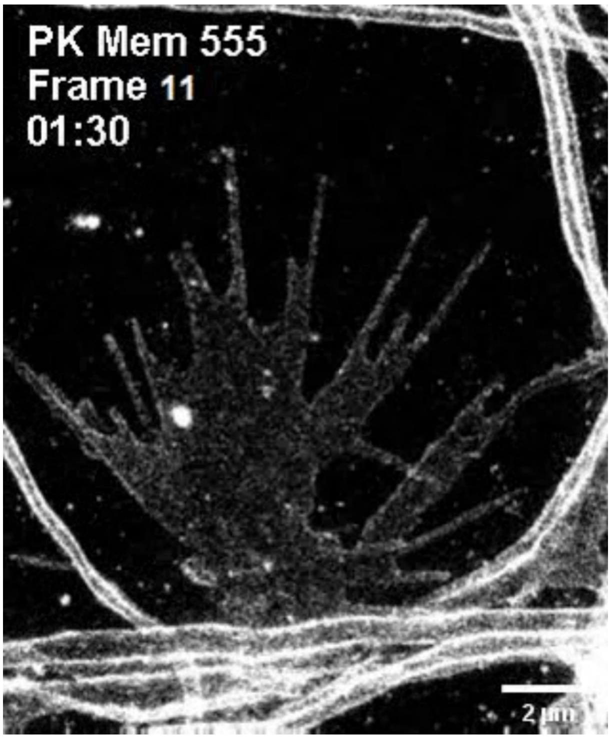

Video S6. Time-lapse STED recording (2.1 min, 15 frames) of the growth cone from a live hippocampal neuron stained with PK Mem 590. Scale bar = 2 µm.

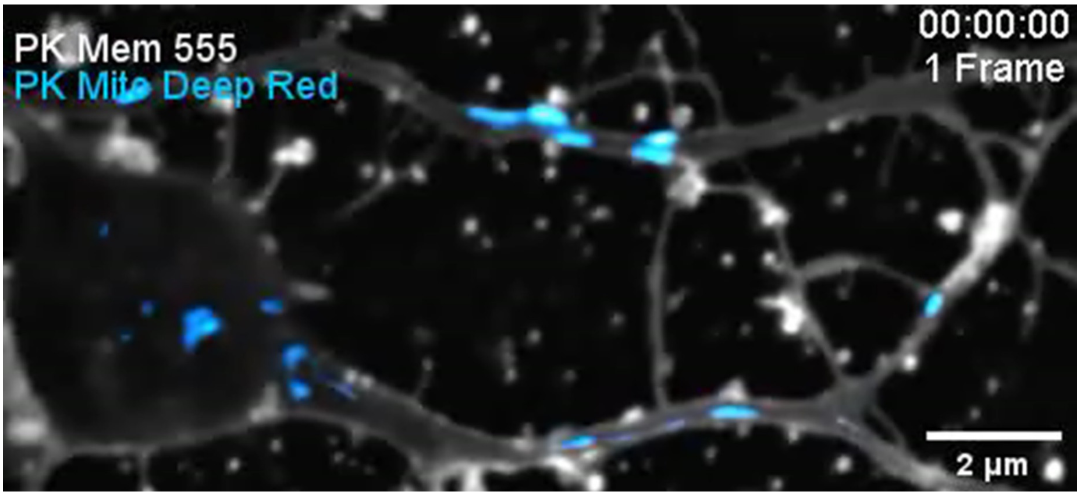

Video S7. Time-lapse SIM imaging (4.2 h, 7600 frames) of an axon from a live hippocampal neuron. The cell membrane was stained with PK Mem 555. Mitochondria were stained with PK Mito Deep Red. Scale bar = 2 μm.

### Supplementary Tables

Table S1. Spectral properties of PK Mem dyes in various solvents.

| Solvent | | PK Mem 555 | PK Mem 590 | PK Mem 650 |
|---|---|---|---|---|
| **MeOH** | $\lambda_{abs}$ (nm) | 560 | 596 | 654 |
| | $\lambda_{emi}$ (nm) | 579 | 612 | 677 |
| | $\Phi_f$ | 0.22 | 0.27 | 0.40 |
| **PBS** | $\lambda_{abs}$ (nm) | 594 | 608 | 626 |
| | $\lambda_{emi}$ (nm) | 607 | 626 | 653 |
| | $\Phi_f$ | 0.01 | 0.01 | 0.002 |
| **DOPC SUVs** | $\lambda_{abs}$ (nm) | 564 | 598 | 662 |
| | $\lambda_{emi}$ (nm) | 579 | 615 | 681 |
| | $\Phi_f$ | 0.32 | 0.43 | 0.33 |

The absorption and emission spectra of PK Mem dyes in different solutions were measured using a Duetta fluorescence and absorbance spectrometer (Horiba, Kyoto, Japan) in a 1 cm square quartz cuvette. Maximum emission wavelengths were measured with 510 nm excitation light for PK Mem 555, 540 nm extinction light for PK Mem 590 and 580 nm excitation light for PK Mem 650 For vesicles, PK Mem dyes were added into a PBS solution of 200 μM DOPC, and the measurements were done after 30 minutes. Quantum yields were measured with Rhodamine B in MeOH as reference (Φ=0.65), The fluorescence quantum yields, $\Phi_f$ (sample), were calculated according to the equation as follows:

$$\frac{\Phi_{f,sample}}{\Phi_{f,ref}} = \frac{OD_{ref} \cdot I_{sample}}{OD_{sample} \cdot I_{ref}}$$

$\Phi_f$: quantum yield of fluorescence;

I: integrated emission intensity;

OD: optical density at the excitation wavelength;

Table S2. Photobleaching half-life and ROS generation of MemGlow, Az-Cy, and PK Mem dyes

| **Fluorophore** | **Excitation (nm)** | **Illumination intensity ($\mu W \cdot cm^{-2}$)** | **Photobleaching half-life $t_{1/2}$ (s)** [a] | **$\Phi$ ($^1O_2$)** |
|---|---|---|---|---|
| **MemGlow 560** | 561 | 2.5 | 39.6 | n.d. |
| **Az-Cy3** | 561 | 2.5 | 64.9 | 0.0050 |
| **PK Mem 555** | 561 | 2.5 | 92.6 | 0.0039 |
| **MemGlow 590** | 561 | 16.6 | 9.3 | n.d. |
| **Az-Cy3.5** | 561 | 16.6 | 14.3 | 0.0089 |
| **PK Mem 590** | 561 | 16.6 | 38.3 | 0.0069 |
| **MemGlow 640** | 640 | 38.6 | 19.2 | n.d. |
| **Az-Cy5** | 640 | 38.6 | 18.5 | 0.0120 |
| **PK Mem 650** | 640 | 38.6 | 49.7 | 0.0083 |

[a] measured in formaldehyde-fixed HeLa cells labeled with MemGlow and PK Mem dyes to reduce motion and depolarization induced artifacts. n.d., not determined. "±" represents s.d.

Table S3. List of reagents used in this study

| Reagent | Vendor | Catalog Number |
|---|---|---|
| Dulbecco's Modified Eagle's medium (DMEM) | Gibco | C11995500BT |
| Minimum Essential Medium (MEM) | Gibco | C11095500BT |
| Fetal Bovine Serum (FBS) | VisTech™ | SE200-ES |
| Trypsin-EDTA (0.25%) | Gibco | 25200056 |
| Neurobasal™ Medium | Gibco | 21103049 |
| B-27™ Supplement | Gibco | 17504044 |
| GlutaMAX™ Supplement | Gibco | 35050061 |
| 5-Bromo-2′-deoxyuridine (BrdU) | Sigma | B5002 |
| Collagenase Type II | Worthington | WBC-LS004202 |
| Pancreatin | Sigma | P3292 |
| Penicillin-streptomycin | ThermoFisher Scientific | 15140122 |
| Tyrode's Salts Solution | Yuanye | R20266 |
| Dulbecco's Phosphate-Buffered Saline (DPBS) | Gibco | C14190500BT |
| Hank's Balanced Salt Solution (HBSS) | Gibco | C14175500BT |
| Matrigel® Matrix | Corning | 356234 |
| poly-D-lysine | Sigma | P7280-5X5M |
| Laminin Mouse Protein | Gibco | 23017015 |
| Opti-MEM™ Medium | Gibco | 31985062 |
| Formaldehyde solution | Yuanye | R20497 |
| WGA-iFluor ™ 488 | AAT Bioquest | 25530 |
| Hoechst 33342 | ThermoFisher Scientific | H1399 |
| SPY505-DNA | Spirochrome | SC101 |
| SPY650-tubulin | Spirochrome | SC503 |
| MemGlow 560 (MEMBRIGHT Family Probe) | Cytoskeleton | MG02-02 |
| MemGlow 590 (MEMBRIGHT Family Probe) | Cytoskeleton | MG03-02 |
| MemGlow 640 (MEMBRIGHT Family Probe) | Cytoskeleton | MG04-02 |
| Lyso-Tracker Green | Beyotime | C1047S |
| PK Mito Red | Genvivo | PKMR-1 |
| PK Mito Deep Red | Genvivo | PKMDR-1 |
| Anti-L1CAM antibody | Abcam | ab272733 |
| Anti-VGluT1 antibody | Abcam | ab227805 |
| Phalloidin-488 | AAT Bioquest | 23115 |
| Donkey anti-Rabbit IgG (H+L) Highly Cross-Adsorbed Secondary Antibody, Alexa Fluor™ 488 | ThermoFisher Scientific | A-21206 |
| Goat Anti-Rabbit IgG H&L (Alexa Fluor® 647) | Abcam | ab150079 |

### Supplementary Figures

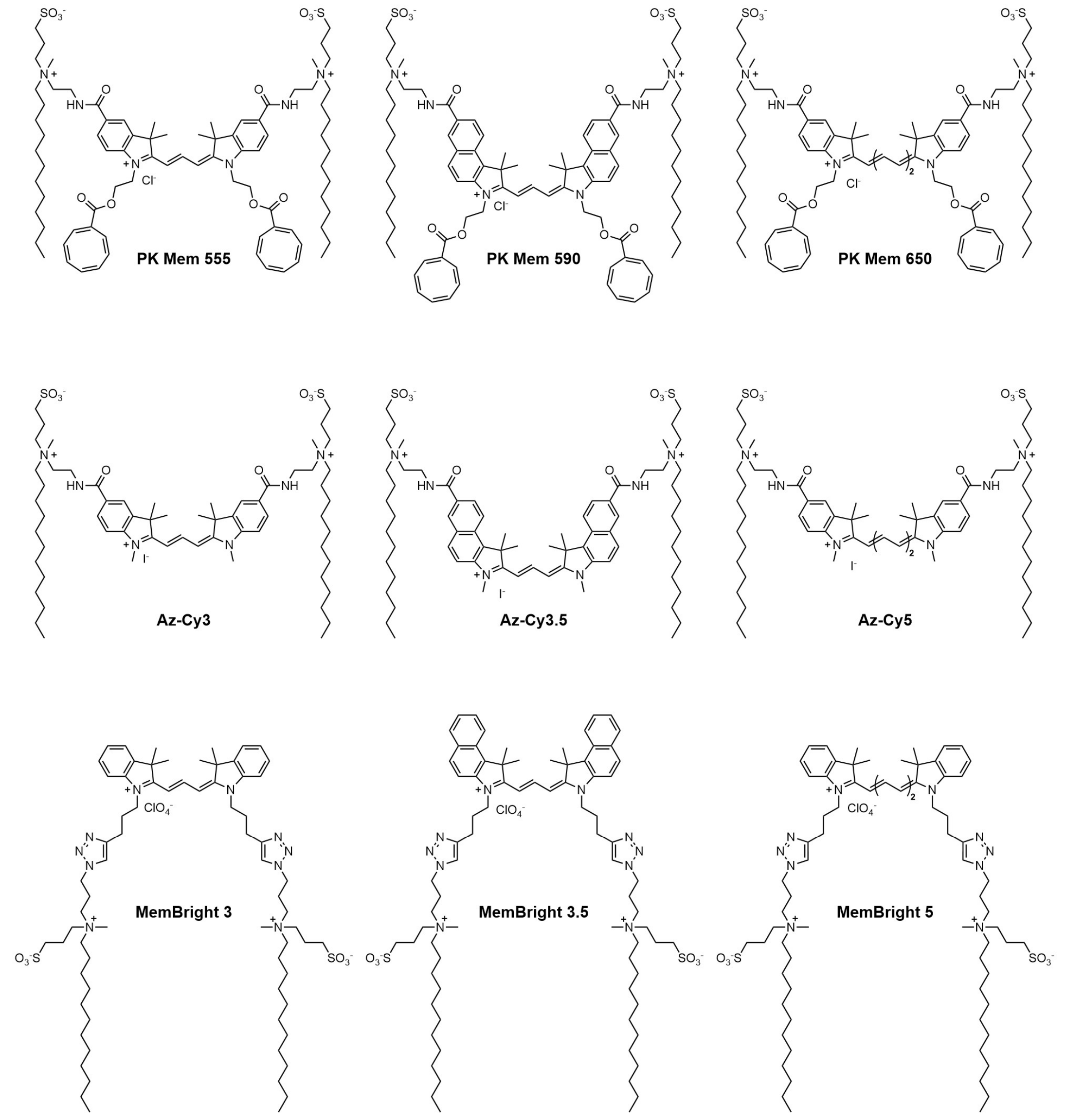

Figure S1. Chemical structures of PK Mem, Az-Cy and MemBright dyes.

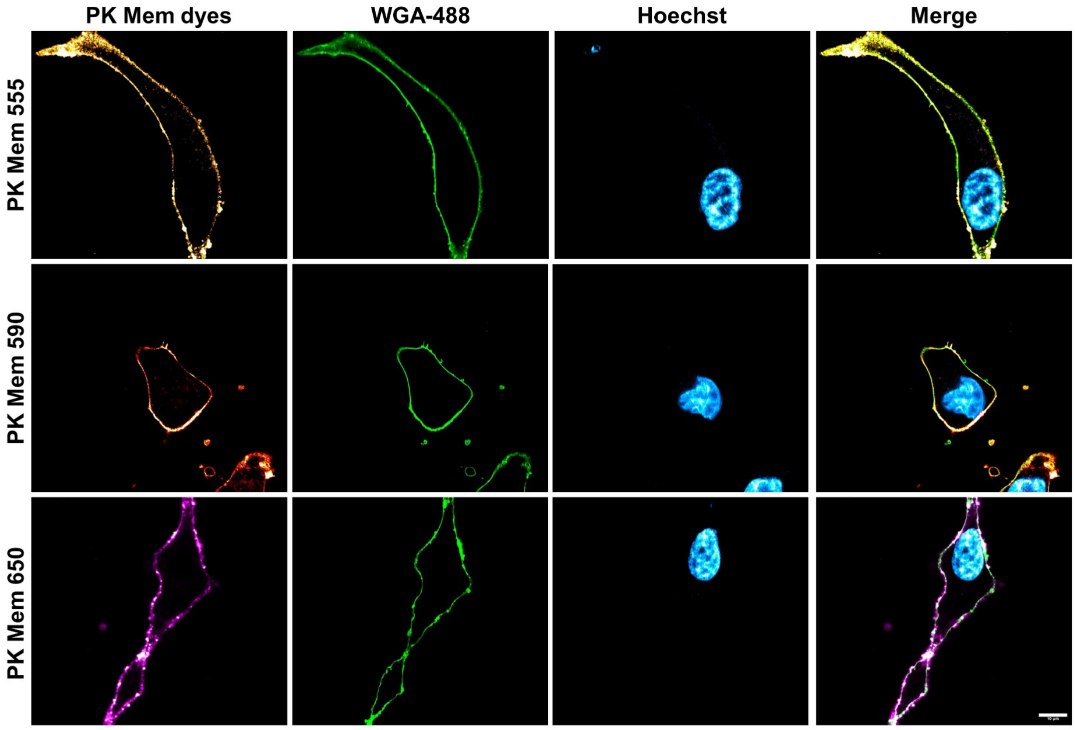

Figure S2. Laser scanning confocal microscopy of HeLa cells labeled with PK Mem dyes (20 nM) for 5 min without washing. WGA-488 (5 µg/mL) was used as a co-staining marker. The nucleus was stained with Hoechst (5 µg/mL). Scale bar = 10 µm.

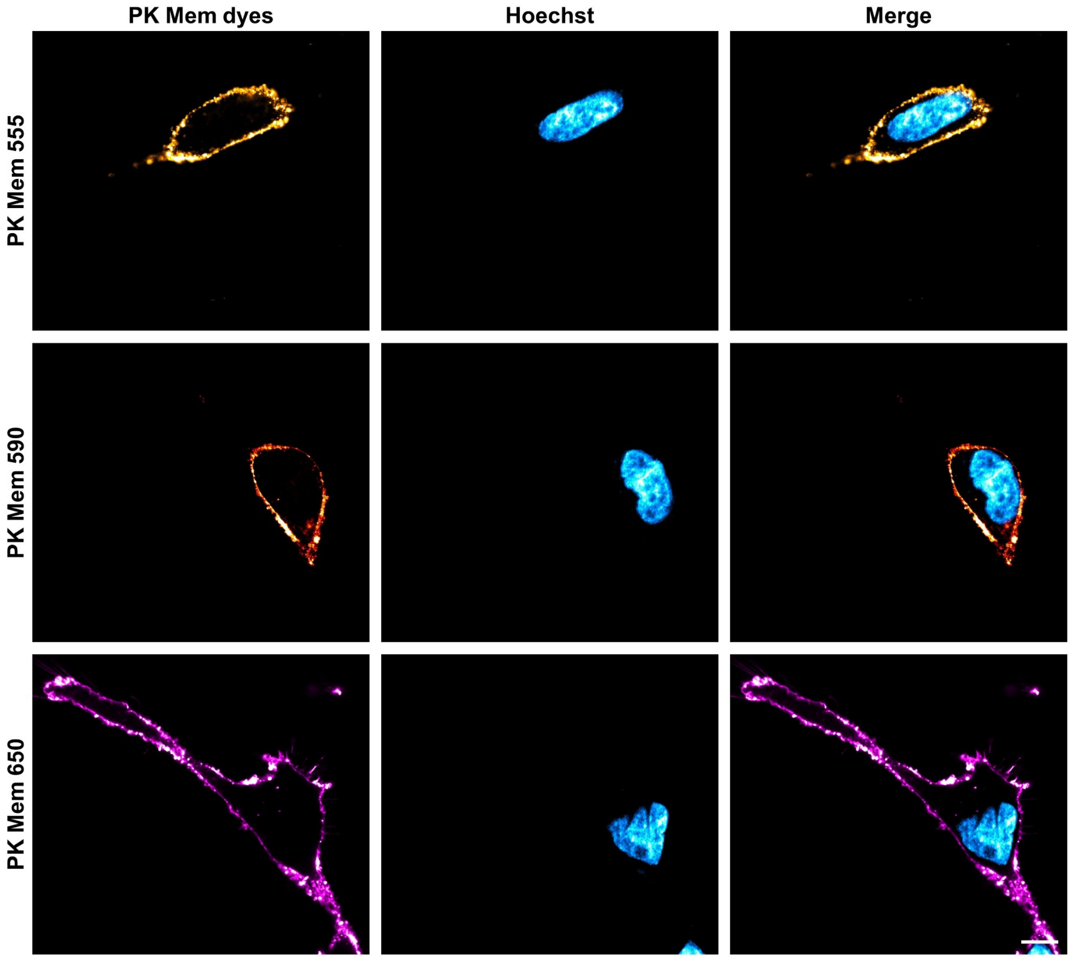

Figure S3. Laser scanning confocal microscopy of HeLa cells labeled with PK Mem dyes (20 nM) at 37° C for 120 min without washing. The nucleus was stained with Hoechst (5 μg/mL). Scale bar = 10 μm.

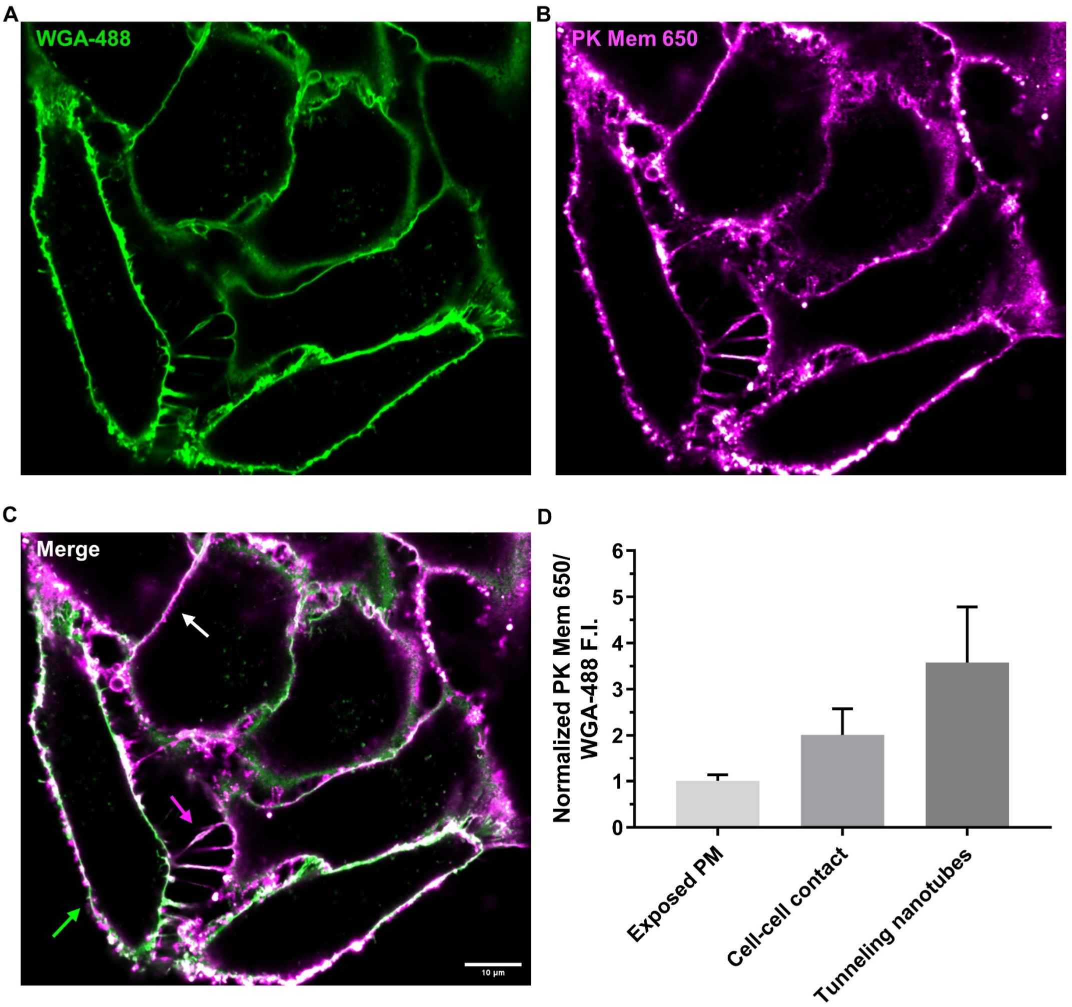

Figure S4. Laser scanning confocal microscopy of confluent HeLa cells labeled by (A) WGA-488 (5 μg/mL), (B) PK Mem 650 (20 nM), (C) merge of the green and red channels. Scale bar = 10 μm. The green arrow indicates a region of exposed plasma membrane, the white arrow indicates a cell-cell contact region, and the magenta arrow indicates a tunneling nanotube. (D) Normalized PK Mem 650 fluorescence intensity / WGA-488 fluorescence intensity ratios at exposed plasma membranes, cell-cell contact, and tunneling nanotubes. Each measure was performed on 10 different regions of plasma membranes.

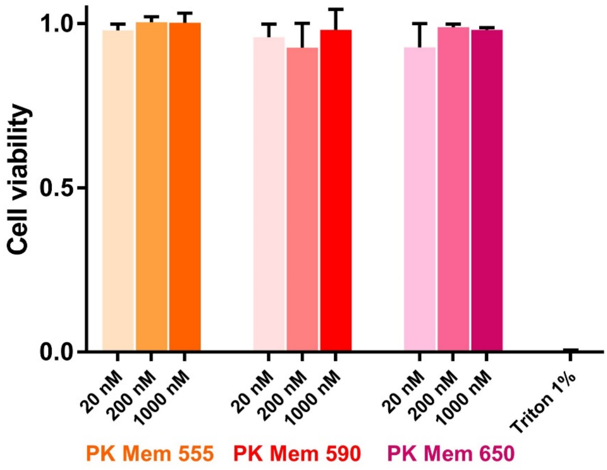

Figure S5. CCK-8 assay measuring HeLa cell viability after exposing to different concentrations of PK Mem dyes for 1 h. Triton (1%) was used as a positive control of cytotoxicity

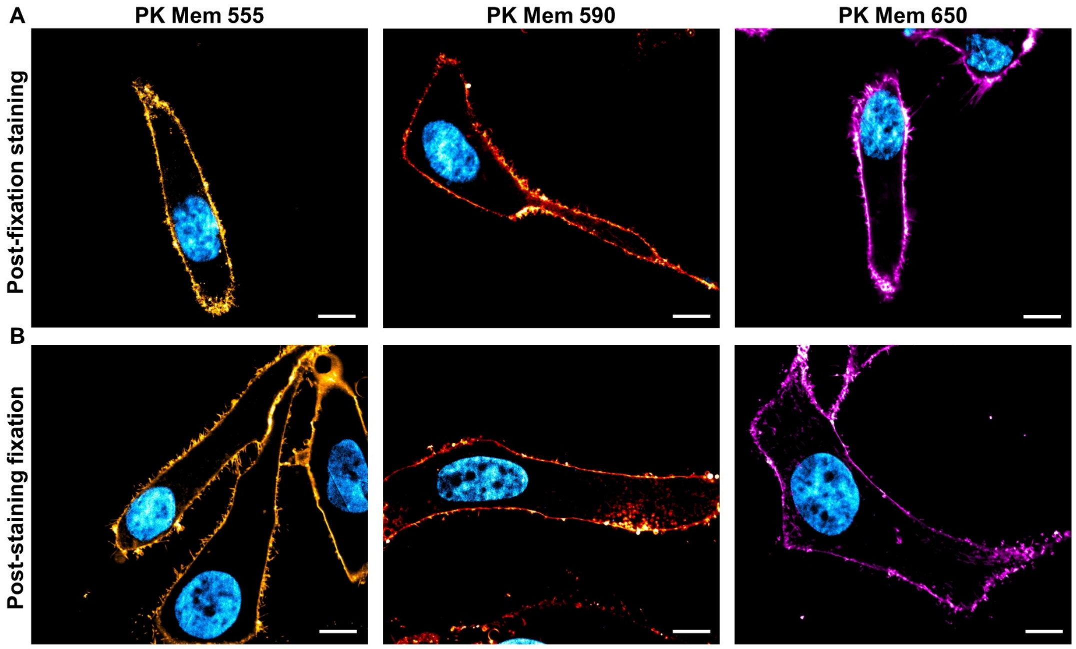

Figure S6. Laser scanning confocal microscopy of HeLa cells labeled with PK Mem dyes (200 nM). (A) Laser scanning confocal microscopy of HeLa cells staining with PK Mem dyes (200 nM) after formaldehyde (PFA) fixation. The nucleus was stained with Hoechst (5 µg/mL). Scale bar = 10 µm. (B) Laser scanning confocal microscopy of HeLa cells fixed with PFA after incubation with PK Mem dyes (20 nM). The nucleus was stained with Hoechst (5 µg/mL). Scale bars = 10 µm.

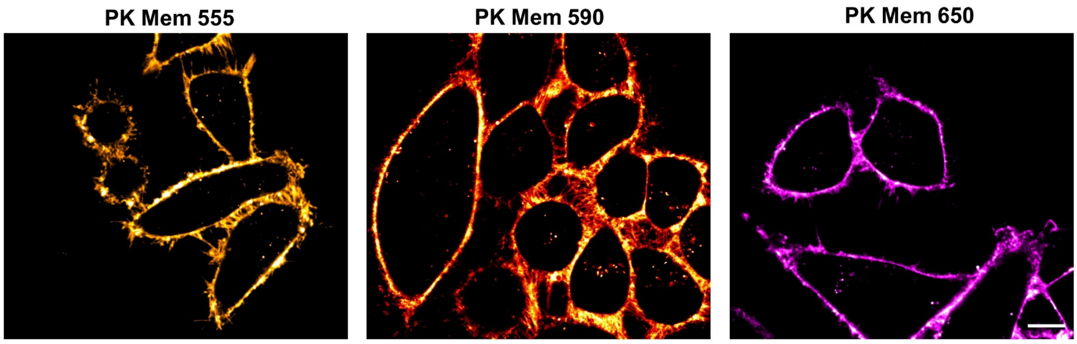

Figure S7. Laser scanning confocal microscopy of live KB cells 5 min after the addition of PK Mem dyes (20 nM) without washing. Scale bar = 10 µm.

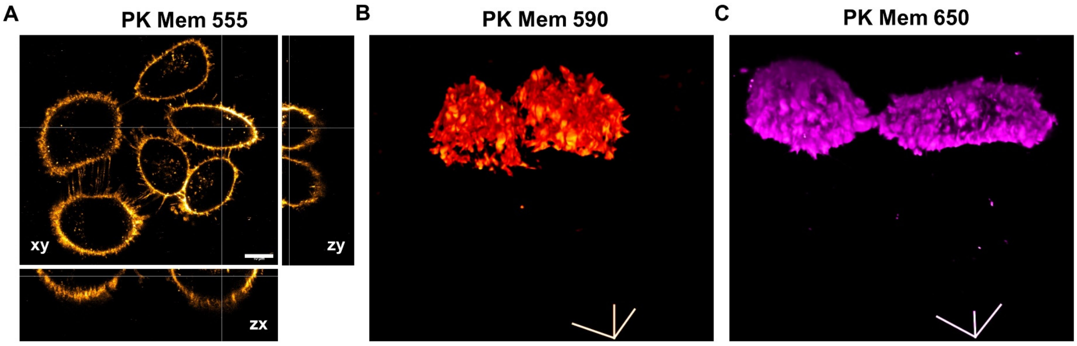

Figure S8. Laser scanning confocal microscopy of KB cells labeled with PK Mem dyes (20 nM) at 5 min without washing. (A) Orthogonal projection obtained from z stacks of PK Mem 555. Scale bar = 10 µm. (B) 3D reconstruction of live KB cells stained with PK Mem 590. Scale bar = 10 µm. (C) 3D reconstruction of live KB cells stained with PK Mem 650. Scale bar = 10 µm.

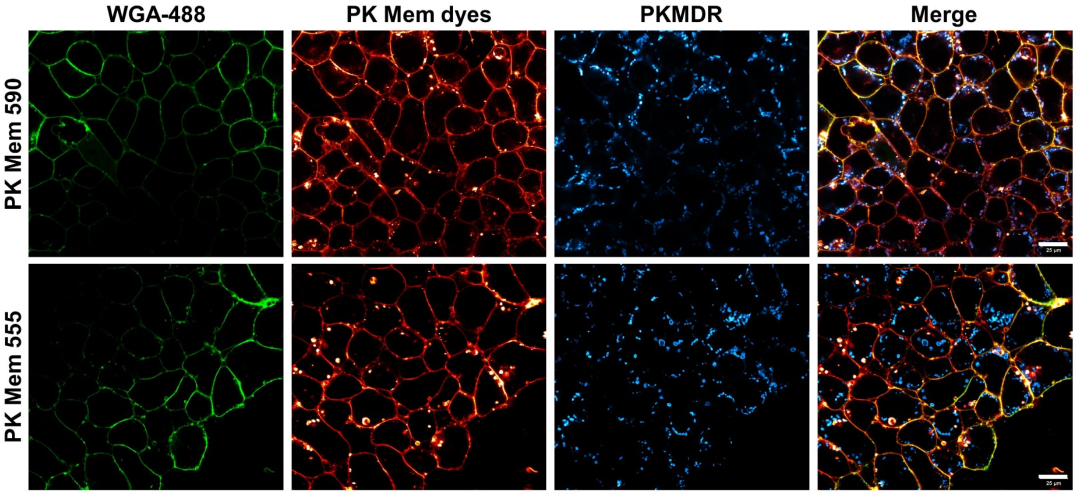

Figure S9. Laser scanning confocal microscopy of Mouse Embryonic Stem Cells (mESC) labeled with PK Mem dyes (200 nM) at 10 min without washing. WGA-488 (5 µg/mL) was used as a co-staining marker. Mitochondria were stained with PK Mito Deep Red. Scale bar = 25 µm.

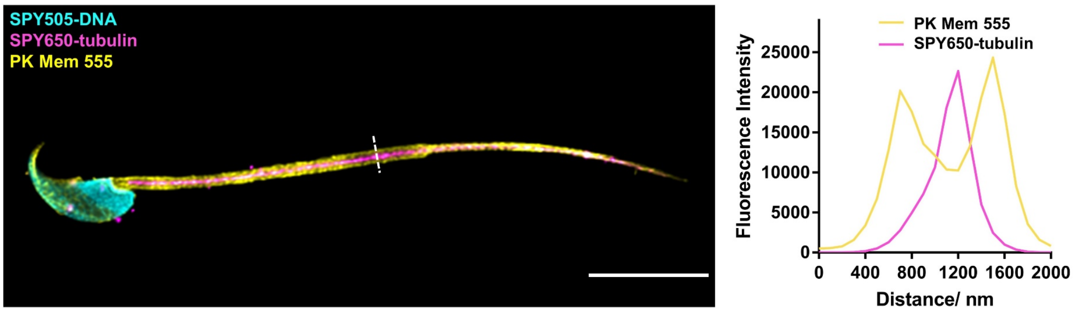

Figure S10. Laser scanning confocal microscopy images of a fixed sperm (left). The sample was stained with PK Mem 555 (200 nM), SPY505-DNA and SPY650-tubulin. Fixation was performed after staining. Scale bar = 10 μm. Fluorescence intensity line profiles are plotted (right).

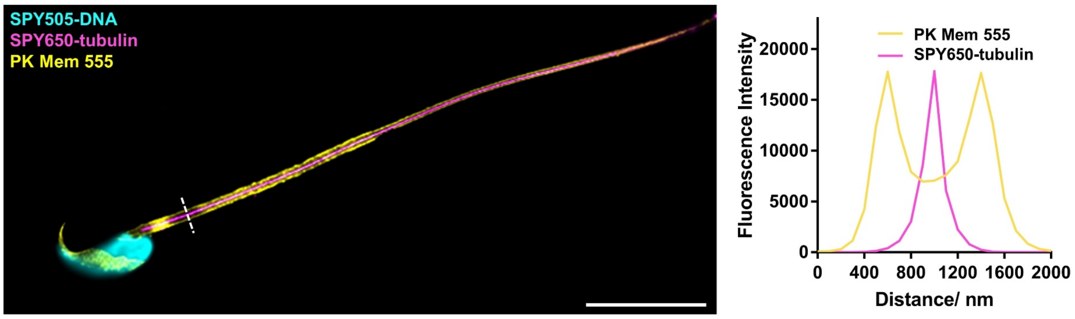

Figure S11. Laser scanning confocal microscopy images of a fixed sperm (left). The sample was fixed for 10 minutes at RT in 4% formaldehyde. After fixation, the sample was stained with PK Mem 555 (200 nM), SPY505-DNA and SPY650-tubulin. Scale bar = 10 μm. Fluorescence intensity line profiles are plotted (right).

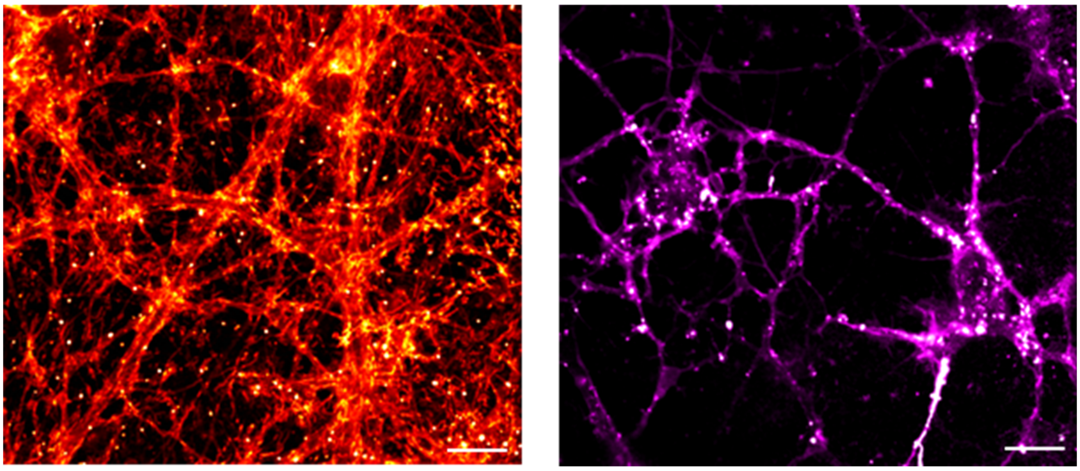

Figure S12. Laser scanning confocal microscopy of live hippocampal primary neurons stained with PK Mem 590 (20 nM, left) or PK Mem 650 (20 nM, right) 10 min without washing. Scale bar = 10 μm.

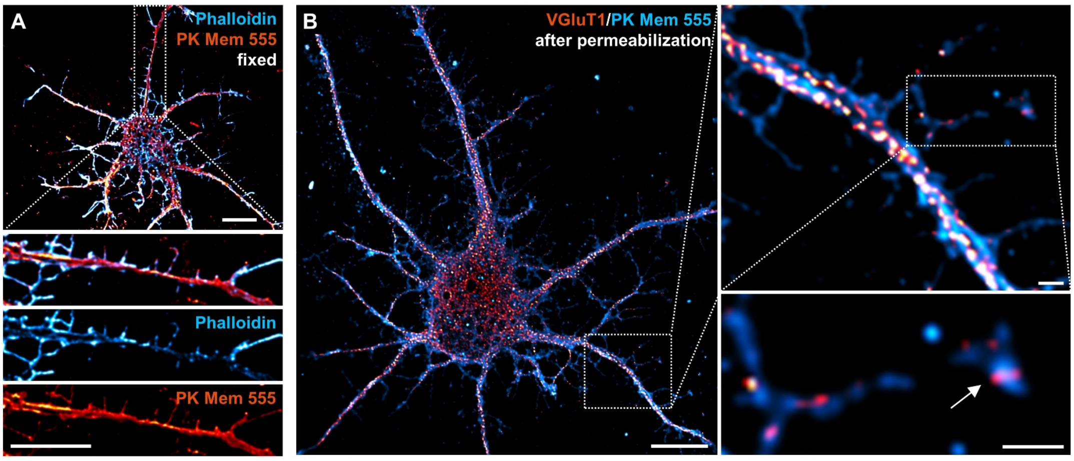

Figure S13. Multicolor imaging using PK Mem 555 and various labels. (A) LSCM images of fixed primary hippocampal neurons incubated with PK Mem 555 (red) and Phalloidin AF488 (cyan), highlighting dendritic spine head and neck. The white boxed area is further magnified (bottom). Scale bars = 10 μm. (B) LSCM images of a primary hippocampal neuron fixed, permeabilized, and stained with PK Mem 555 (cyan) and VGluT1 monoclonal antibody (red, visualized with Goat Anti-Rabbit IgG AF647). Scale bar: 10 μm. The white boxed area is further magnified (right). The white arrow indicated a VGluT1 positive synapse. Scale bar = 1 μm.

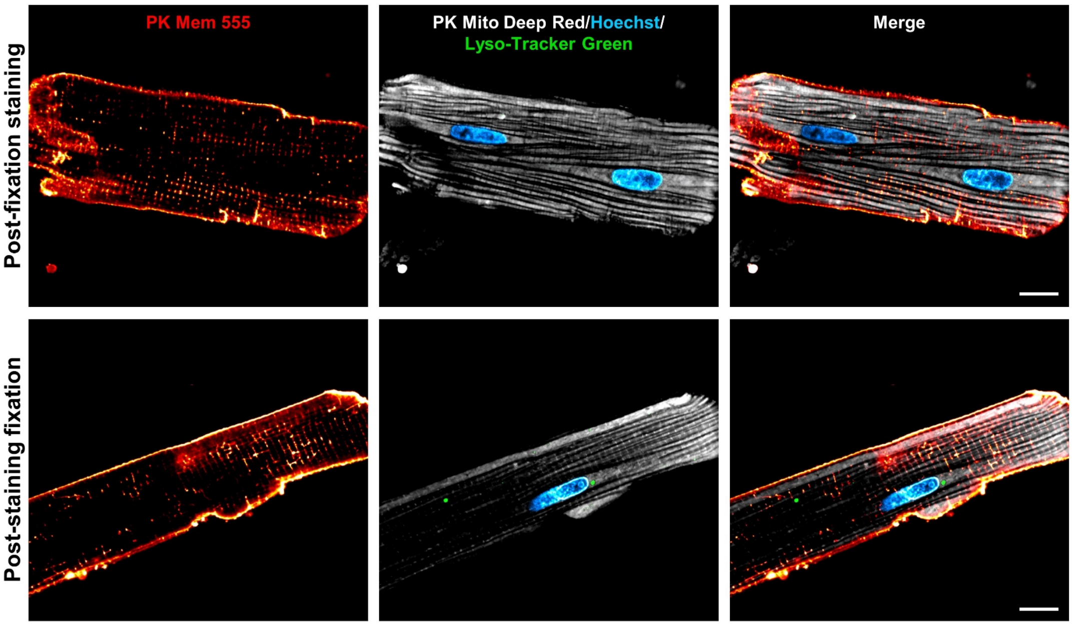

Figure S14. Adult rat cardiomyocytes labeled with PK Mem 555 (1 μM) before or after fixation were imaged with laser scanning confocal microscopy. The nucleus was stained with Hoechst. Lysosomes were stained with Lyso-Tracker Green. Mitochondria were stained with PK Mito Deep Red. Cardiomyocytes were fixed for 10 minutes at RT in 4% formaldehyde. Scale bar = 10 μm.

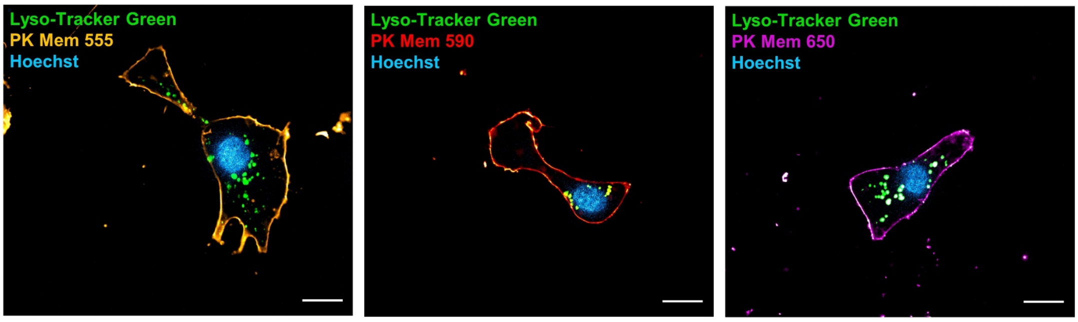

Figure S15. Laser scanning confocal microscopy images of neonatal rat cardiomyocytes 10 min after the addition of PK Mem dyes (200 nM) without washing. The nucleus was stained with Hoechst. Lysosome was stained with Lyso-Tracker Green. Scale bar = 10 μm.

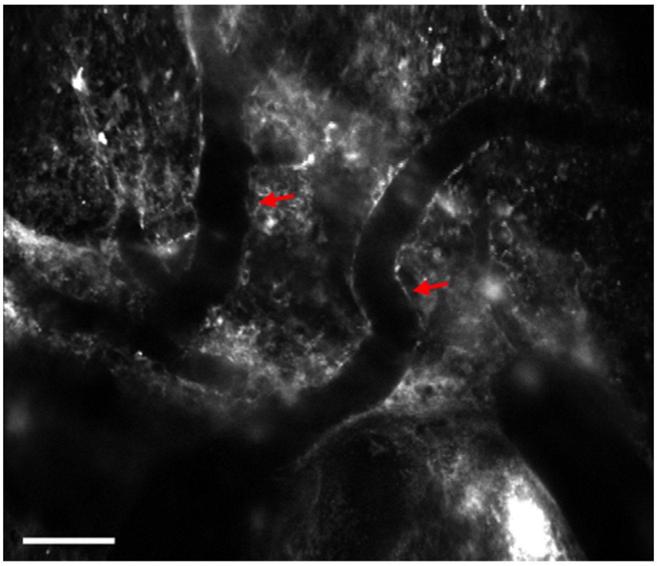

Figure S16. Two-photon image of a mouse brain slice labeled with PK Mem 555. Blood vessels are identified (red arrow). Scale bar = 50 μm.

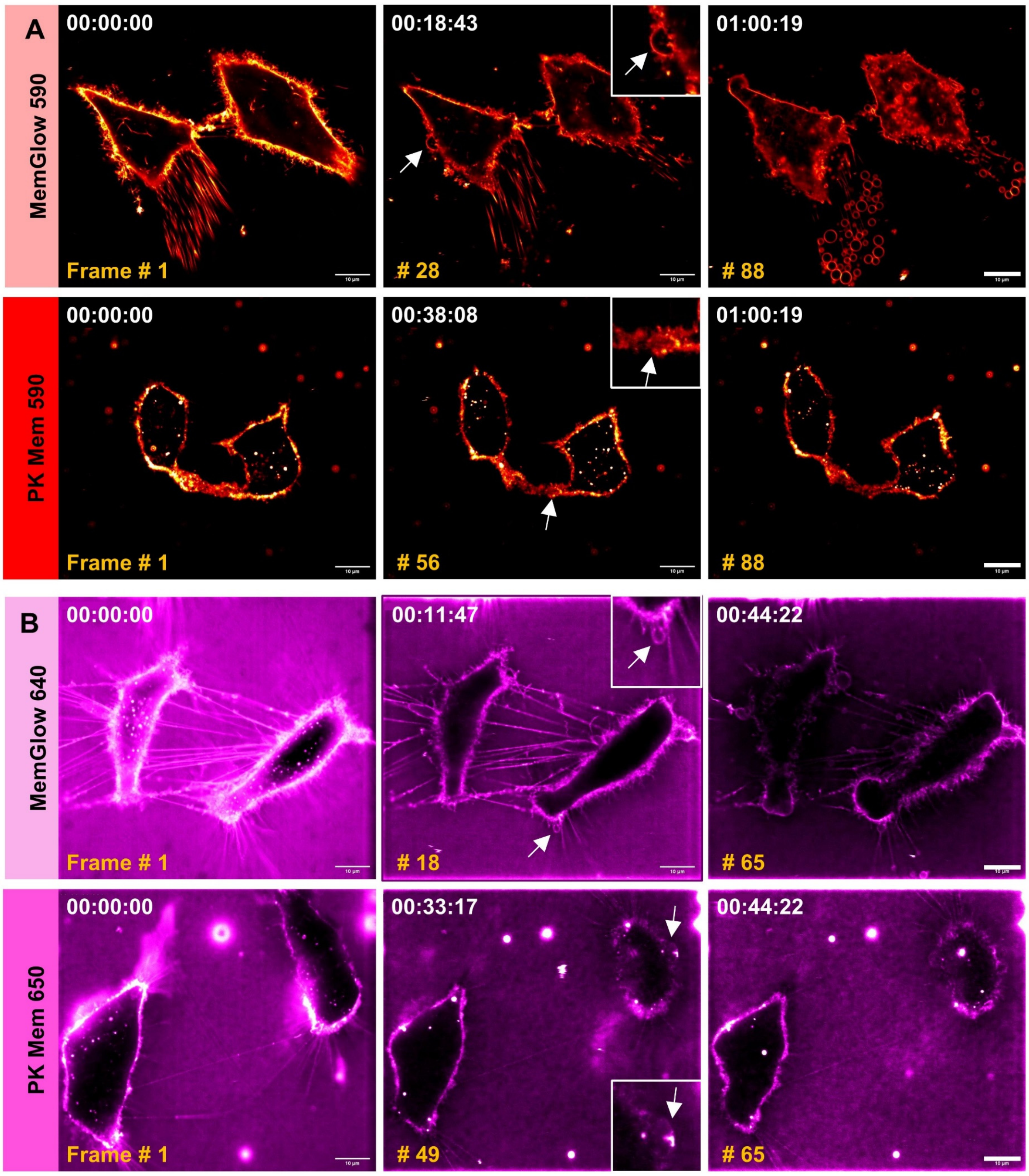

Figure S17. Long-term imaging of cell membrane morphology. (A) Time-lapse confocal recordings showing blebbing events of HeLa cells labeled with MemGlow 590 and PK Mem 590. Scale bar = 10 µm. (B) Time-lapse confocal recordings showing blebbing events of HeLa cells labeled with MemGlow 640 and PK Mem 650. Scale bar = 10 µm.

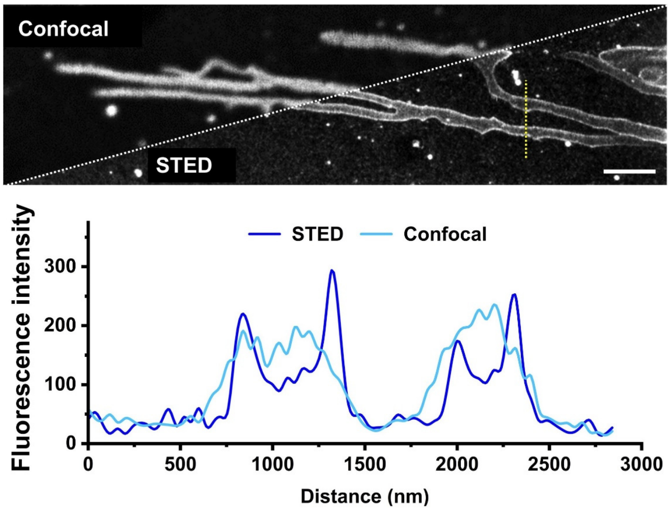

Figure S18. Top, confocal and STED images of membrane structures in live L929 cells. Bottom, intensity profiles corresponding to the yellow dotted line of the STED and confocal images. Scale bar = 2 μm.

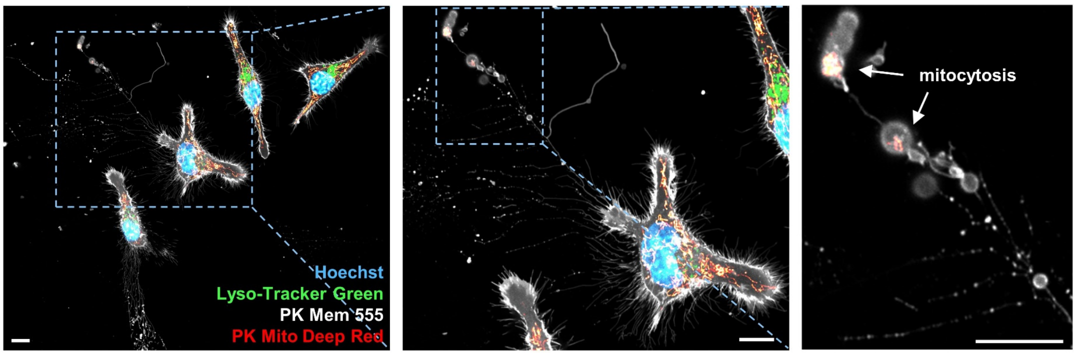

Figure S19. Left, LSCM of live L929 cells 5 min after addition of PK Mem 590 (1 μM) without washing. Middle, magnified view of the blue boxed area of the left image. Right, magnified view of the blue boxed area of the middle image. The nucleus was stained with Hoechst. Mitochondria were stained with PK Mito Deep Red. Lysosome was stained with Lyso-Tracker Green. Scale bars = 10 μm.

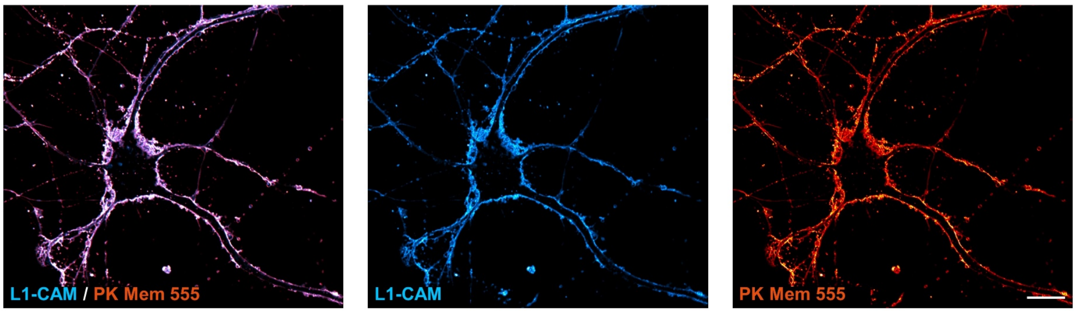

Figure S20. Laser scanning confocal microscopy images of primary hippocampal neurons incubated with PK Mem 555 (red) and L1-CAM monoclonal antibody (cyan, visualized with Goat Anti-Rabbit IgG AF647) for 10 min at 37°C. Scale bar = 10 μm.

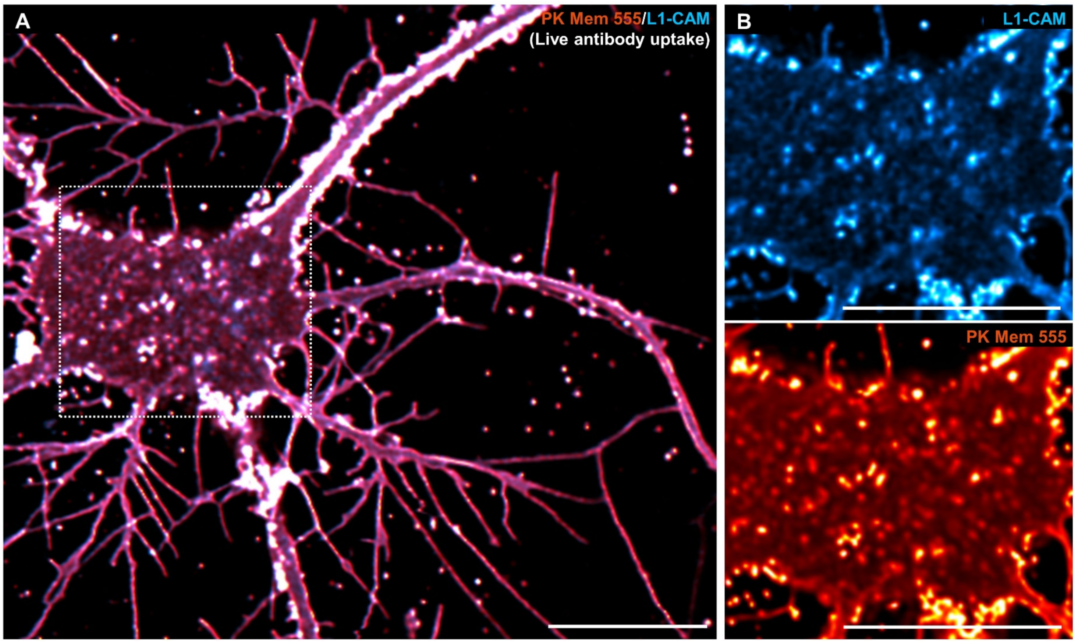

Figure S21. PK Mem 555 enables imaging of internalized vesicles. (A) LSCM images of the internalized vesicles of a primary hippocampal neuron tracked with L1-CAM monoclonal antibody (cyan, visualized with Goat Anti-Rabbit IgG AF647) and PK Mem 555 (red). Scale bar = 10 μm. (B) The white boxed area in Figure C is further magnified. Internalized vesicles can be tracked with L1-CAM antibody (top) or with PK Mem 555 (bottom). Scale bars = 10 μm.

### Supplementary Methods

#### UV-vis and fluorescence spectroscopy.

Stock solutions of PK Mem dyes were prepared in DMSO and diluted with MeOH to 1 μM. The absorption and emission spectra of PK Mem dyes in MeOH were measured using a Duetta fluorescence and absorbance spectrometer (Horiba, Kyoto, Japan) in a 1 cm square quartz cuvette. The concentration of PK Mem 555, 590, and 650 was determined based on the absorbance at 560 nm, 596 nm, and 654 nm, respectively. ($\varepsilon_{PK\ Mem\ 555,\ MeOH} = 1.7 \times 10^5\ M^{-1}\cdot cm^{-1}$, $\varepsilon_{PK\ Mem\ 590,\ MeOH} = 1.05 \times 10^5\ M^{-1}\cdot cm^{-1}$, $\varepsilon_{PK\ Mem\ 650,\ MeOH} = 2.6 \times 10^5\ M^{-1}\cdot cm^{-1}$)[1, 2].

#### Measurement of ROS generation of fluorophores

1,3-Diphenylisobenzofuran (DPBF) was applied to evaluate the ROS ($^1O_2$) generation of fluorophores. The mixed solution containing 1 μM PK Mem dyes was illuminated with an LED lamp with the corresponding wavelength (520-530 nm, 0.050 $W\cdot cm^{-2}$; 590-600nm, 0.050 $W\cdot cm^{-2}$; 620-630 nm, 0.0125 $W\cdot cm^{-2}$). At selected time points, the change of the UV–Vis absorption spectrum of DPBF was measured at 415 nm, giving a linear plot of the absorbance intensity versus time. TMRE and SiR-COOMe in MeOH were used as a reference. The quantum yield of $^1O_2$ was determined by comparing the slope for the samples with that obtained for the standards[3].

#### Preparation of DOPC small unilamellar vesicles (SUVs)

Dioleoylphosphatidylcholine (DOPC) SUVs were prepared as described previously[4]. Briefly, 15.7 mg DOPC was dissolved in 2 mL chloroform to obtain a 10 mM solution. Evaporate 0.5 mL DOPC solution in a 10 mL glass bottle under reduced pressure to form a layer of DOPC film. Completely evaporate residual chloroform by putting the bottle under a high vacuum at room temperature overnight. Add 0.5 mL PBS to the film and vortex vigorously to make an opaque solution. Sonicate the lipid mixture in a water-bath sonicator until it became clear, which indicated the formation of SUVs.

#### HeLa, L929 and KB cell culture

HeLa and L929 cells were incubated in Dulbecco's modified eagle medium (DMEM) supplemented with 10% v/v fetal bovine serum and 1% penicillin-streptomycin in an incubator at 37 °C with 5% $CO_2$. KB cells were incubated in Minimum Essential Medium (MEM) supplemented with 10% v/v fetal bovine serum and 1% penicillin-streptomycin in an incubator at 37 °C with 5% $CO_2$. For experiments, cells were seeded in confocal dishes (cellvis or Standard Imaging).

#### Cytotoxicity Assay

Cytotoxicity assay of the PK Mem dyes was performed using Cell Counting Kit 8 (CCK8). A total of $1\times10^4$ HeLa cells/well were plated in a 96-well plate 24 h prior to the cytotoxicity assay in Dulbecco's Modified Eagle Medium supplemented with 10% fetal bovine serum, 1% penicillin-streptomycin and were incubated in a 5% $CO_2$ incubator at 37°C. Then, an amount of 100 μL DMEM containing 20 nM, 200 nM, and 1000 nM of MemBright (PK Mem 555, PK Mem 590, and PK Mem 650) was added to HeLa cells and incubated for 1 h in the incubator. The cells were treated with 1% Triton as a positive control of cytotoxicity or DMEM having the same amount of DMSO (0.1% v/v) as the solution with the tested dyes as a control. After 1 h of dye incubation, 10 μl of the CCK-8 solution was added to each well of the plate. Then, the plate was incubated for 4 hours in the incubator. The absorbance at 450 nm was measured. In three independent experiments, sextuplicate samples of each dye concentration were evaluated. In comparison to the control DMEM+ 0.1% DMSO, the percentage of cell viability for each concentration was calculated.

**Mouse embryonic stem cell culture**

Clustered mESCs were incubated in DMEM supplemented with 15% FBS, 1% nucleosides (100×, Millipore), 1 mmol/L l-glutamine (Gibco), 1% nonessential amino acid (Gibco), 0.1 mmol/L 2-mercaptoethanol (Sigma), 1,000 U/mL LIF (Millipore), 3 μmol/L CHIR99021 (Stemcell), and 1 μmol/L PD0325901 (Stemcell) in 37°C and 5% $CO_2$.

**Neonatal rat neuron culture**

For primary rat hippocampus neuron culture, 14-mm glass coverslips were pre-coated with the poly-D-lysine solution and Laminin Mouse Protein solution at 37 °C within 2 days. After that, the coverslips were washed twice with $ddH_2O$ and let dry at room temperature. To isolate neurons, neonatal Sprague-Dawley rats' heads were cut off with scissors. Then, the brain was isolated from the skull and put into the ice-chilled dissection solution (DMEM with high glucose and penicillin-streptomycin antibiotics). Next, the hippocampi were separated from brains under a dissection scope, cut into small pieces, and incubated with Trypsin-EDTA (0.25%) for 15 min at 37°C. Thereafter, the trypsin was gently replaced with DMEM containing 10% FBS. After being repeatedly pipetted for 1 min and incubated on ice for 5 min, the tissue fragments were sedimented at the bottom of the centrifuge tube. The supernatant was collected and diluted with neural culture medium (Neurobasal™ medium supplemented with B-27™ supplement, GlutaMAX™ supplement, and penicillin-streptomycin) to a final cell density of $6\times10^4$ cells/mL. 1 mL cell suspension was added to each 24-well containing one coverslip. Every four days, half of the neural culture medium was replaced with the fresh medium.

**Isolation and culture of adult rat cardiomyocytes**

Adult male Sprague-Dawley rats weighing 150-180 g were anesthetized by intraperitoneal injection of 10% trichloroacetaldehyde monohydrate (0.3 mL/mg). Single ventricular myocytes were enzymatically isolated from the hearts. Freshly isolated cardiomyocytes were plated on laminin-coated culture dishes for 1 h and the attached cells were then maintained in M199 media (Sigma, M3769-1L) as described previously[5].

**Neonatal rat cardiomyocyte and myocardial fibroblast culture**

Neonatal rat cardiomyocytes were isolated from Sprague-Dawley rats born within 24 h. The whole hearts were isolated, minced and rinsed in Tyrode's buffer. Then, six cycles of digestion using collagenase type II (0.08% w/v, Worthington) and pancreatin (0.1% w/v, Sigma) were performed for each cycle for about 6 min at 37 °C. At the end of each cycle, the suspension was centrifuged for 5 min at 100 g and the supernatant was collected, pooled and resuspended in cardiomyocytes culture medium (DMEM containing 10% v/v FBS, 100 μM BrdU and penicillin-streptomycin solution). Isolated cells were pre-plated for 2 h on culture flasks in a humidified incubator at 37 °C with 5% $CO_2$ to separate cardiomyocytes and fibroblasts. Isolated myocardial fibroblasts were resuspended in DMEM containing 10% v/v FBS and penicillin-streptomycin. For experiments, myocardial fibroblasts were seeded in confocal dishes. Isolated cardiomyocytes were resuspended in a cardiomyocyte culture medium. The density of cardiomyocytes was adjusted to $1.5\times10^5$ cells/mL. After neonatal rat cardiomyocytes had been cultured for 36-48 h, the medium was rinsed for further experiments.

**Cellular Imaging**

Cells were seeded in glass-bottom dishes 24 hours prior to imaging. For single-color imaging, cells were stained with Opti-MEM supplemented with 20-200 nM PK Mem dyes at 37 °C for 5-10 min after rinsing with HBSS. Without any washing step, Cells could be imaged directly. For multi-color imaging, The nucleus was stained with Hoechst 33342(5 μg/mL) or SPY505-DNA in Opti-MEM and the cells were incubated for 10-30 minutes at 37°C, lysosome was stained with Lyso-Tracker-Green (1:20000) in Opti-MEM and the cells were incubated for 30 minutes at 37°C, tubulin was stained with SPY650-tubulin in Opti-MEM and the cells were incubated for 30 minutes at 37°C, and mitochondria were stained with PK Mito Red/ PK Mito Deep Red (250 nM) in Opti-MEM and the cells were incubated for 30 minutes at 37°C. After removing the staining solution, the cells were then washed with HBSS three times and stained with PK Mem dyes for imaging without washing. Prior to imaging, the PM was co-stained by the addition of WGA-488 at a final concentration of 5 μg/mL.

For fixed cells, staining was after fixation. Cells were seeded onto the confocal dishes, washed once with DPBS, and then fixed for 10 minutes at room temperature in 4% formaldehyde. Before the customary labeling and imaging, the preserved cells were washed three times with DPBS to remove the extra PFA. For fixable cells, staining followed by fixation is necessary for effective labeling. The cells were seeded onto the confocal dishes, stained with 20 nM of PK Mem dyes for 10 minutes at room temperature, and then fixed with 4% formaldehyde in HBSS for 10 minutes at room temperature before being rinsed three times with

HBSS.

**Brain slice imaging**

Mice were killed by cervical dislocation under ether anesthesia and the brain was immediately removed from the skull. The brain was sliced into 150 μm coronal sections in a cutting solution with a Vibratome (Leica). Brain slices were placed at 4°C in 1 mL of freshly prepared solution of PK Mem 555 (5 μM in ACSF) and 10 μg/mL Hoechst overnight. Slices were then mounted on glass slides and coverslipped. All images were acquired with an LSM700 Zeiss confocal laser scanning microscope (Zeiss, Göttingen, Germany). Single-section confocal images (1024 × 1024) were collected and exported in TIFF file format.

**In vivo fluorescent dye loading**

For loading PK Mem 555 into neurons, PK Mem 555 was dissolved in DMSO to obtain a 10 mM/L stock solution. This stock solution was mixed with ACSF (final concentration is 20 μM) and applied to the dura-free cortical surface within the craniotomy for 5 min.

**Miniature two-photon imaging**

For miniature two-photon experiments, a miniature hand-held cranial drill (China, RWD) was used to create a 4mm diameter window centered on the skull directly above the dye loading site. A ~4mm glass slide (thickness: 100 um) was then inserted into the skull window and glued to the skull with biological tissue glue (USA, 3M). After recovery, The miniature two-photon microscope (FHIRM-TPM V2.0, Field of view: 400 × 400 μm2; Resolution: ~850 nm; working distance: 1000 μm) was detachable while its holder was mounted permanently. Imaging data was acquired using the imaging software (GINKGO-MTPM, Transcend Vivoscope Biotech Co., Ltd, China) at a frame rate of 8.71 Hz (600 × 512 pixels) with a femtosecond fiber laser (~35 mW at the objective, TVS-FL-01, Transcend Vivoscope Biotech Co., Ltd, China). PK Mem 555 was excited at 920 nm

**Ethics approval and consent to participate**

Primary cell isolation procedures were approved by the Peking University Animal Use and Care Committee and complied with the standards of the Association for Assessment and Accreditation of Laboratory Animal Care (code number IMM-ChenZX-1).

The in vivo two-photon microscopy protocol has been reviewed and approved by the Animal Ethical and Welfare group PKU-Nanjing Joint Institute of Translational Medicine, Raygen Health Molecular Medicine Technology Co., Ltd.. (code number IACUC-2024-008).

**Imaging apparatus**

All confocal and STED imaging experiments were performed with Facility Line or

STEDYCON STED microscopes (Abberior Instruments GmbH, Göttingen, Germany) equipped with a CFI Plan Apochromat Lambda D 100x oil, NA1.45 objective (Nikon, Tokyo, Japan) or Plan Apochromat Lambda D 40x oil, NA1.3 objective (Nikon, Tokyo, Japan). PK Mem 555 was excited at 561 nm wavelength with dwell times of 3 µs. PK Mem 590 was excited at 561 nm wavelength and STED was performed using a pulsed depletion laser at 775 nm wavelength with gating of 1-7 ns and dwell times of 10 µs. PK Mem 555 was excited at 640 nm wavelength with dwell times of 3 µs. For multi-color confocal imaging of cells, Hoechst 33342 was excited at 405 nm wavelength, WGA-488, Lyso-Tracker Green, and SPY505-DNA were excited at 488 nm wavelength, PK Mito Red was excited at 560 nm wavelength, and PK Mito Deep Red and SPY650-tubulin were excited at 650 nm wavelength. The fluorescence signal was usually accumulated over 2-5 line steps. Imaging parameters were adjusted based on the individual samples. We typically used pixel sizes of 100 nm, line accumulations of 5 times, and dwell times of 10 µs in the confocal mode. In the STED mode, each line was scanned 25 times, and the pixel sizes were 20-40 nm. The pinhole was set to 0.7 - 1.0 AU.

All SIM images were collected through the Polar-SIM system (Airy Technology Co., Ltd., China) equipped with a SRHP Apo TIRF 100x oil, NA 1.49 objective (Nikon, Tokyo, Japan). A 561 nm laser was used to excite PK Mem 555 and a 640 nm laser was used to excite PK Mito Deep Red with GC modality.

**Image processing**

STED nanoscopy images were deconvoluted using Huygens software (Scientific Volume Imaging B.V., Hilversum, The Netherlands). STED nanoscopy images were deconvoluted using the Richardson-10 Lucy algorithm in the Imspector software (Abberior Instruments GmbH; version 0.14.11616). 3D reconstruction data were performed using Fiji/ImageJ (version 1.53t). Resolution estimation was performed by Gaussian fitting of fluorescence intensity line profiles. The full width at half maxima (FWHM) was estimated via Analysis >> Fitting >> Nonlinear curve fit >> Gaussian fit in the Origin Pro 2020b software (OriginLab Corporation, Northampton MA, USA). The SIM reconstruction process was conducted using the Airy-SIM software with pre-processing (Dark).

**Data Availability Statement**

Data presented in this study are provided in Supplementary Tables. Data are available from the corresponding authors upon request.

**General procedures on synthetic chemistry**

Unless otherwise stated, all commercially available materials were purchased at the highest commercial quality and used without further purification. Anhydrous dichloromethane (DCM), N, N-dimethylformamide (DMF), and pyridine (Py)were purchased from MREDA (China) or Innochem (China). All reactions were carried out with dry solvents, unless otherwise mentioned.

Reactions were monitored by Thin Layer Chromatography on plates (GF254) (Yantai Chemicals) using UV light as visualizing agent or by LC/MS (4.6 mm × 150 mm 5 μm C18 column; 2 μL injection; 5-100% $CH_3CN/H_2O$, linear gradient, with constant 0.1 % v/v formic acid additive; 6-10 min run; 0.5 mL/min flow; ESI; positive or negative ion mode; UV detection with ACQUITY PDA). If not specially mentioned, flash column chromatography uses silica gel (200-300 mesh, Tsingtao Haiyang Chemicals). Preparative HPLC separations were performed using Teledyne Isco EZ Prep UV-Vis and a RediSep Prep C18 column (100 Å, 5 μm, 20 × 150 mm).

NMR spectra were recorded on Brüker Advance 400 ($^1$H 400 MHz, $^{13}$C 101 MHz) and are calibrated using residual undeuterated solvent (Chloroform-d at 7.26 ppm $^1$H NMR, 77.16 ppm $^{13}$C NMR; $CD_3OD$ at 3.31 ppm $^1$H NMR, 49.00 ppm $^{13}$C NMR; DMSO-$d_6$ at 2.50 ppm $^1$H NMR, 39.52 ppm $^{13}$C NMR). Data for $^1$H NMR spectra are reported as follows: chemical shift (δ ppm), multiplicity (s = singlet, d = doublet, t = triplet, dd = doublet of doublets, m = multiplet, br = broad), coupling constant (Hz), integration. Data for $^{13}$C NMR are reported by chemical shift (δ ppm).

### Chemical synthesis

#### Supplementary Scheme 1. Synthetic route of PK Mem 590

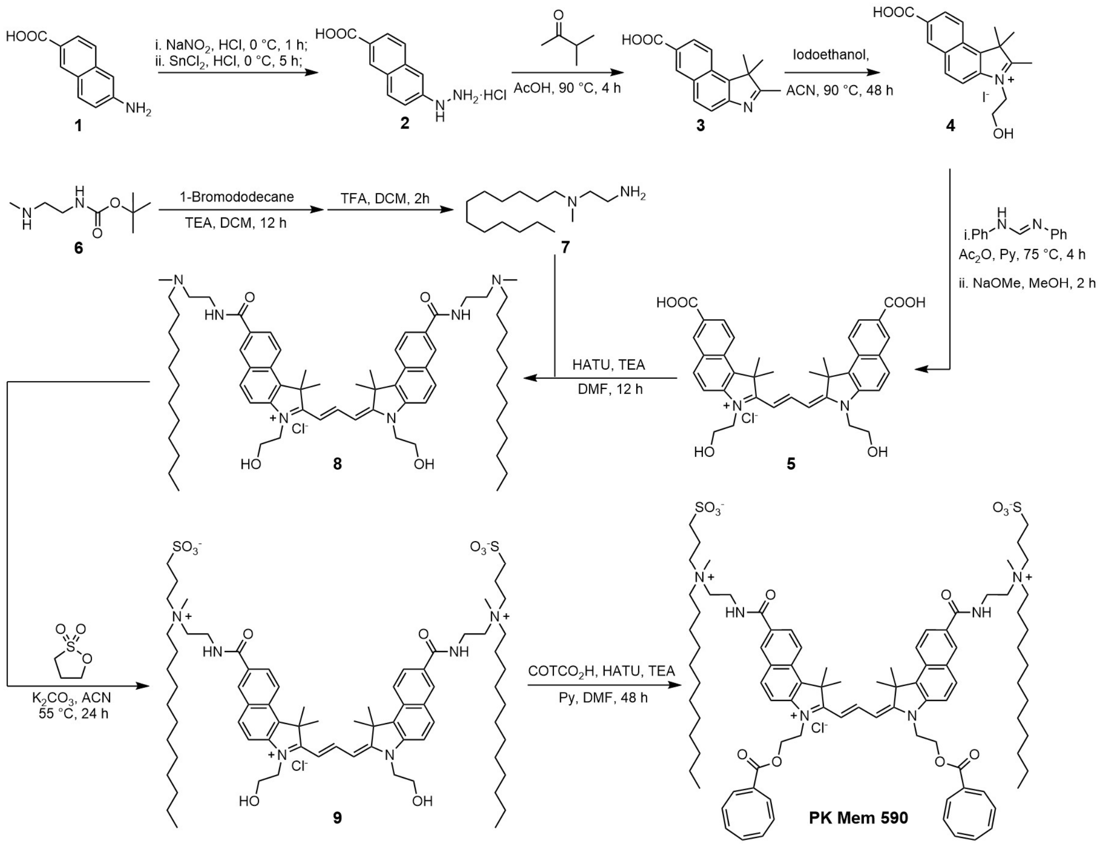

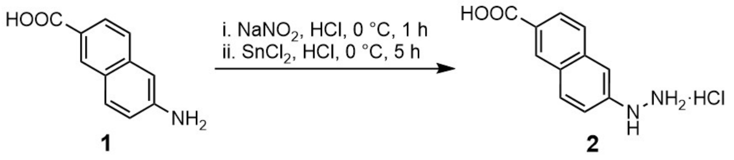

Compound **2**: $NaNO_2$ (243 mg, 3.53 mmol, 1.1 eq) in $H_2O$ was slowly added over a solution of 6-amino-2-naphthalenecarboxylic acid (Compound **1**; 600 mg, 3.21 mmol) in 3 mL conc. HCl cooled in an $H_2O$-ice bath. The resulting solution was stirred in an $H_2O$-ice bath for 1 h and a solution of $SnCl_2$ (3650 mg, 19.23 mol, 6 eq) in conc. HCl (2 mL) was added slowly. The resulting suspension was stirred in an $H_2O$-ice bath for 5 h and then filtered. The solid was successively washed with $H_2O$ (2×20 mL), with EtOH (20 mL), and with $Et_2O$ (2x20 mL). The solid was dried to afford compound **2** (568 mg, 2.38 mmol, 74% yield) as an off-white solid. $^1H$ NMR (400 MHz, DMSO-$d_6$) δ 10.51 (s, 3 H), 8.87 (s, 1 H), 8.49 (s, 1 H), 8.01 (d, $J$ = 8.8 Hz, 1 H), 7.93 (d, $J$ = 8.8 Hz, 1 H), 7.77 (d, $J$ = 8.8 Hz, 1 H), 7.31 (s, 1 H), 7.28 (d, $J$ = 8.8 Hz, 1 H). $^{13}C$ NMR (101 MHz, DMSO-$d_6$) δ 167.45, 145.45, 135.87, 130.50, 130.45, 127.61, 126.55, 126.18, 125.76, 117.46, 106.92. MS (ESI) calculated for $C_{11}H_{11}N_2O_2$ ($MH^+$) 203.2, observed 203.0.

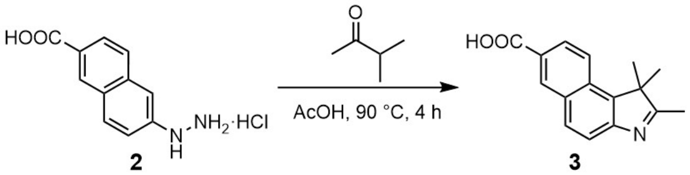

Compound **3**: 6-Hydrazinyl-2-naphthalenecarboxylic acid (Compound **2**; 500 mg, 2.09 mmol), 3-methyl-2-butanone (541 mg, 6.27 mmol, 3 eq) were dissolved in AcOH (3 mL). The resulted brown mixture was stirred for 30 min at room temperature and was heated to 90 °C and stirred at the same temperature for 4 h. After cooling the reaction mixture to room temperature, acetic acid was removed under reduced pressure and the residue was purified by column chromatography on $SiO_2$ (35 % EA/PE) to give compound **3** (502 mg, 1.98 mmol, 94% yield) as a yellow solid.

$^1$H NMR (400 MHz, DMSO-$d_6$) δ 8.70 (d, $J$ = 1.6 Hz, 1 H), 8.18 (d, $J$ = 8.8 Hz, 1 H), 8.10 (d, $J$ = 8.8 Hz, 1 H), 8.05 (dd, $J$ = 8.8, 1.6 Hz, 1 H), 7.77 (d, $J$ = 8.8 Hz, 1 H), 2.31 (s, 3 H), 1.45 (s, 6 H). $^{13}$C NMR (101 MHz, DMSO-$d_6$) δ 191.02, 167.44, 152.71, 138.84, 132.10, 130.73, 130.37, 130.04, 126.36, 125.81, 122.94, 120.04, 54.99, 22.00, 15.05. MS (ESI) calculated for $C_{16}H_{15}NO_2$ ($MH^+$) 254.1, observed 254.6.

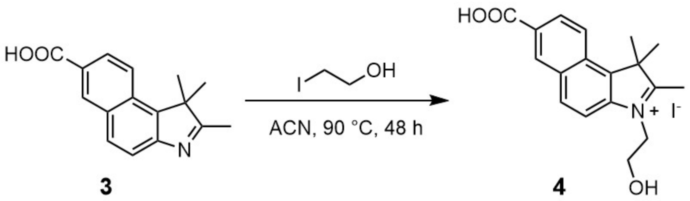

Compound **4**: 2-Iodoethanol (155 mg, 0.90 mmol, 3 eq) and 1,1,2-trimethyl-1*H*-benz[*e*]indole-7-carboxylic acid (Compound **3**; 76 mg, 0.30 mmol) were dissolved in ACN (5 mL) and the mixture was allowed to stirred at 90 °C for 48 h in a sealed tube. After cooling to room temperature, the solvent was removed under reduced pressure and the residue was washed with $Et_2O$ (2×20 mL) to obtain compound **4** (56 mg, 0.13 mmol, 44% yield) as a green solid.

$^1$H NMR (400 MHz, DMSO-$d_6$) δ 8.87 (s, 1 H), 8.50 (d, $J$ = 7.2 Hz, 2 H), 8.22 (t, $J$ = 8.4 Hz, 2 H), 4.74 (br, 2 H), 3.94 (br, 2 H), 2.94 (s, 3 H), 1.79 (s, 6 H). $^{13}$C NMR (101 MHz, DMSO-$d_6$) δ 198.93, 166.93, 140.35, 136.76, 132.33, 132.10, 129.04, 127.41, 123.91, 121.81, 115.00, 114.41, 57.99, 55.71, 50.56, 21.52, 14.34. MS (ESI) calculated for $C_{18}H_{20}NO_3$ ($M^+$) 298.1, observed 298.7.

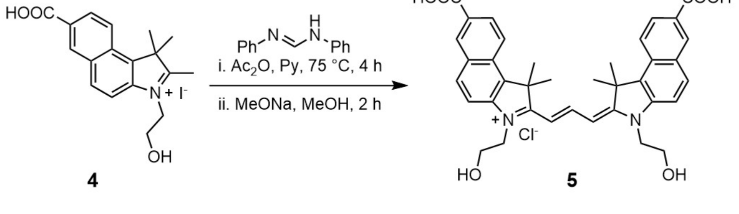

Compound **5**：N, N'-Diphenylformamidine (20 mg, 0.10 mmol) and compound **4** (91 mg, 0.21 mmol, 2.1 eq) were added in a mixture of $Ac_2O$ (2 mL) and 1 mL pyridine. The solution was stirred at 75 °C for 4 h. After cooling to room temperature, the solvent was removed under reduced pressure and the residue was dissolved to a solution of MeONa (43 mg, 0.80 mmol, 8 eq) in MeOH (3 mL) and stirred for 2 h. The solvent was removed under reduced pressure and the condensate was purified by HPLC (eluent, a 30 min linear gradient, from 30% to 70%

solvent B; flow rate, 10 mL/min; detection wavelength, 590 nm; eluent A ($ddH_2O$) and eluent B ($CH_3CN$)) to obtain compound **5** (35 mg, 0.055 mmol, 55% yield) as a purple solid.

$^1$H NMR (400 MHz, DMSO-$d_6$) δ 8.73 (s, 2 H), 8.67 (t, $J$ = 13.2 Hz, 1 H), 8.37 (d, $J$ = 8.8 Hz, 2 H), 8.30 (d, $J$ = 8.8 Hz, 2 H), 8.12 (d, $J$ = 6.0 Hz, 2 H), 7.84 (d, $J$ = 6.0 Hz, 2 H), 6.63 (d, $J$ = 9.2 Hz, 2 H), 4.38 (br, 4 H), 3.89 (br, 4 H), 2.03 (s, 12 H). $^{13}$C NMR (101 MHz, DMSO-$d_6$) δ 176.47, 167.14, 149.08, 142.31, 132.95, 132.51, 131.89, 130.61, 129.26, 127.01, 126.72, 122.53, 113.26, 103.27, 58.37, 50.62, 48.56, 27.06. MS (ESI) calculated for $C_{37}H_{37}N_2O_6$ ($M^+$) 605.3, observed 605.8.

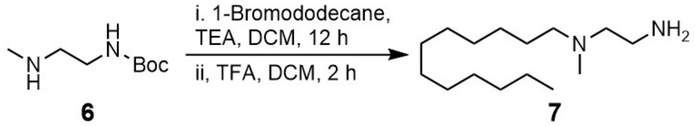

**6** **7**

Compound **7**: 1-Bromo-dodecane (8.58 g, 34.4 mmol, 2 eq) and triethylamine (6.97 g, 68.8 mmol, 4 eq) were added to a solution of tert-butyl-2-(methylamino)ethylcarbamate (Compound **6**; 3.0 g, 17.2 mmol) in DCM (20 mL). The mixture was stirred for 12 h at room temperature. The reaction was washed with brine and extracted with DCM (3 × 20 mL). The organic phase was combined and dried with $Na_2SO_4$, filtered and concentrated *in vacuo*. To the crude intermediate was added DCM (10 mL) and TFA (2 mL) and the solution was stirred for 2 h. The solvent was removed *in vacuo* and the crude product was purified by column chromatography on $SiO_2$ (1-6 % MeOH/DCM) to obtain compound **7** (1.80 g, 7.42 mmol, 43% yield in two steps) as a yellowish solid.

$^1$H NMR (400 MHz, Chloroform-$d$) δ 3.66 (s, 3H), 2.65 (t, $J$ = 6.0 Hz, 2H), 2.30 (t, $J$ = 6.0 Hz, 2H), 2.17 (t, $J$ = 7.6 Hz, 2H), 2.04 (s, 3H), 1.28 (t, $J$ = 6.4 Hz, 2H), 1.09 (s, 18 H), 0.71 (t, $J$ = 6.8 Hz, 2H). $^{13}$C NMR (101 MHz, Chloroform-$d$) δ 57.64, 54.33, 45.83, 41.05, 36.42, 32.06, 29.76, 29.72, 29.66, 29.50, 27.21, 25.91, 22.83, 14.26, 8.69. MS (ESI) calculated for $C_{15}H_{35}N_2$ ($MH^+$) 243.3, observed 243.7.

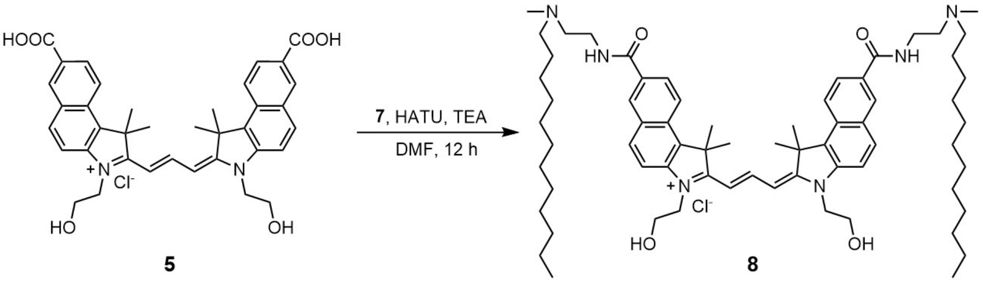

**5** **8**

Compound **8**: Compound **5** (15 mg, 0.023 mmol), HATU (26 mg, 0.069 mmol, 3 eq) and TEA (15 mg, 0.14 mmol, 6 eq) were dissolved in DMF (2 mL) and the solution was stirred for 10 min. Compound **7** (21 mg, 0.081 mmol, 4 eq) was added and the mixture was stirred for 12 h. The mixture was purified by HPLC (eluent, a 30 min linear gradient, from 50% to 80% solvent B; flow rate, 10 mL/min; detection wavelength, 590 nm; eluent A ($ddH_2O$) and eluent B ($CH_3CN$)) to obtain compound **8** (19 mg, 0.017 mmol, 74% yield) as a purple solid.

$^1$H NMR (400 MHz, Methanol-$d_4$) δ 8.87 (t, $J$ = 9.2 Hz, 1 H), 8.59 (d, $J$ = 0.8 Hz, 2 H), 8.40 (d, $J$ = 5.6 Hz, 2 H), 8.16 (d, $J$ = 6.0 Hz, 2 H), 8.13 (dd, $J$ = 5.6, 0.8 Hz, 2 H), 7.79 (d, $J$ = 6.0 Hz, 2 H), 6.58 (d, $J$ = 9.2 Hz, 2 H), 4.45 (t, $J$ = 3.6 Hz, 4 H), 4.06 (t, $J$ = 3.6 Hz, 4 H), 3.87 (t, $J$ = 6.0 Hz, 4 H), 3.01 (s, 6 H), 2.13 (s, 12 H), 1.78 (br, 4 H), 1.29 (br, 4 H), 1.26 (br, 36 H), 0.89 (t, $J$ = 4.8 Hz, 6 H). $^{13}$C NMR (101 MHz, Methanol-$d_4$) δ 178.75, 170.59, 151.49, 143.41,

134.83, 133.01, 132.61, 131.27, 130.87, 130.73, 126.80, 123.83, 113.98, 104.38, 60.28, 57.81, 57.00, 52.48, 48.24, 41.20, 36.50, 33.03, 30.69, 30.58, 30.46, 30.42, 30.18, 27.98, 27.50, 25.18, 23.70, 14.41. MS (ESI) calculated for $C_{67}H_{103}N_6O_4$ ($MH_2^{3+}$) 351.9, observed 352.2.

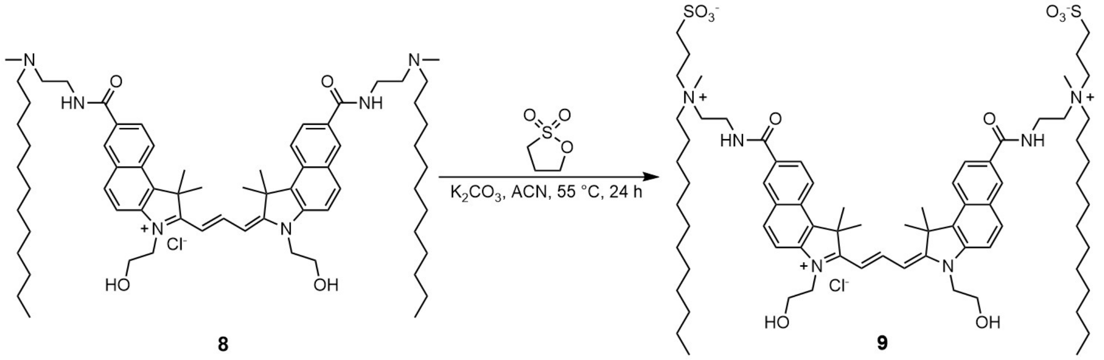

Compound **9**: Compound **8** (12 mg, 11.0 μmol) was dissolved in 1 mL ACN. 1,3-propanesultone (13 mg, 110 μmol, 10 eq) and $K_2CO_3$ (15 mg, 110 μmol, 10 eq) were added to the mixture. The reaction was heated to 55 °C for 24 h under stirring. The solvent was then removed under vacuum and the condensate was purified by HPLC (eluent, a 30 min linear gradient, from 50% to 70% solvent B; flow rate, 10 mL/min; detection wavelength, 590 nm; eluent A ($ddH_2O$) and eluent B ($CH_3CN$)) to obtain compound **9** (5 mg, 3.8 μmol, 35% yield) as a purple solid.

$^1$H NMR (400 MHz, Methanol-*d*4) δ 8.84 (t, *J* = 13.4 Hz, 1 H), 8.56 (s, 2 H), 8.38 (d, *J* = 8.9 Hz, 2 H), 8.17 (d, *J* = 8.9 Hz, 2 H), 8.10 (d, *J* = 9.0 Hz, 2 H), 7.77 (d, *J* = 9.0 Hz, 2 H), 6.59 (d, *J* = 13.5 Hz, 2 H), 4.44 (s, 4 H), 4.05 (t, *J* = 5.0 Hz, 4 H), 3.92 (t, *J* = 6.2 Hz, 4 H), 3.67 – 3.63 (m, 8H) 3.45 – 3.41 (m, 4H), 3.19 (s, 6 H), 2.93 (t, *J* = 6.6 Hz, 4 H), 2.33 – 2.24 (m, 4 H), 2.10 (s, 12 H), 1.85 (br, 4 H), 1.35-1.23 (m, 40 H), 0.88 (t, *J* = 6.9 Hz, 6 H). $^{13}$C NMR (101 MHz, Methanol-*d*4) δ 178.67, 169.72, 151.45, 143.39, 134.79, 133.08, 132.67, 131.16, 130.90, 130.69, 126.73, 123.93, 114.01, 104.43, 63.78, 62.03, 60.29, 52.44, 49.85, 48.23, 34.83, 33.06, 30.71, 30.61, 30.59, 30.46, 30.27, 27.99, 27.46, 23.73, 23.27, 19.65, 14.46. MS (ESI) calculated for $C_{73}H_{114}N_6O_{10}S_2$ ($MH^{2+}$) 649.4, observed 649.7.

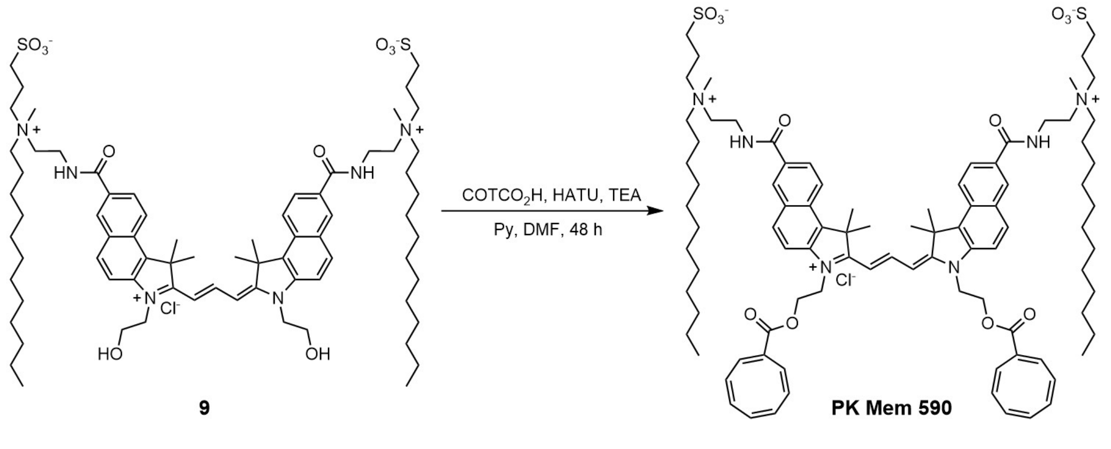

**PK Mem 590**: Cyclooctatetraenecarboxylic acid (2 mg, 15.0 μmol, 4 eq), HATU (7 mg, 18.7 μmol, 5 eq), TEA (5 mg, 45.0 μmol, 12 eq) and pyridine (0.2 mL) were added in DMF (1 mL) and the solution was stirred for 10 min. Compound **9** (5 mg, 3.75 μmol) was added and the mixture was stirred for 48 h. The mixture was purified by HPLC (eluent, a 30 min linear gradient, from 50% to 100% solvent B; flow rate, 10 mL/min; detection wavelength, 590 nm;

eluent A ($ddH_2O$) and eluent B ($CH_3CN$)) to obtain **PK Mem 590** (4 mg, 2.52 µmol, 67% yield) as a purple solid.
$^{1}H$ NMR (400 MHz, Methanol-$d_4$) δ 8.82 (t, $J$ = 13.4 Hz, 1H), 8.59 (s, 2H), 8.40 (d, $J$ = 9.0 Hz, 2H), 8.20 (d, $J$ = 9.0 Hz, 2H), 8.12 (d, $J$ = 9.0 Hz, 2H), 7.81 (d, $J$ = 8.9 Hz, 2H), 6.65 (d, $J$ = 13.5 Hz, 2H), 5.76 – 5.49 (m, 13H), 4.70 (br, 8H), 3.92 (t, $J$ = 6.2 Hz, 4H), 3.66 – 3.60 (m, 8H), 3.45 – 3.41 (m, 4H), 3.19 (s, 6H), 2.91 (t, $J$ = 6.5 Hz, 4H), 2.31 – 2.22 (m, 4H), 2.11 (s, 12H), 1.85 (s, 4H), 1.36 (br, 4H), 1.25 (br, 36H), 0.88 (t, $J$ = 6.8 Hz, 6H). $^{13}C$ NMR (101 MHz, Methanol-$d_4$) δ 178.57, 169.72, 166.54, 144.90, 142.88, 135.35, 134.86, 134.32, 133.77, 133.39, 132.52, 131.12, 130.69, 130.38, 129.98, 126.82, 123.98, 120.46, 116.19, 113.81, 104.67, 63.79, 62.02, 60.23, 52.52, 44.62, 34.83, 33.06, 30.71, 30.60, 30.58, 30.46, 30.26, 29.53, 28.03, 27.46, 23.73, 23.27, 19.63, 14.46. MS (ESI) calculated for $C_{91}H1_{26}N_6O_{12}S_2$ ($MH^{2+}$) 779.9, observed 779.6.

**Supplementary Scheme 2. Synthetic route of PK Mem 555**

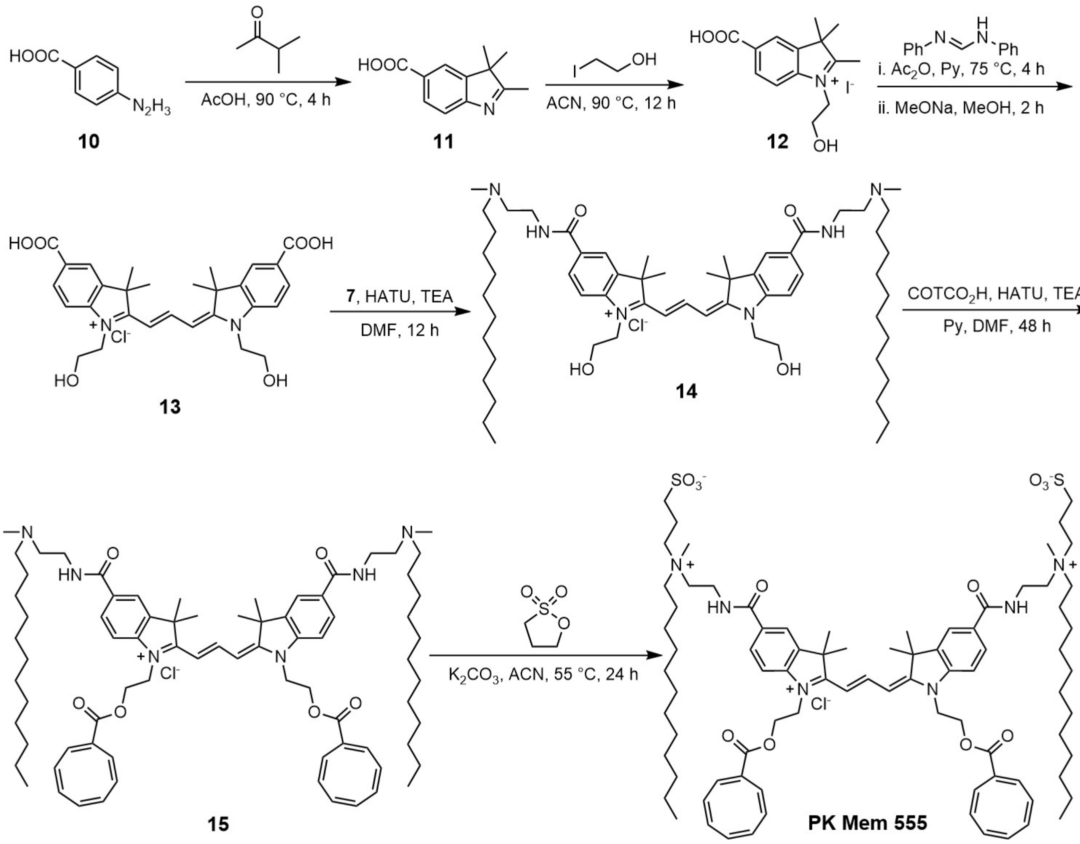

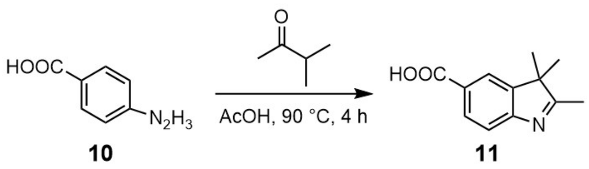

Compound **11**: 4-hydrazinobenzoic acid (Compound **10**; 200 mg, 1.32 mmol), 3-methyl-2-butanone (342 mg, 3.96 mmol, 3 eq) were dissolved in AcOH (5 mL). The resulted brown mixture was stirred for 30 min at room temperature and was heated to 90 °C and stirred at the same temperature for 4 h. After cooling the reaction mixture to room temperature, acetic acid was removed *in vacuo* and the residue was purified by column chromatography on $SiO_2$ (50 %

EA/PE) to obtain compound **11** (162 mg, 0.797 mmol, 60% yield) as a yellow solid.
[1]H NMR (400 MHz, DMSO-*d*6) δ 8.12 (s, 1 H), 8.00 (dd, *J* = 8.0, 1.2 Hz, 1 H), 7.61 (d, *J* = 8.0 Hz, 1 H), 4.63 (s, 2H), 3.88 (s, 2 H), 2.87 (s, 3 H), 1.59 (s, 6 H).[13]C NMR (101 MHz, DMSO-*d*6) δ 200.78, 166.43, 144.33, 142.17, 131.62, 130.31, 124,36, 115.86, 57.76, 54.60, 50.61, 21.83, 14.90. MS (ESI) calculated for $C_{12}H_{14}NO_2$ ($MH^+$) 204.1, observed 204.4.

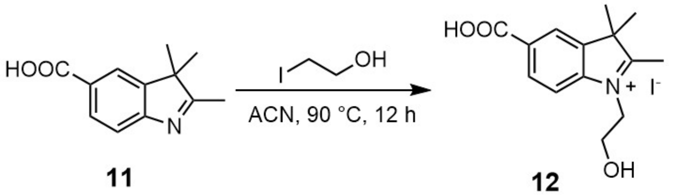

**11** **12**

Compound **12**: 2-Iodoethanol (169 mg, 0.98 mmol, 2 eq) and 2,3,3-trimethyl-3*H*-indole-5-carboxylic acid (Compound **11**; 100 mg, 0.49 mmol) were dissolved in ACN (5 mL) and the mixture was allowed to stirred at 90 °C for 12 h in a sealed tube. After cooling to room temperature, the solvent was removed under reduced pressure and the residue was washed with $Et_2O$ (2 × 20 mL) to obtain compound **12** (149 mg, 0.40 mmol, 81% yield) as a light pink salt.
[1]H NMR (400 MHz, DMSO-*d*6) δ 8.40 (s, 1 H), 8.18 (d, *J* = 8.0, 1.2 Hz, 2 H), 8.07 (d, *J* = 8.0 Hz, 2 H), 2.47 (s, 3 H), 1.37 (s, 6 H). [13]C NMR (101 MHz, DMSO-*d*6) δ 196.07, 167.37, 151.18, 144.92, 130.31, 129.41, 123.71, 118.07, 54.10, 22.20, 15.29. MS (ESI) calculated for $C_{14}H_{18}NO_3$ ($M^+$) 248.1, observed 248.5.

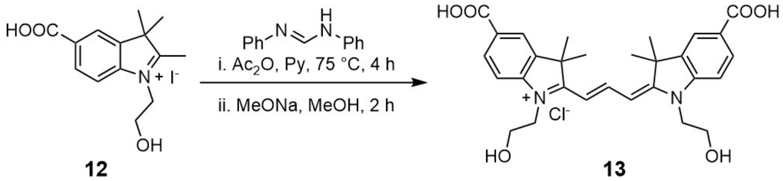

**12** **13**

Compound **13**: N, N'-Diphenylformamidine (8 mg, 0.041 mmol) and Compound **12** (30 mg, 0.082 mmol, 2 eq) was added to a mixture of $Ac_2O$ (2 mL) and pyridine (1 mL). The solution was stirred at 75 °C for 4 h. After cooling to room temperature, the solvent was removed under reduced pressure and the residue was dissolved to a solution of MeONa (17 mg, 0.32 mmol, 8 eq) in MeOH (2 mL) and stirred for 2 h. The solvent was removed *in vacuo* and the crude product was purified by HPLC (eluent, a 30 min linear gradient, from 30% to 70% solvent B; flow rate, 10 mL/min; detection wavelength, 550 nm; eluent A ($ddH_2O$) and eluent B ($CH_3CN$)) to obtain compound **13** (16 mg, 0.030 mmol, 73% yield) as a deep pink solid.
[1]H NMR (400 MHz, Methanol-*d*4) δ 8.63 (t, J = 13.6 Hz, 1 H), 8.12 (d, J = 1.2 Hz, 2 H), 8.10 (dd, J = 8.4, 1.2 Hz, 2 H), 7.44 (d, J = 8.4 Hz, 2 H), 6.55 (d, J = 13.6 Hz, 2 H), 4.31 (t, J = 5.2 Hz, 4 H), 3.96 (t, J = 5.2 Hz, 4 H), 1.79 (s, 12 H). [13]C NMR (101 MHz, Methanol-*d*4) δ 178.34, 168.98, 153.43, 147.71, 142.32, 132.20, 129.12, 124.67, 112.85, 105.80, 60.04, 50.67, 48.30, 28.24. MS (ESI) calculated for $C_{29}H_{33}N_2O_6$ ($M^+$) 505.2, observed 505.9.

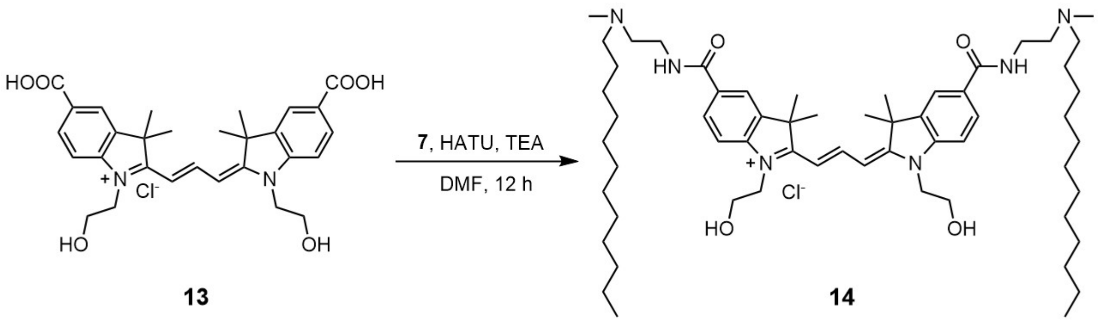

Compound **14**: Compound **13** (80 mg, 0.13 mmol), Hexafluorophosphate azabenzotriazole tetramethyl uronium (HATU, 145 mg, 0.38 mmol, 3 eq) and TEA (100 mg, 0.76 mmol, 6 eq) were dissolved in 5 mL DMF and the solution was stirred for 10 min. Compound **7** (90 mg, 0.38 mmol, 3 eq) was added and the mixture was stirred for 12 h. The mixture was directly subjected to HPLC purification (eluent, a 30 min linear gradient, from 50% to 90% solvent B; flow rate, 10 mL/min; detection wavelength, 550 nm; eluent A (dd$H_2O$) and eluent B ($CH_3CN$)) to obtain compound **14** (103 mg, 0.10 mmol, 75% yield) as a deep pink solid.

$^{1}$H NMR (400 MHz, Methanol-*d*$_4$) δ 8.62 (t, *J* = 13.2 Hz, 1 H), 8.03 (s, 2 H), 7.96 (d, *J* = 8.4 Hz, 2 H), 7.42 (d, *J* = 8.4 Hz, 2 H), 6.54 (d, *J* = 13.2 Hz, 2 H), 4.30 (br, 4 H), 3.77 (t, *J* = 4.8 Hz, 4 H), 3.77 (t, *J* = 5.6 Hz, 4 H), 3.50 – 3.43 (m, 2 H), 3.27 (s, 4 H), 3.15 – 3.11 (m, 2 H), 2.94 (s, 6 H), 1.79 (s, 12 H), 1.72 (t, *J* = 8 Hz, 4 H), 1.34 (br, 4 H), 1.25 (br, 36 H), 0.85 (t, *J* = 6.8 Hz, 6 H). $^{13}$C NMR (101 MHz, Methanol-*d*$_4$) δ 178.06, 170.09, 153.23, 147.04, 142,40, 131.66, 129.93, 122.68, 112.94, 105.73, 60.06, 57.75, 56.89, 50.74, 48.29, 41.07, 36.39, 33.04, 30.71, 30.60, 30.48, 30.44, 30.18, 28.25, 27.49, 25.13, 23.71, 14.43. MS (ESI) calculated for $C_{59}H_{99}N_6O_4$ ($MH_2^{3+}$) 318.6, observed 318.8.

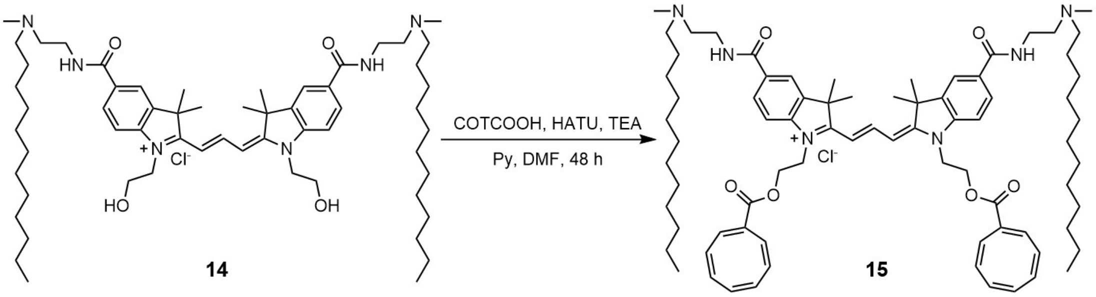

Compound **15**: Cyclooctatetraenecarboxylic acid (16 mg, 0.11 mmol, 4 eq), HATU (42 mg, 0.11 mmol, 4 eq), triethylamine (TEA, 29 mg, 0.22 mmol, 8 eq) and pyridine (0.5 mL) were added in DMF (2 mL) and the solution was stirred for 10 min. Compound **14** (30 mg, 0.028 mmol) was added and the mixture was stirred for 48 h. The mixture was purified by HPLC (eluent, a 30 min linear gradient, from 60% to 90% solvent B; flow rate, 10 mL/min; detection wavelength, 550 nm; eluent A (dd$H_2O$) and eluent B ($CH_3CN$)) to obtain compound **15** (21 mg, 0.017 mmol, 60% yield) as a deep pink solid.

$^{1}$H NMR (400 MHz, Methanol-*d*$_4$) δ 8.62 (t, *J* = 13.6 Hz, 1 H), 8.09 (d, *J* = 1.6 Hz, 2 H), 8.02 (dd, *J* = 8.4, 1.6 Hz, 2 H), 7.52 (d, *J* = 8.4 Hz, 2 H), 6.45 (d, *J* = 13.6 Hz, 2 H), 5.78 – 5.68 (m, 14 H), 4.66 (br, 4 H), 4.58 (br, 4 H), 3.82 (t, *J* = 5.6 Hz, 4 H), 3.51 – 3.48 (m, 2 H), 3.42 – 3.32 (m, 4 H), 3.18 – 3.12 (m, 2 H), 2.98 (s, 6 H), 1.81 (s, 12 H), 1.78 – 1.74 (m, 4 H), 1.38 (br, 4 H), 1.29 (br, 36 H), 0.89 (t, *J* = 6.8 Hz, 6 H). $^{13}$C NMR (101 MHz, Methanol-*d*$_4$) δ 178.01, 169.91, 166.54, 153.60, 146.42, 142.29, 135.40, 134.35, 133.84, 133.01, 132.56, 131.94,

130.90, 130.22, 130.07, 122.80, 112.86, 105.93, 61.83, 57.76, 56.88, 50.81, 47.88, 44.76, 41.10, 36.42, 33.05, 30.72, 30.60, 30.48, 30.45, 30.19, 28.26, 27.50, 25.16, 23.72, 14.44. MS (ESI) calculated for $C_{77}H_{111}N_6O_6$ ($MH_2^{3+}$) 405.3, observed 405.6.

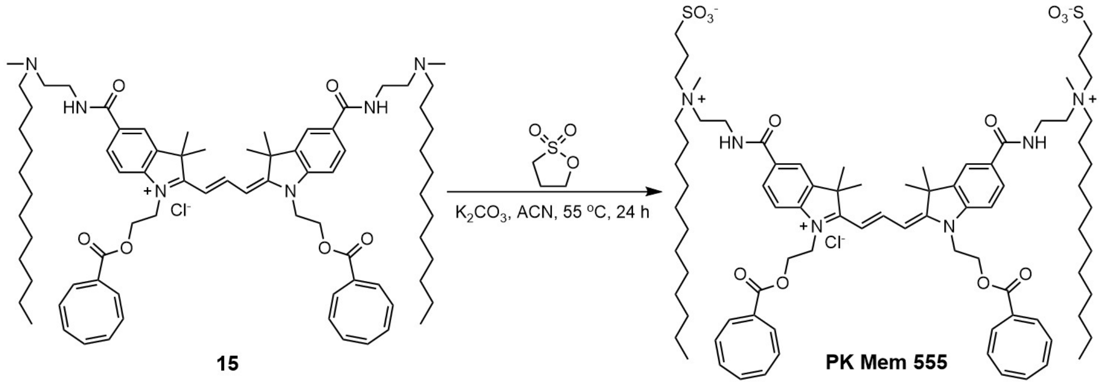

**PK Mem 555**: Compound **15** (30 mg, 0.024 mmol) was dissolved in ACN (5 mL). 1,3-propanesultone (15 mg, 0.12 mmol, 5 eq) and $K_2CO_3$ (17 mg, 0.12 mmol, 5 eq) have been added to the mixture. The reaction was heated to 55 °C for 24 h under stirring. The solvent was then removed under vacuum and the condensate was purified by HPLC (eluent, a 30 min linear gradient, from 50% to 80% solvent B; flow rate, 10 mL/min; detection wavelength, 550 nm; eluent A ($ddH_2O$) and eluent B ($CH_3CN$)) to obtain **PK Mem 555** (17 mg, 0.011 mmol, 47% yield) as a deep pink solid.

$^1$H NMR (400 MHz, Methanol-$d_4$) δ 8.62 (t, J = 13.2 Hz, 2 H), 8.09 (s, 2 H), 7.96 (d, $J$ = 8.0 Hz, 2 H), 7.53 (d, $J$ = 8.0 Hz, 2 H), 6.57 (d, $J$ = 12.4 Hz, 1 H), 5.81 – 5.67 (m, 14 H), 4.66 (br, 4 H), 4.59 (br, 4 H), 3.88 (br, 4 H), 3.66 – 3.62 (m, 4 H), 3.59 (br, 4 H), 3.18 (s, 6 H), 2.92 (t, $J$ = 5.6 Hz, 4 H), 2.27 (br, 4 H), 1.81 (br, 16 H), 1.38 (s, 4 H), 1.28 (s, 36 H), 0.89 (t, $J$ = 7.2 Hz, 6 H). $^{13}$C NMR (101 MHz, Methanol-$d_4$) δ 177.98, 169.11, 166.53, 153.59, 146.42, 144.94, 142.38, 135.40, 134.31, 133.84, 133.03, 132.58, 131.98, 130.87, 130.10, 122.74, 112.95, 105.98, 63.81, 61.94, 60.25, 50.85, 44.77, 34.77, 33.06, 30.74, 30.64, 30.59, 30.46, 30.26, 28.34, 27.45, 23.73, 23.23, 19.67, 14.47. MS (ESI) calculated for $C_{83}H_{122}N_6O_{12}S_2$ ($MH^{2+}$) 729.4, observed 729.9.

**Supplementary Scheme 3. Synthetic route of PK Mem 650**

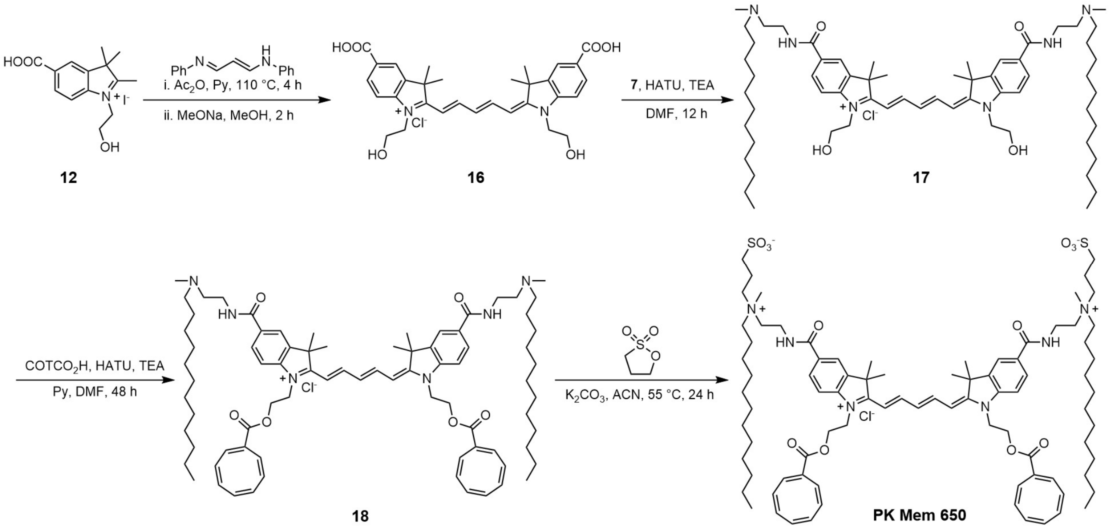

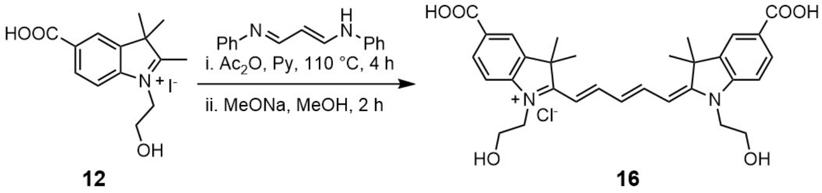

Compound **16**: N-(3-(phenylamino)allylidene)aniline (20 mg, 0.090 mmol) and compound **12** (74 mg, 0.20 mmol, 2.2 eq) was added in a mixture of 2 mL $Ac_2O$ and 1 mL pyridine. The solution was stirred at 110 °C for 4 h. After cooling to room temperature, the solvent was removed under reduced pressure and the residue was dissolved to a solution of MeONa (39 mg, 0.72 mmol, 8 eq) in 3 mL MeOH and stirred for 2 h. The solvent was removed under reduced pressure and the condensate was purified by HPLC (eluent, a 30 min linear gradient, from 40% to 70% solvent B; flow rate, 10 mL/min; detection wavelength, 650 nm; eluent A ($ddH_2O$) and eluent B ($CH_3CN$)) to obtain compound **16** (37 mg, 0.065 mmol, 72%) as a deep blue solid.
$^1$H NMR (400 MHz, Methanol-$d_4$) δ 8.35 (t, $J$ = 13.2 Hz, 2 H), 8.10 (dd, $J$ = 4, 2.4 Hz, 4 H), 7.41 (d, $J$ = 8.8 Hz, 2 H), 6.67 (d, $J$ = 12.4 Hz, 1 H), 6.45 (d, $J$ = 13.6 Hz, 2 H), 4.28 (t, $J$ = 5.2 Hz, 4 H), 3.95 (t, $J$ = 5.2 Hz, 4 H), 1.79 (s, 12 H). $^{13}$C NMR (101 MHz, Methanol-$d_4$) δ 176.58, 169.14, 156.61, 147.98, 142.71, 132.06, 128.44, 128.30, 124.51, 112.32, 106.27, 60.06, 50.46, 47.97, 27.89. MS (ESI) calculated for $C_{31}H_{35}N_2O_6$ ($M^+$) 531.3, observed 531.6.

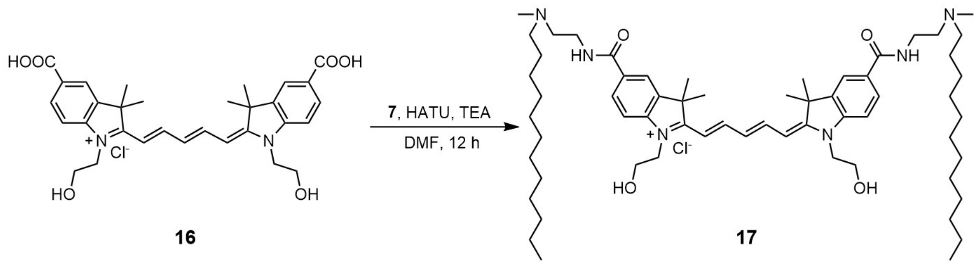

Compound **17**: Compound **16** (12 mg, 0.021 mmol), HATU (20 mg, 0.053 mmol, 2.5 eq) and TEA (13 mg, 0.13 mmol, 6 eq) were dissolved in 2 mL DMF and the solution was stirred for 10 min. Compound **7** (8 mg, 0.033 mmol, 3 eq) was added and the mixture was stirred for 12 h. The mixture was purified by HPLC (eluent, a 30 min linear gradient, from 50% to 80% solvent B; flow rate, 10 mL/min; detection wavelength, 650 nm; eluent A ($ddH_2O$) and eluent

B ($CH_3CN$)) to obtain compound **17** (11 mg, 0.011 mmol, 52%) as a deep blue solid.
$^1$H NMR (400 MHz, Methanol-$d_4$) δ 8.35 (t, *J* = 13.2 Hz, 2 H), 8.02 (d, *J* = 1.6 Hz, 2 H), 7.97 (dd, *J* = 8.4, 1.6 Hz, 2 H), 7.42 (d, *J* = 8.4 Hz, 2 H), 6.66 (d, *J* = 12.4 Hz, 1 H), 6.45 (d, *J* = 13.6 Hz, 2 H), 4.28 (t, *J* = 5.2 Hz, 4 H), 3.95 (t, *J* = 5.2 Hz, 4 H), 3.80 (t, *J* = 5.6 Hz, 4 H), 3.54 – 3.44 (m, 2 H), 3.36 (br, 4 H), 3.18 – 3.10 (br, 2 H), 2.97 (s, 6 H), 1.79 (s, 12 H), 1.74 (t, *J* = 8 Hz, 4 H), 1.37 (br, 4 H), 1.29 (br, 36 H), 0.89 (t, *J* = 6.8 Hz, 6 H). $^{13}$C NMR (101 MHz, Methanol-$d_4$) δ 176.37, 170.22, 156.55, 147.31, 142.84, 130.97, 129.76, 128.27, 122.55, 112.40, 106.20, 60.06, 57.75, 56.94, 50.55, 47.95, 47.95, 41.10, 36.41, 33.05, 30.71, 30.60, 30.48, 30.45, 30.19, 27.90, 27.50, 25.14, 23.72, 14.43. MS (ESI) calculated for $C_{61}H_{101}N_6O_4$ ($MH_2^{3+}$) 327.3, observed 327.2.

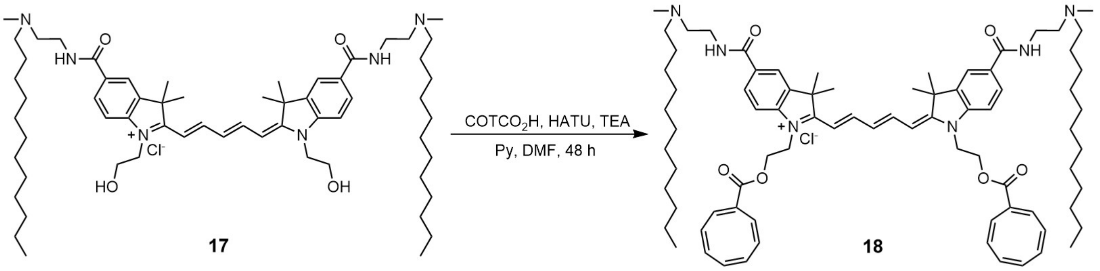

Compound **18**: Cyclooctatetraenecarboxylic acid (75 mg, 0.51 mmol, 3 eq), HATU (194 mg, 0.51 mmol, 3 eq), TEA (175 mg, 1.36 mmol, 8 eq) and 0.5 mL pyridine were added in DMF (5 mL) and the solution was stirred for 10 min. Compound **17** (176 mg, 0.17 mmol) was added and the mixture was stirred for 48 h. The mixture was purified by HPLC (eluent, a 30 min linear gradient, from 50% to 90% solvent B; flow rate, 10 mL/min; detection wavelength, 650 nm; eluent A ($ddH_2O$) and eluent B ($CH_3CN$)) to obtain compound **18** (152 mg, 0.12 mmol, 73% yield) as a deep blue solid.
$^1$H NMR (400 MHz, Methanol-$d_4$) δ 8.39 (t, *J* = 13.2 Hz, 2 H), 8.02 (s, 2 H), 7.99 (d, *J* = 8.4 Hz, 2 H), 7.44 (d, *J* = 8.4 Hz, 2 H), 6.69 (d, *J* = 12.4 Hz, 1 H), 6.49 (d, *J* = 13.6 Hz, 2 H), 5.83 – 5.70 (t, *J* = 21.6 Hz, 14 H), 4.62 (br, 4 H), 4.45 (br, 4 H), 3.81 (br, 4 H), 3.47 – 3.43 (m, 2 H), 3.35 (br, 4 H), 3.21 – 3.13 (m, 2 H), 2.98 (s, 6 H), 1.76 (s, 12 H), 1.70 (t, *J* = 8 Hz, 4 H), 1.37 (br, 4 H), 1.28 (br, 36 H), 0.89 (t, *J* = 6.8 Hz, 6 H). $^{13}$C NMR (101 MHz, Methanol-$d_4$) δ 176.40, 170.06, 166.61, 157.03, 146.63, 144.93, 142.75, 135.41, 134.35, 133.87, 133.04, 132.55, 131.18, 130.81, 130.01, 129.00, 122.65, 112.17, 106.48, 61.84, 57.77, 56.91, 50.60, 44.39, 41.14, 36.42, 33.03, 30.69, 30.58, 30.46, 30.43, 30.17, 27.85, 27.48, 25.15, 23.70, 14.43. MS (ESI) calculated for $C_{79}H_{113}N_6O_6$ ($MH_2^{3+}$) 414.0, observed 414.4.

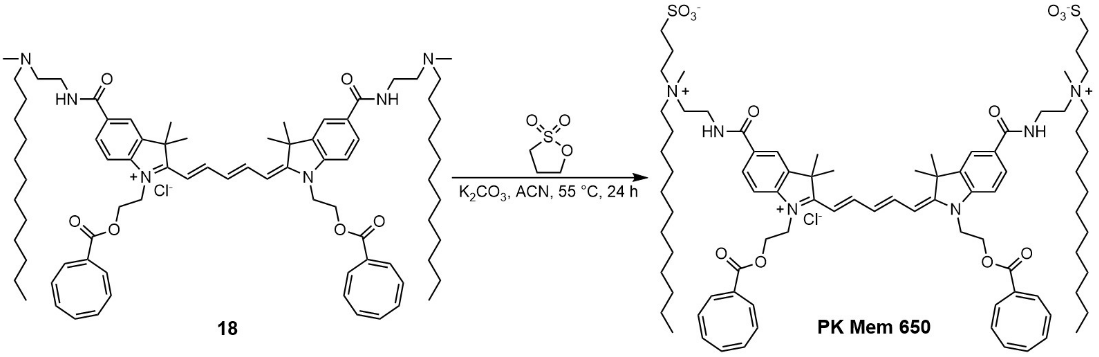

**PK Mem 650**: Compound **18** (30 mg, 0.023 mmol) was dissolved in ACN (2 mL). 1,3-Propanesultone (14 mg, 0.11 mmol, 5 eq) and $K_2CO_3$ (15 mg, 0.11 mmol, 5 eq) were added to the mixture. The reaction was heated to 55 °C for 24 h under stirring. The solvent was then removed under vacuum and the condensate was purified by HPLC (eluent, a 30 min linear gradient, from 40% to 80% solvent B; flow rate, 10 mL/min; detection wavelength, 650 nm; eluent A ($ddH_2O$) and eluent B ($CH_3CN$)) to obtain **PK Mem 650** (6 mg, 0.0039 mmol, 21%) as a deep blue solid.

$^1$H NMR (400 MHz, Methanol-$d_4$) δ 8.39 (t, $J$ = 13.2 Hz, 2 H), 8.00 (d, $J$ = 1.2 Hz, 2 H), 7.96 (dd, $J$ = 8.4, 1.2 Hz, 2 H), 7.46 (d, $J$ = 8.4 Hz, 2 H), 6.69 (d, $J$ = 12.4 Hz, 1 H), 6.49 (d, $J$ = 13.6 Hz, 2 H), 5.84 – 5.70 (m, 14 H), 4.63 (br, 4 H), 4.52 (br, 4 H), 3.86 (t, $J$ = 6 Hz, 4 H), 3.64 – 3.60 (m, 4 H), 3.58 – 3.53 (m, 4 H), 3.16 (s, 6 H), 2.89 (t, $J$ = 6.8 Hz, 4 H), 2.25 (t, $J$ = 6.4 Hz, 4 H), 1.85 – 1.82 (m, 4 H), 1.77 (s, 12 H), 1.38 (br, 4 H), 1.29 (br, 36 H), 0.89 (t, $J$ = 6.8 Hz, 6 H).$^{13}$C NMR (101 MHz, Methanol-$d_4$) δ 176.42, 169.37, 166.63, 157.04, 146.66, 142.85, 135.37, 134.44, 133.89, 133.05, 132.59, 131.31, 130.09, 129.84, 128.97, 126.29, 122.62, 112.28, 106.48, 63.85, 61.88, 60.27, 50.66, 44.42, 34.75, 33.05, 30.72, 30.62, 30.58, 30.45, 30.24, 27.91, 27.45, 25.24, 23.72, 23.23, 19.62, 14.44. MS (ESI) calculated for $C_{85}H_{124}N_6O_{12}S_2$ ($MH^{2+}$) 742.4, observed 743.0.

**Supplementary Scheme 4. Synthetic route of Az-Cy3.5**

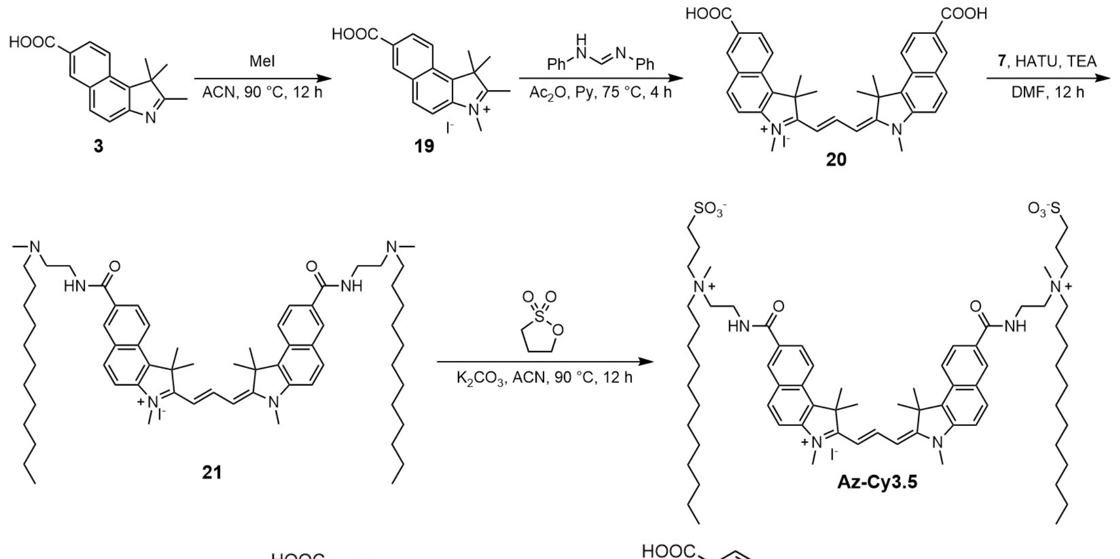

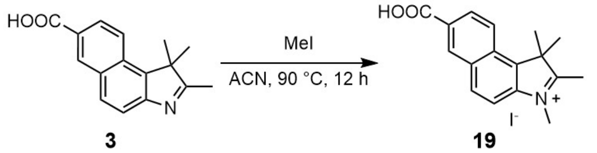

Compound **19**: 2-Iodoethanol (493 mg, 3.48 mmol, 3 eq) and 2,3,3-trimethyl-3*H*-naphindole-5-carboxylic acid (Compound **3**; 235 mg, 1.16 mmol) were dissolved in ACN (5 mL) and the mixture was allowed to stirred at 90 °C for 12 h in a sealed tube. After cooling to room temperature, the solvent was removed under reduced pressure and the residue was washed with $Et_2O$ (2 × 20 mL) to obtain compound **19** (287 mg, 0.84 mmol, 72% yield) as a light yellow salt.

$^1$H NMR (400 MHz, DMSO-$d_6$) δ 8.86 (s, 1H), 8.51 (d, $J$ = 8.9 Hz, 1H), 8.48 (d, $J$ = 8.9 Hz, 1H), 8.20 (dd, $J$ = 8.8, 3.2 Hz, 2H), 4.12 (s, 3H), 2.91 (s, 3H), 1.77 (s, 6H). $^{13}$C NMR (101 MHz, DMSO-$d_6$) δ 197.12, 166.92, 141.17, 136.40, 132.32, 132.14, 132.11, 128.93, 128.86, 127.37, 123.93, 114.08, 55.37, 35.24, 21.18, 14.24. MS (ESI) calculated for $C_{17}H_{18}NO_2$ ($M^+$) 268.1, observed 268.6.

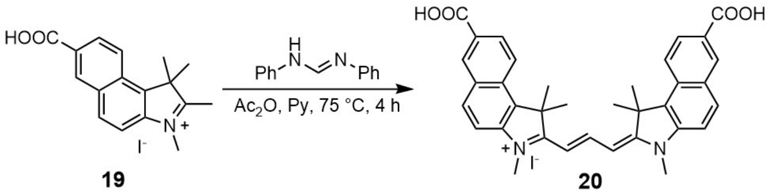

Compound **20**: N, N'-Diphenylformamidine (10 mg, 0.050 mmol) and compound **19** (30 mg, 0.10 mmol, 2 eq) were added in a mixture of 2 mL $Ac_2O$ and 1 mL pyridine. The solution was stirred at 75 °C for 4 h. After cooling to room temperature, the solvent was removed under reduced pressure and the condensate was purified by HPLC (eluent, a 30 min linear gradient, from 40% to 70% solvent B; flow rate, 10 mL/min; detection wavelength, 590 nm; eluent A ($ddH_2O$) and eluent B ($CH_3CN$)) to obtain compound **20** (21 mg, 0.037 mmol, 73%) as a purple solid.

$^1H$ NMR (400 MHz, Methanol-$d_4$) δ 8.35 (t, $J$ = 13.2 Hz, 2 H), 8.10 (dd, $J$ = 4, 2.4 Hz, 4 H), 7.41 (d, $J$ = 8.8 Hz, 2 H), 6.67 (d, $J$ = 12.4 Hz, 1 H), 6.45 (d, $J$ = 13.6 Hz, 2 H), 4.28 (t, $J$ = 5.2 Hz, 4 H), 3.95 (t, $J$ = 5.2 Hz, 4 H), 1.79 (s, 12 H). MS (ESI) calculated for $C_{35}H_{33}N_2O_4$ ($M^+$) 545.2, observed 545.8.

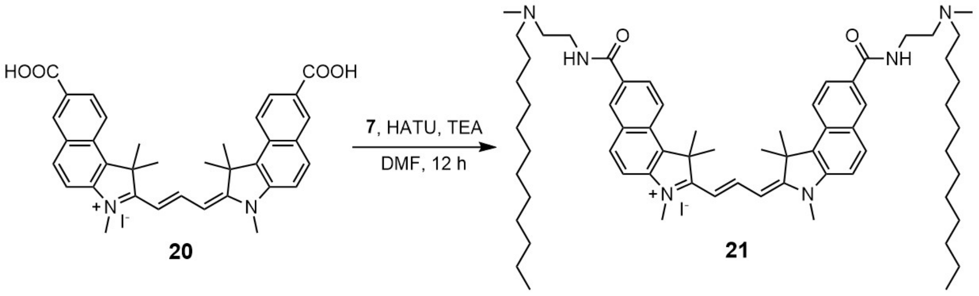

Compound **21**: Compound **20** (10 mg, 0.018 mmol), HATU (20 mg, 0.053 mmol, 3 eq) and TEA (11 mg, 0.11 mmol, 6 eq) were dissolved in 2 mL DMF and the solution was stirred for 10 min. Compound **7** (13 mg, 0.052 mmol, 3 eq) was added and the mixture was stirred for 12 h. The mixture was purified by HPLC (eluent, a 30 min linear gradient, from 50% to 80% solvent B; flow rate, 10 mL/min; detection wavelength, 590 nm; eluent A ($ddH_2O$) and eluent B ($CH_3CN$)) to obtain compound **21** (17 mg, 0.008 mmol, 46%) as a purple solid.

$^1H$ NMR (400 MHz, Methanol-$d_4$) δ8.81 (t, $J$ = 13.5 Hz, 1H), 8.59 (d, $J$ = 1.6 Hz, 2H), 8.39 (d, $J$ = 9.0 Hz, 2H), 8.20 (d, $J$ = 8.9 Hz, 2H), 8.13 (dd, $J$ = 8.9, 1.8 Hz, 2H), 7.78 (d, $J$ = 8.9 Hz, 2H), 6.54 (d, $J$ = 13.5 Hz, 2H), 3.88 – 3.84 (d, $J$ = 9.2 Hz, 10H), 3.56 – 3.52 (m, 2H), 3.22 – 3.17 (m, 2H), 3.01 (s, 6H), 2.11 (s, 12H), 1.81 – 1.73 (m, 4H), 1.38 (br, 4 H), 1.30 – 1.26 (m, 36H), 0.89 (t, $J$ = 6.9 Hz, 6H). MS (ESI) calculated for $C_{65}H_{99}N_6O_2$ ($MH_2^{3+}$) 331.9, observed 332.3.

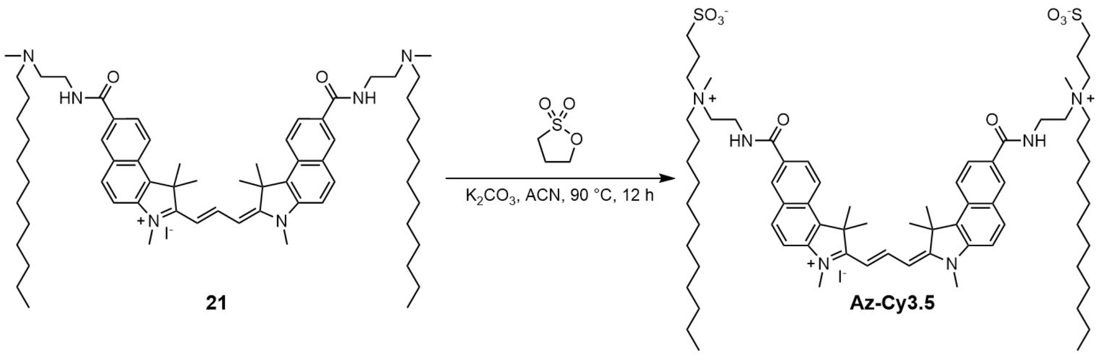

**Az-Cy3.5**: Compound **21** (38 mg, 0.018 mmol) was dissolved in ACN (2 mL). 1,3-Propanesultone (11 mg, 0.090 mmol, 5 eq) and $K_2CO_3$ (12 mg, 0.090 mmol, 5 eq) were added to the mixture. The reaction was heated to 90 °C for 12 h under stirring. The solvent was then removed under vacuum and the condensate was purified by HPLC (eluent, a 30 min linear gradient, from 40% to 80% solvent B; flow rate, 10 mL/min; detection wavelength, 590 nm; eluent A (dd$H_2O$) and eluent B ($CH_3CN$)) to obtain **Az-Cy3.5** (12 mg, 0.010 mmol, 55%) as a purple solid.

$^1$H NMR (400 MHz, Methanol-$d_4$) δ 8.81 (t, $J$ = 13.6 Hz, 1H), 8.58 (d, $J$ = 1.7 Hz, 2H), 8.40 (d, $J$ = 9.0 Hz, 2H), 8.22 (d, $J$ = 8.9 Hz, 2H), 8.12 (dd, $J$ = 8.9, 1.8 Hz, 2H), 7.77 (d, $J$ = 8.9 Hz, 2H), 6.54 (d, $J$ = 13.5 Hz, 2H), 3.92 (t, $J$ = 6.0 Hz, 4H), 3.84 (s, 6H), 3.68 – 3.59 (m, 8H), 3.45 – 3.41 (m, 4H), 3.18 (s, 6H), 2.90 (t, $J$ = 6.6 Hz, 4H), 2.28 – 2.24 (m, 4H), 1.88 – 1.83 (m, 4H), 1.36 (br, 4H), 1.26 (br, 36H), 0.89 (t, $J$ = 6.9 Hz, 6H). MS (ESI) calculated for $C_{71}H_{110}N_6O_8S_2$ ($MH^{2+}$) 619.4, observed 620.2.

**Supplementary Scheme 5. Synthetic route of Az-Cy3**

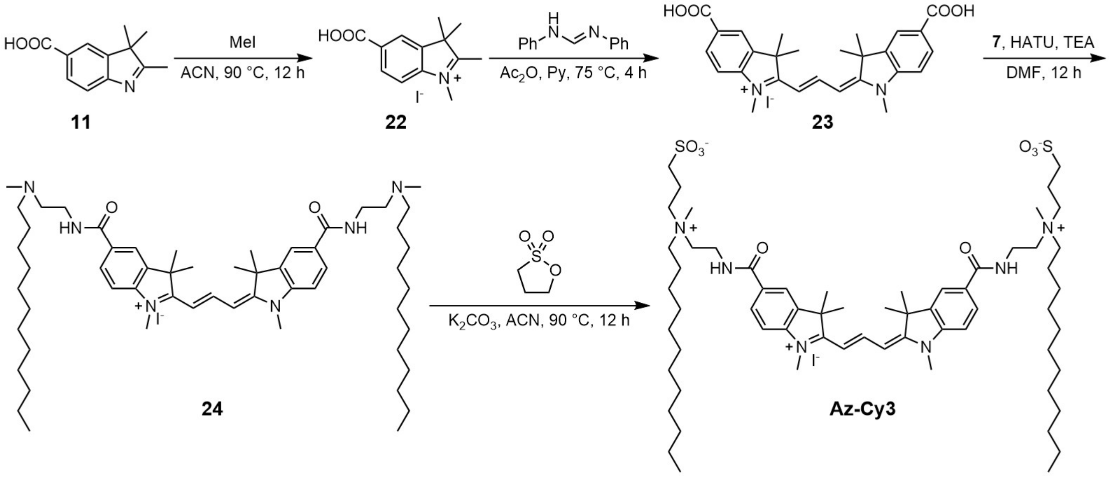

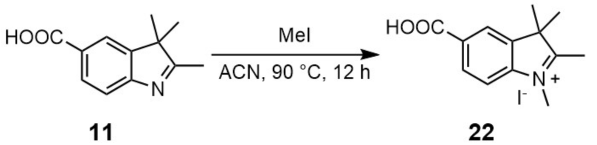

Compound **22**: 2-Iodoethanol (493 mg, 3.48 mmol, 3 eq) and 2,3,3-trimethyl-3*H*-indole-5-carboxylic acid (Compound **11**; 235 mg, 1.16 mmol) were dissolved in ACN (5 mL) and the mixture was allowed to stirred at 90 °C for 12 h in a sealed tube. After cooling to room temperature, the solvent was removed under reduced pressure and the residue was washed with

$Et_2O$ (2 × 20 mL) to obtain compound **22** (287 mg, 0.84 mmol, 72% yield) as a light yellow salt.

$^{1}H$ NMR (400 MHz, DMSO-$d_6$) δ 8.40 (s, 1 H), 8.18 (d, $J$ = 8.0, 1.2 Hz, 2 H), 8.07 (d, $J$ = 8.0 Hz, 2 H), 2.47 (s, 3 H), 1.37 (s, 6 H). $^{13}C$ NMR (101 MHz, DMSO-$d_6$) δ 196.07, 167.37, 151.18, 144.92, 130.31, 129.41, 123.71, 118.07, 54.10, 22.20, 15.29. MS (ESI) calculated for $C_{13}H_{16}NO_2$ ($M^+$) 218.1, observed 218.5.

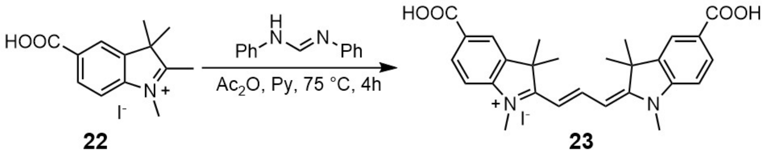

Compound **23**: N, N'-Diphenylformamidine (10 mg, 0.050 mmol) and compound **22** (30 mg, 0.10 mmol, 2 eq) were added in a mixture of 2 mL $Ac_2O$ and 1 mL pyridine. The solution was stirred at 75 °C for 4 h. After cooling to room temperature, the solvent was removed under reduced pressure and the condensate was purified by HPLC (eluent, a 30 min linear gradient, from 40% to 70% solvent B; flow rate, 10 mL/min; detection wavelength, 560 nm; eluent A ($ddH_2O$) and eluent B ($CH_3CN$)) to obtain compound **23** (21 mg, 0.037 mmol, 73%) as a deep pink solid.

$^{1}H$ NMR (400 MHz, Methanol-$d_4$) δ 8.61 (t, $J$ = 13.2 Hz, 2 H), 8.12-8.09 (m, 4 H), 7.42 (d, $J$ = 8.8 Hz, 2 H), 6.50 (d, $J$ = 12.4 Hz, 2 H), 3.69 (s, 6 H), 1.77 (s, 12 H). $^{13}C$ NMR (101 MHz, Methanol-$d_4$) δ 177.99, 168.92, 153.36, 147.74, 142.27, 132.39, 129.22, 124.66, 112.21, 105.33, 50.51, 49.00, 32.09, 28.07. MS (ESI) calculated for $C_{27}H_{29}N_2O_4$ ($M^+$) 445.2, observed 445.7.

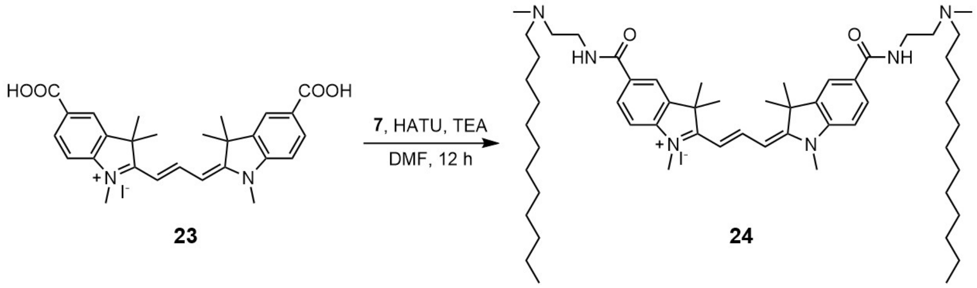

Compound **24**: Compound **23** (10 mg, 0.018 mmol), HATU (20 mg, 0.053 mmol, 3 eq) and TEA (11 mg, 0.11 mmol, 6 eq) were dissolved in 2 mL DMF and the solution was stirred for 10 min. Compound **7** (13 mg, 0.052 mmol, 3 eq) was added and the mixture was stirred for 12 h. The mixture was purified by HPLC (eluent, a 30 min linear gradient, from 50% to 80% solvent B; flow rate, 10 mL/min; detection wavelength, 560 nm; eluent A ($ddH_2O$) and eluent B ($CH_3CN$)) to obtain compound **24** (17 mg, 0.008 mmol, 46%) as a deep pink solid.

$^{1}H$ NMR (400 MHz, Methanol-$d_4$) δ 8.60 (t, $J$ = 13.5 Hz, 1H), 8.07 (d, $J$ = 1.5 Hz, 2H), 8.02 (dd, $J$ = 8.4, 1.7 Hz, 2H), 7.46 (d, $J$ = 8.4 Hz, 2H), 6.54 (d, $J$ = 13.5 Hz, 2H), 3.82 (t, $J$ = 5.8 Hz, 4H), 3.73 (s, 6H), 3.56 – 3.46 (m, 2H), 3.21 – 3.12 (m, 2H), 2.98 (s, 6H), 1.81 (s, 12H), 1.79 – 1.73 (m, 4H), 1.32 (m, 40H), 0.88 (t, $J$ = 6.8 Hz, 6H). $^{13}C$ NMR (101 MHz, Methanol-$d_4$) δ 177.71, 170.00, 153.14, 147.01, 142.34, 131.75, 130.13, 122.67, 112.26, 105.29, 57.72, 56.84, 50.57, 49.85, 41.06, 36.37, 33.04, 32.08, 30.71, 30.59, 30.47, 30.44, 30.17, 28.08, 27.48, 25.12, 23.71, 14.44. MS (ESI) calculated for $C_{57}H_{65}N_6O_2$ ($MH_2^{3+}$) 298.6, observed 299.0.

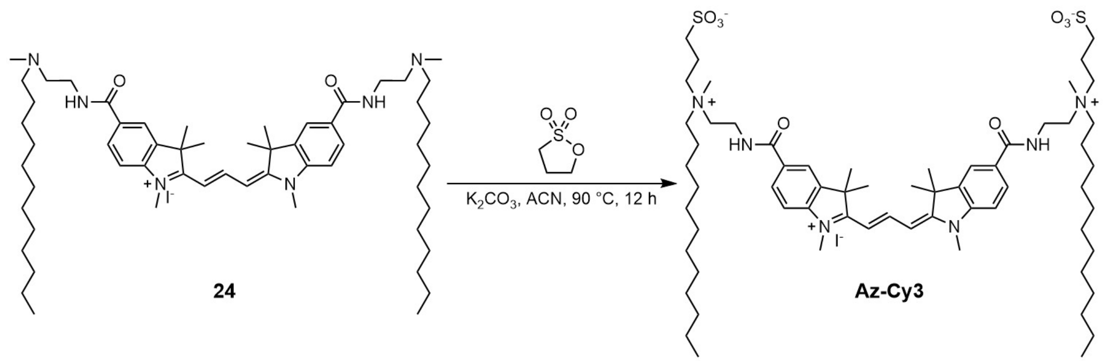

**Az-Cy3**: Compound **24** (38 mg, 0.018 mmol) was dissolved in ACN (2 mL). 1,3-Propanesultone (11 mg, 0.090 mmol, 5 eq) and $K_2CO_3$ (12 mg, 0.090 mmol, 5 eq) were added to the mixture. The reaction was heated to 90 °C for 12 h under stirring. The solvent was then removed under vacuum and the condensate was purified by HPLC (eluent, a 30 min linear gradient, from 40% to 80% solvent B; flow rate, 10 mL/min; detection wavelength, 560 nm; eluent A ($ddH_2O$) and eluent B (MeOH)) to obtain **Az-Cy3** (12 mg, 0.010 mmol, 55%) as a deep pink solid.

$^1$H NMR (400 MHz, Methanol-$d_4$) δ 8.60 (t, $J$ = 13.4 Hz, 1H), 8.12 – 8.06 (m, 2H), 8.02 (dd, $J$ = 8.4, 1.5 Hz, 2H), 7.48 (d, $J$ = 8.4 Hz, 2H), 6.54 (d, $J$ = 13.4 Hz, 2H), 3.88 (t, $J$ = 6.2 Hz, 4H), 3.73 (s, 6H), 3.70 – 3.54 (m, 8H), 3.42 (dd, $J$ = 10.8, 6.0 Hz, 4H), 3.18 (s, 6H), 2.91 (t, $J$ = 6.6 Hz, 4H), 2.36 – 2.20 (m, 4H), 1.81 (s, 16H), 1.37 – 1.32 (m, 40H), 0.88 (t, $J$ = 7.0 Hz, 6H). $^{13}$C NMR (101 MHz, Methanol-$d_4$) δ 177.73, 169.20, 153.14, 147.01, 142.45, 131.83, 130.05, 122.66, 112.37, 105.30, 63.74, 61.98, 60.27, 50.63, 49.85, 41.12, 34.78, 33.05, 32.12, 30.73, 30.63, 30.60, 30.49, 30.45, 30.25, 28.15, 28.12, 27.50, 27.44, 25.14, 23.72, 23.22, 19.65, 14.46. MS (ESI) calculated for $C_{63}H_{106}N_6O_8S_2$ ($MH^{2+}$) 569.4, observed 569.9.

**Supplementary Scheme 6. Synthetic route of Az-Cy5**

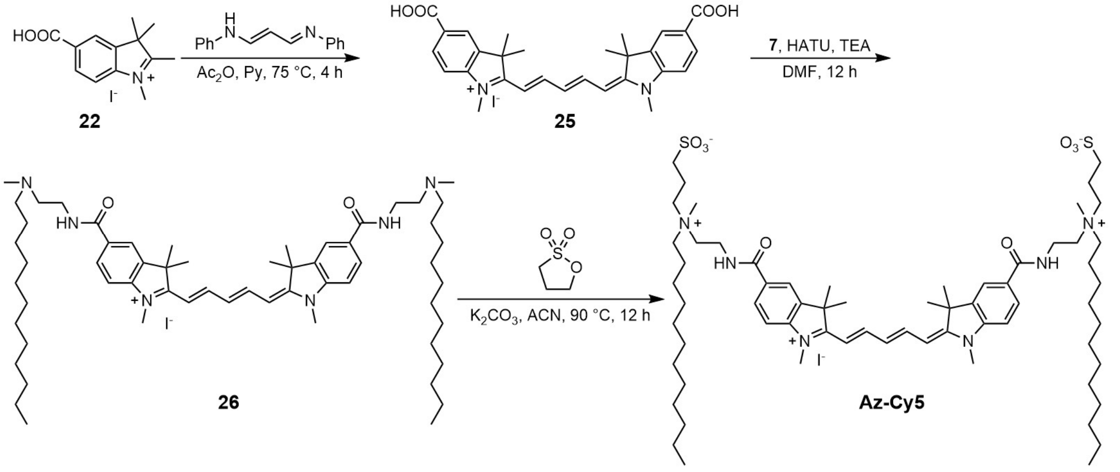

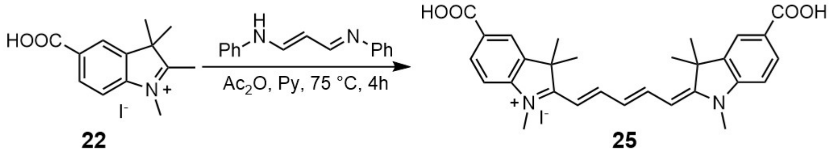

Compound **25**: N-(3-(phenylamino)allylidene)aniline (13 mg, 0.050 mmol) and compound **22** (30 mg, 0.10 mmol, 2 eq) was added in a mixture of 2 mL $Ac_2O$ and 1 mL pyridine. The solution was stirred at 75 °C for 4 h. After cooling to room temperature, the solvent was removed under reduced pressure and the condensate was purified by HPLC (eluent, a 30 min linear gradient, from 40% to 70% solvent B; flow rate, 10 mL/min; detection wavelength, 650 nm; eluent A (dd$H_2O$) and eluent B ($CH_3CN$)) to obtain compound **25** (13 mg, 0.022 mmol, 43%) as a deep blue solid.

$^1$H NMR (400 MHz, Methanol-*d₄*) δ 8.35 (t, *J* = 13.1 Hz, 2H), 8.16 – 8.06 (m, 4H), 7.38 (d, *J* = 8.3 Hz, 2H), 6.71 (t, *J* = 12.4 Hz, 1H), 6.37 (d, *J* = 13.7 Hz, 2H), 3.67 (s, 6H), 1.76 (s, 12H). $^{13}$C NMR (101 MHz, Methanol-$d_4$) δ 176.35, 169.12, 156.72, 148.01, 142.66, 132.28, 128.54, 128.27, 124.54, 111.67, 105.80, 50.32, 31.78, 27.71. MS (ESI) calculated for $C_{29}H_{31}N_2O_4$ ($M^+$) 471.2, observed 471.8.

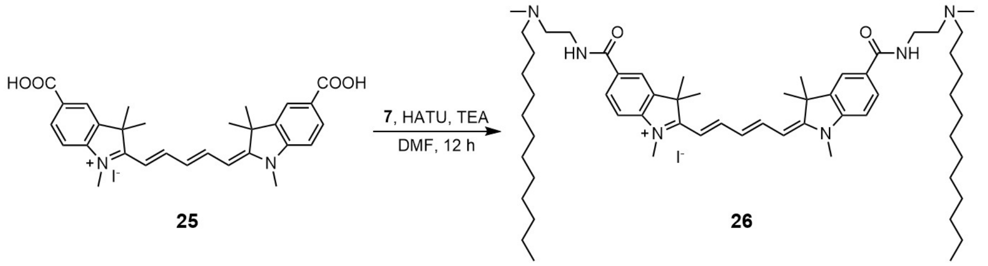

Compound **26**: Compound **25** (70 mg, 0.12 mmol), HATU (111 mg, 0.29 mmol, 2.5 eq) and TEA (71 mg, 0.71 mmol, 6 eq) were dissolved in 5 mL DMF and the solution was stirred for 10 min. Compound **7** (85 mg, 0.35 mmol, 3 eq) was added and the mixture was stirred for 12 h. The mixture was purified by HPLC (eluent, a 30 min linear gradient, from 50% to 80% solvent B; flow rate, 10 mL/min; detection wavelength, 650 nm; eluent A (dd$H_2O$) and eluent B ($CH_3CN$)) to obtain compound **26** (73 mg, 0.070 mmol, 60%) as a deep blue solid.

$^1$H NMR (400 MHz, Methanol-$d_4$) δ 8.35 (t, *J* = 13.1 Hz, 2H), 8.06 – 7.95 (m, 4H), 7.40 (d, *J* = 8.4 Hz, 2H), 6.72 (t, *J* = 12.4 Hz, 1H), 6.37 (d, *J* = 13.7 Hz, 2H), 3.80 (t, *J* = 5.8 Hz, 4H), 3.66 (s, 6H), 2.98 (s, 6H), 1.77 (br, 16H), 1.37 - 1.28 (m, 38H), 0.89 (t, *J* = 6.8 Hz, 6H).$^{13}$C NMR (101 MHz, Methanol-$d_4$) δ 176.14, 170.23, 156.65, 147.33, 142.79, 131.06, 129.97, 128.27, 122.59, 111.73, 105.75, 57.75, 56.97, 54.80, 50.41, 49.85, 41.13, 36.43, 33.06, 31.78, 30.72, 30.61, 30.56, 30.49, 30.46, 30.20, 27.73, 27.49, 25.15, 23.72, 14.43. MS (ESI) calculated for $C_{59}H_{97}N_6O_2$ ($MH_2^{3+}$) 307.3, observed 307.6.

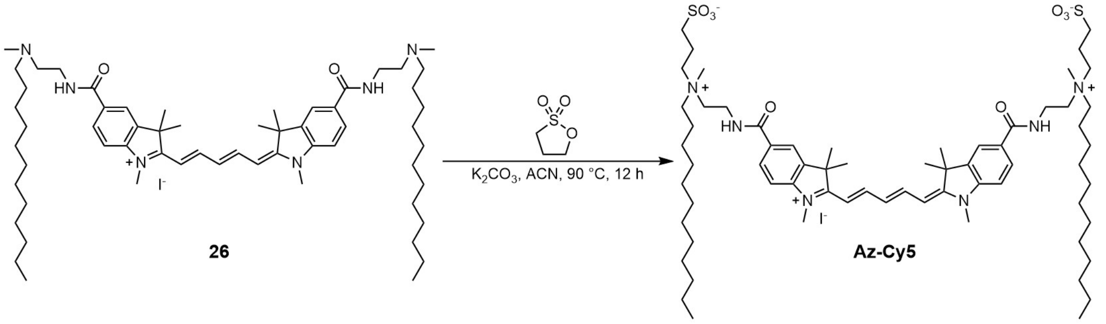

**Az-Cy5**: Compound **26** (40 mg, 0.038 mmol) was dissolved in ACN (2 mL). 1,3-Propanesultone (23 mg, 0.19 mmol, 5 eq) and $K_2CO_3$ (26 mg, 0.19 mmol, 5 eq) were added to the mixture. The reaction was heated to 90 °C for 12 h under stirring. The solvent was then removed under vacuum and the condensate was purified by HPLC (eluent, a 30 min linear gradient, from 40% to 80% solvent B; flow rate, 10 mL/min; detection wavelength, 650 nm; eluent A (dd$H_2O$) and eluent B ($CH_3CN$)) to obtain **Az-Cy5** (19 mg, 0.015 mmol, 39%) as a deep blue solid.

$^{1}H$ NMR (400 MHz, Methanol-$d_4$) δ 8.32 (t, $J$ = 13.0 Hz, 2H), 8.01 (s, 2H), 7.98 (d, $J$ = 8.4 Hz, 2H), 7.41 (d, $J$ = 8.4 Hz, 2H), 6.69 (t, $J$ = 12.3 Hz, 1H), 6.34 (d, $J$ = 13.6 Hz, 2H), 3.85 (t, $J$ = 6.4 Hz, 4H), 3.66 (s, 6H), 3.66 – 3.61 (m, 4H), 3.61 – 3.54 (m, 4H), 3.45 – 3.38 (m, 4H), 3.17 (s, 6H), 2.91 (t, $J$ = 6.3 Hz, 4H), 2.26 (dt, $J$ = 14.6, 6.4 Hz, 4H), 1.82 (s, 4H), 1.75 (s, 12H), 1.31 (d, $J$ = 34.7 Hz, 40H), 0.89 (t, $J$ = 6.8 Hz, 6H). $^{13}C$ NMR (101 MHz, Methanol-$d_4$) δ 176.07, 169.30, 169.27, 156.55, 147.27, 142.85, 131.11, 129.84, 128.18, 122.52, 111.84, 105.73, 63.74, 61.99, 60.23, 50.42, 49.85, 41.14, 34.75, 33.06, 31.83, 30.74, 30.64, 30.61, 30.50, 30.46, 30.27, 30.20, 27.80, 27.50, 27.45, 25.15, 23.73, 23.24, 19.64, 14.47. MS (ESI) calculated for $C_{65}H_{108}N_6O_8S_2$ ($MH^{2+}$) 582.4, observed 582.9.

**NMR spectra and LC-MS**

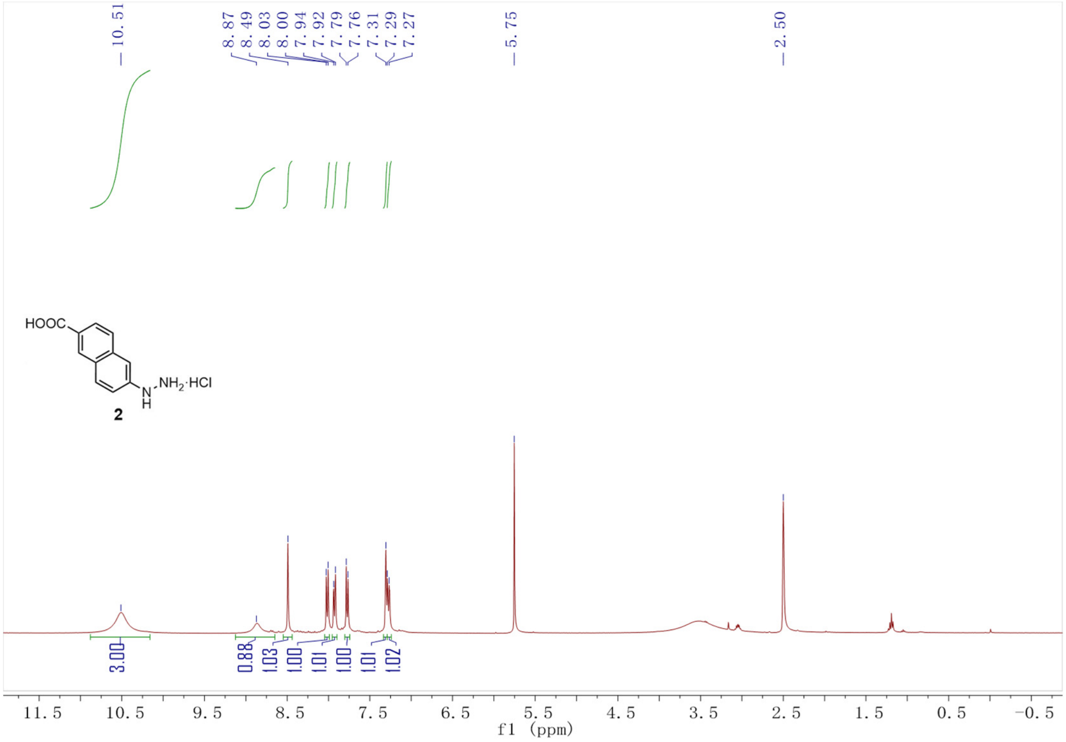

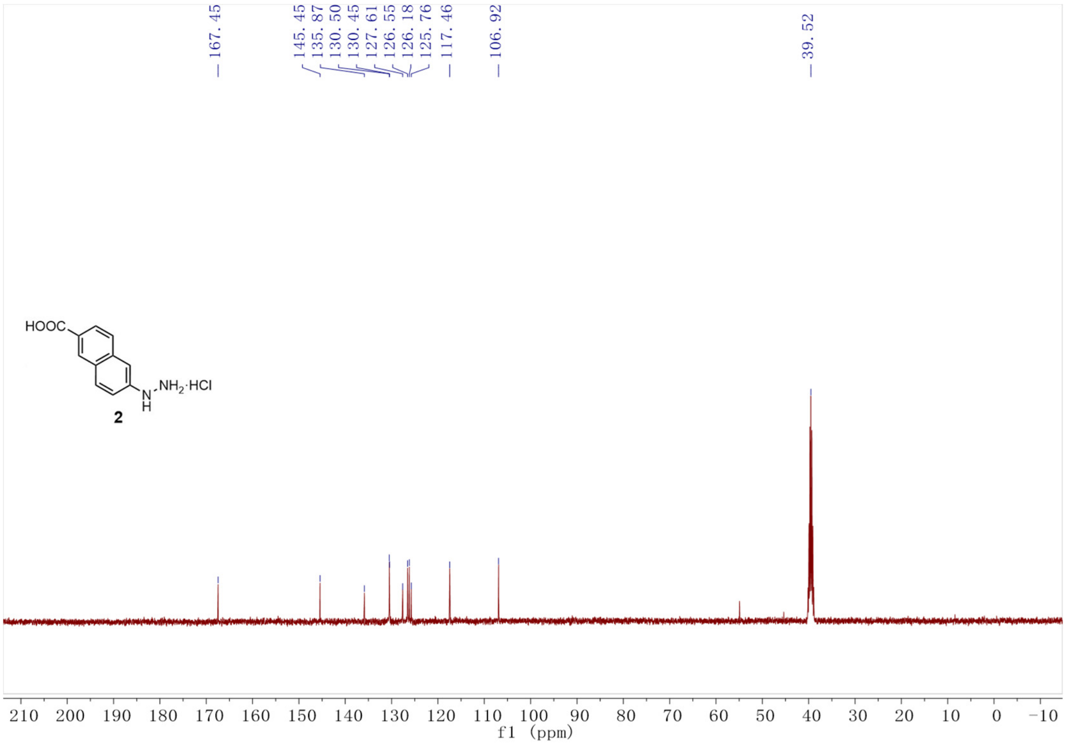

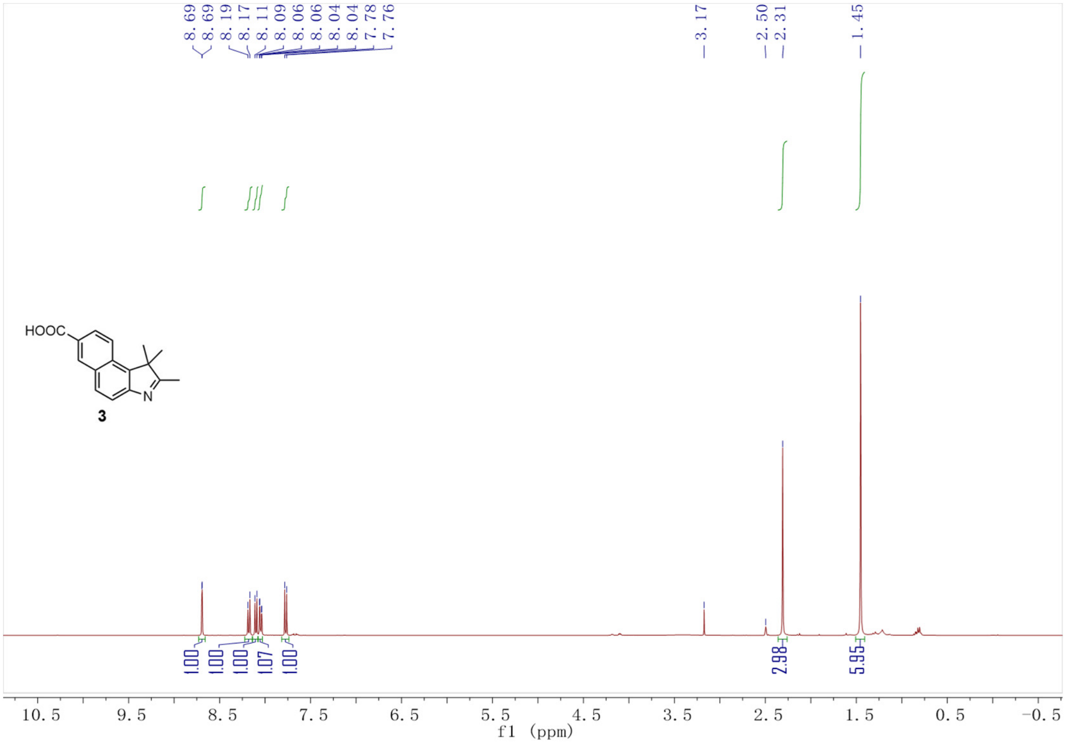
8.69
8.69
8.19
8.17
8.11
8.09
8.06
8.06
8.04
8.04
7.78
7.76
3.17
2.50
2.31
1.45
HOOC
N
3
1.00
1.00
1.00
1.07
1.00
2.98
5.95
10.5
9.5
8.5
7.5
6.5
5.5
4.5
3.5
2.5
1.5
0.5
-0.5
f1 (ppm)

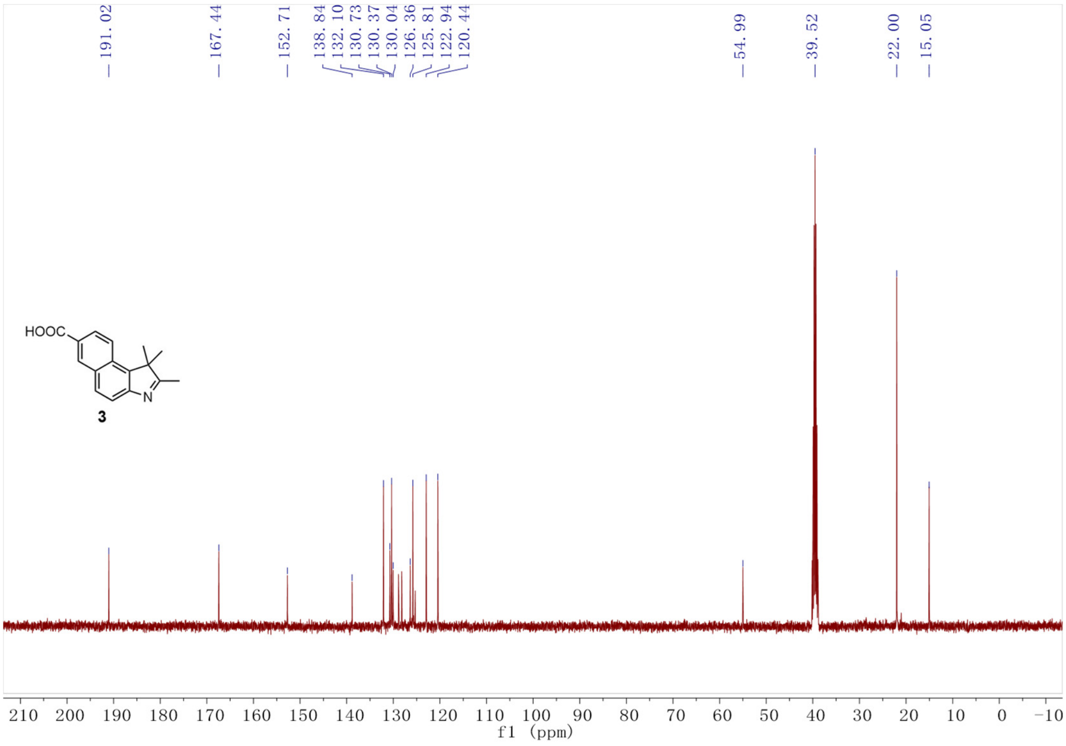
191.02
167.44
152.71
138.84
132.10
130.73
130.37
130.04
126.36
125.81
122.94
120.44
54.99
39.52
22.00
15.05
HOOC
N
3
210
200
190
180
170
160
150
140
130
120
110
100
90
80
70
60
50
40
30
20
10
0
-10
f1 (ppm)

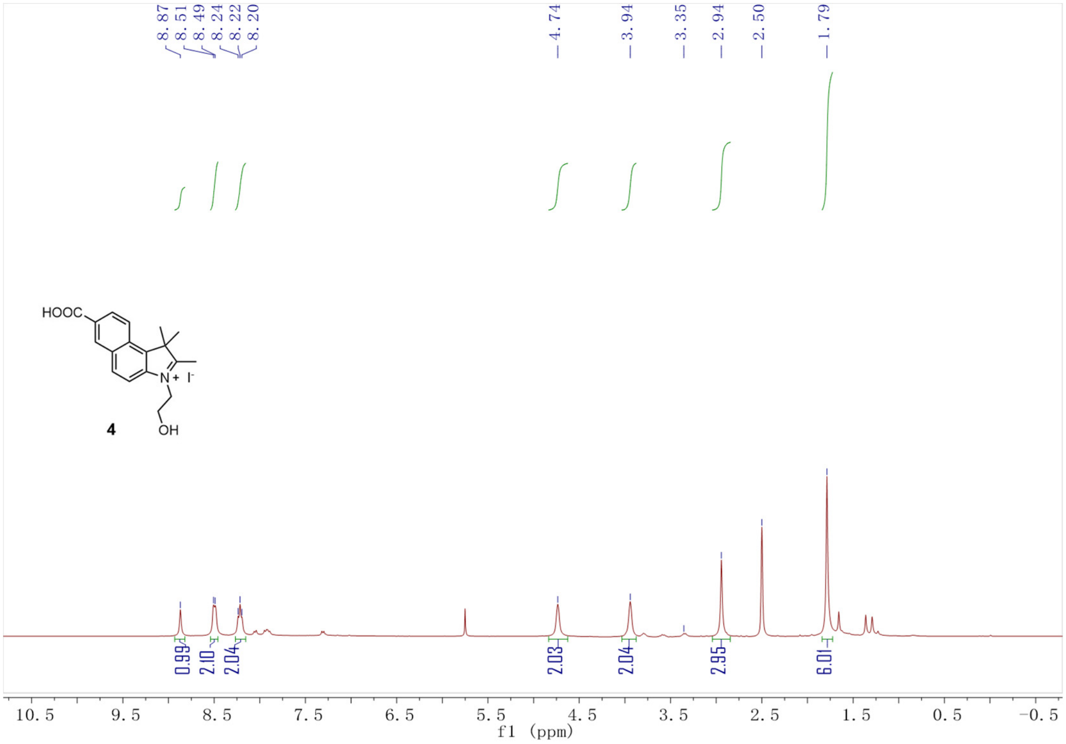
8.87
8.51
8.49
8.24
8.22
8.20
4.74
3.94
3.35
2.94
2.50
1.79
HOOC
N+
I-
4
OH
0.99
2.10
2.04
2.03
2.04
2.95
6.01
10.5 9.5 8.5 7.5 6.5 5.5 4.5 3.5 2.5 1.5 0.5 −0.5
f1 (ppm)

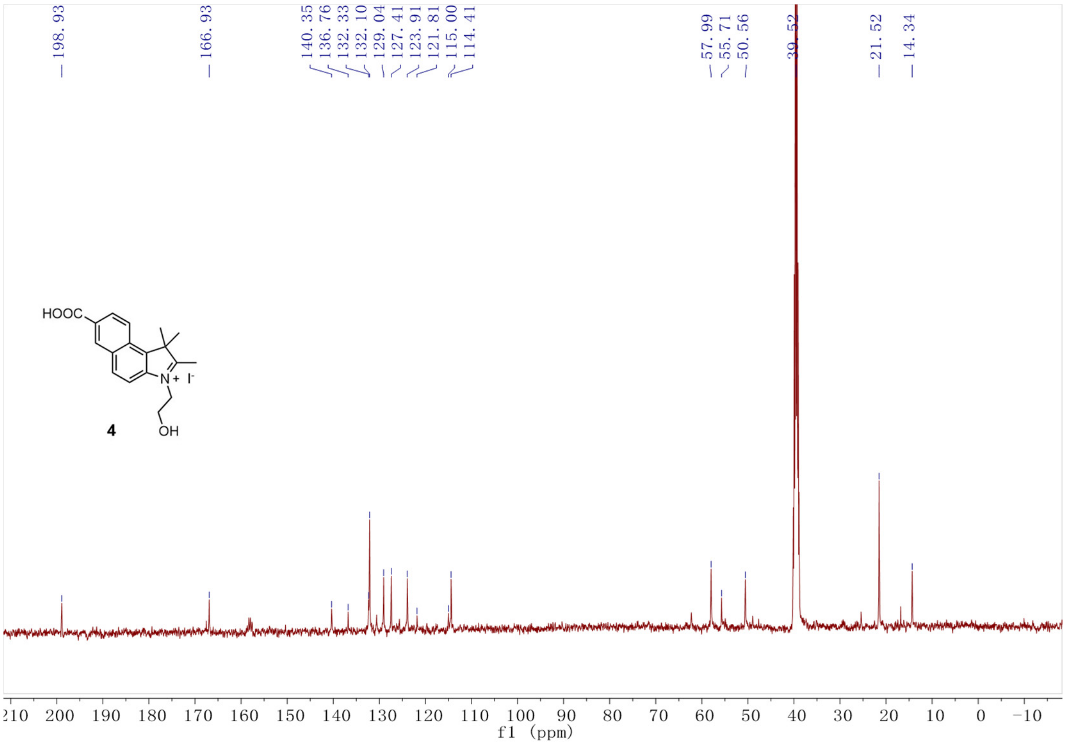
198.93
166.93
140.35
136.76
132.33
132.10
129.04
127.41
123.91
121.81
115.00
114.41
57.99
55.71
50.56
39.52
21.52
14.34
HOOC
N+
I-
4
OH
210 200 190 180 170 160 150 140 130 120 110 100 90 80 70 60 50 40 30 20 10 0 −10
f1 (ppm)

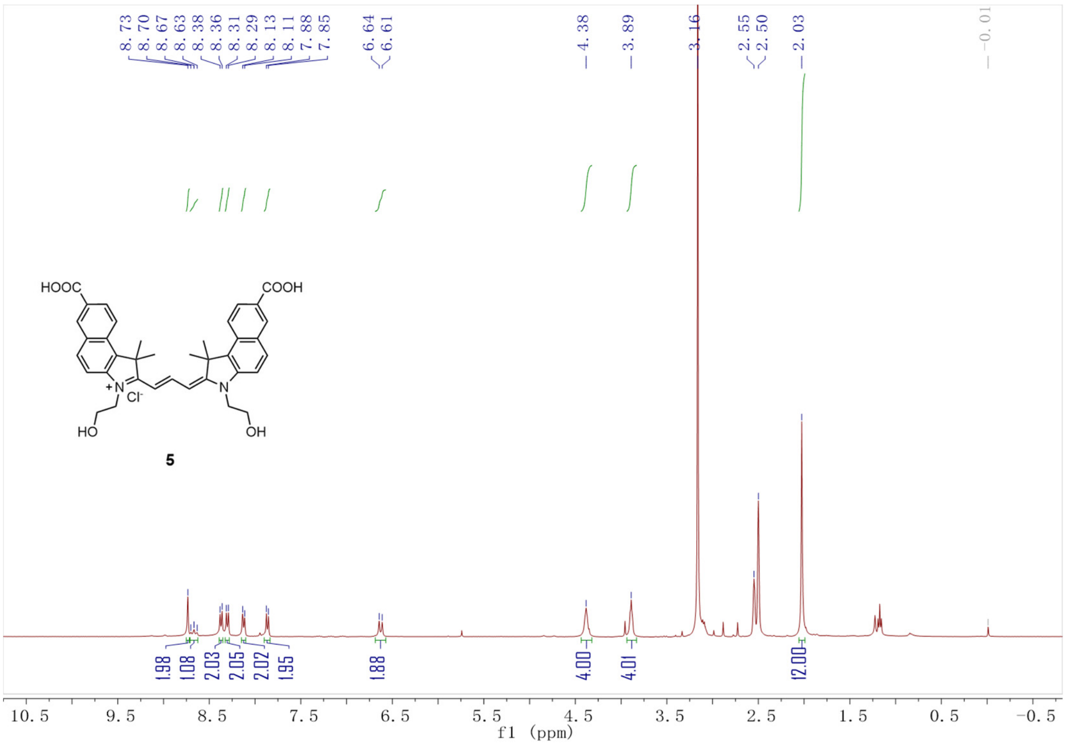
8.73
8.70
8.67
8.63
8.38
8.36
8.31
8.29
8.13
8.11
7.88
7.85
6.64
6.61
4.38
3.89
3.16
2.55
2.50
2.03
-0.01
HOOC
COOH
Cl-
HO
OH
5
1.98
1.08
2.03
2.05
2.02
1.95
1.88
4.00
4.01
12.00
10.5
9.5
8.5
7.5
6.5
5.5
4.5
3.5
2.5
1.5
0.5
-0.5
f1 (ppm)

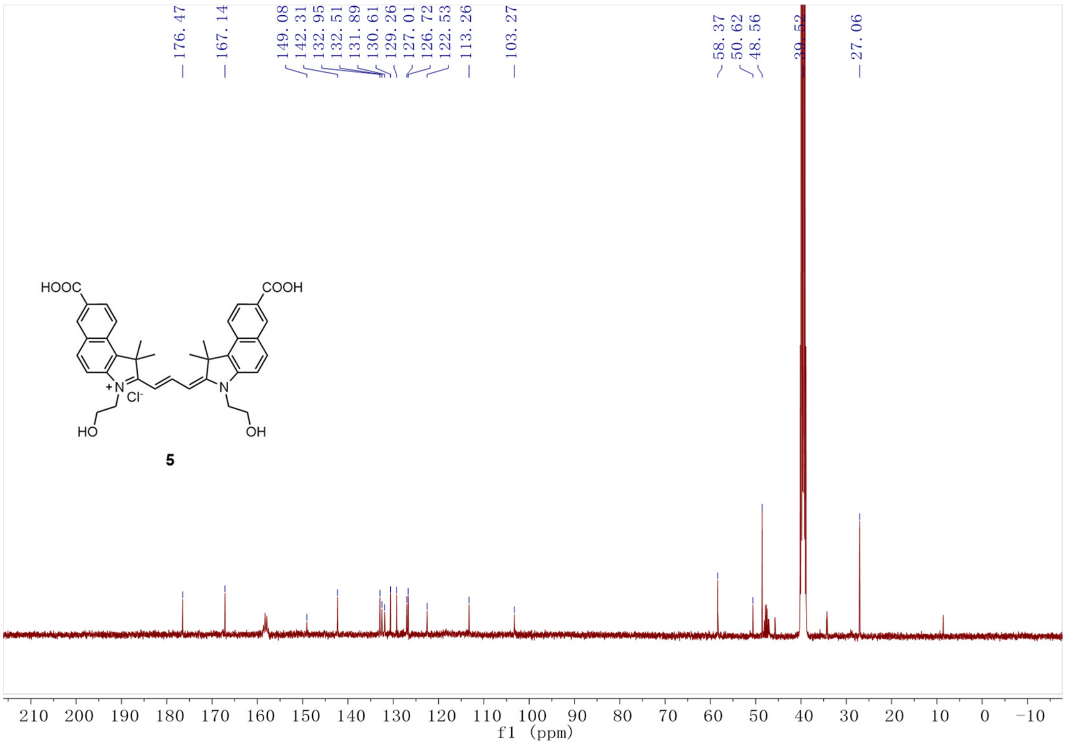
176.47
167.14
149.08
142.31
132.95
132.51
131.89
130.61
129.26
127.01
126.72
122.53
113.26
103.27
58.37
50.62
48.56
27.06
HOOC
COOH
Cl-
HO
OH
5
210
200
190
180
170
160
150
140
130
120
110
100
90
80
70
60
50
40
30
20
10
0
-10
f1 (ppm)

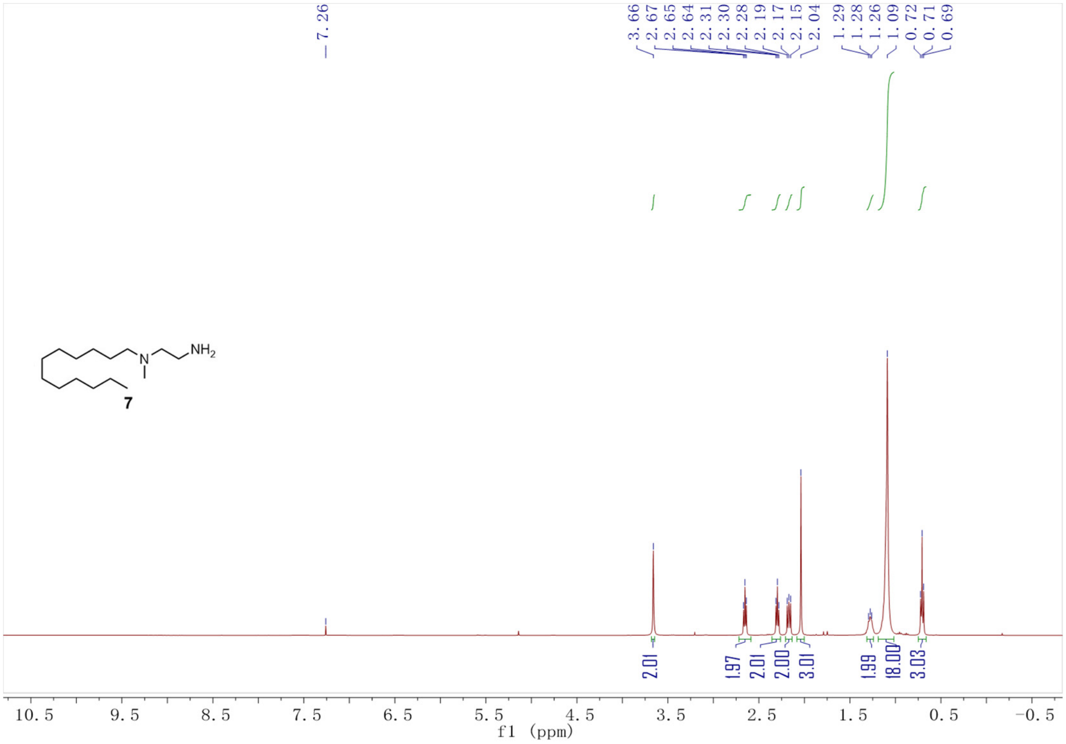
7.26
3.66
2.67
2.65
2.64
2.31
2.30
2.28
2.19
2.17
2.15
2.04
1.29
1.28
1.26
1.09
0.72
0.71
0.69
NH2
N
7
2.01
1.97
2.01
2.00
3.01
1.99
18.00
3.03
10.5 9.5 8.5 7.5 6.5 5.5 4.5 3.5 2.5 1.5 0.5 −0.5
f1 (ppm)

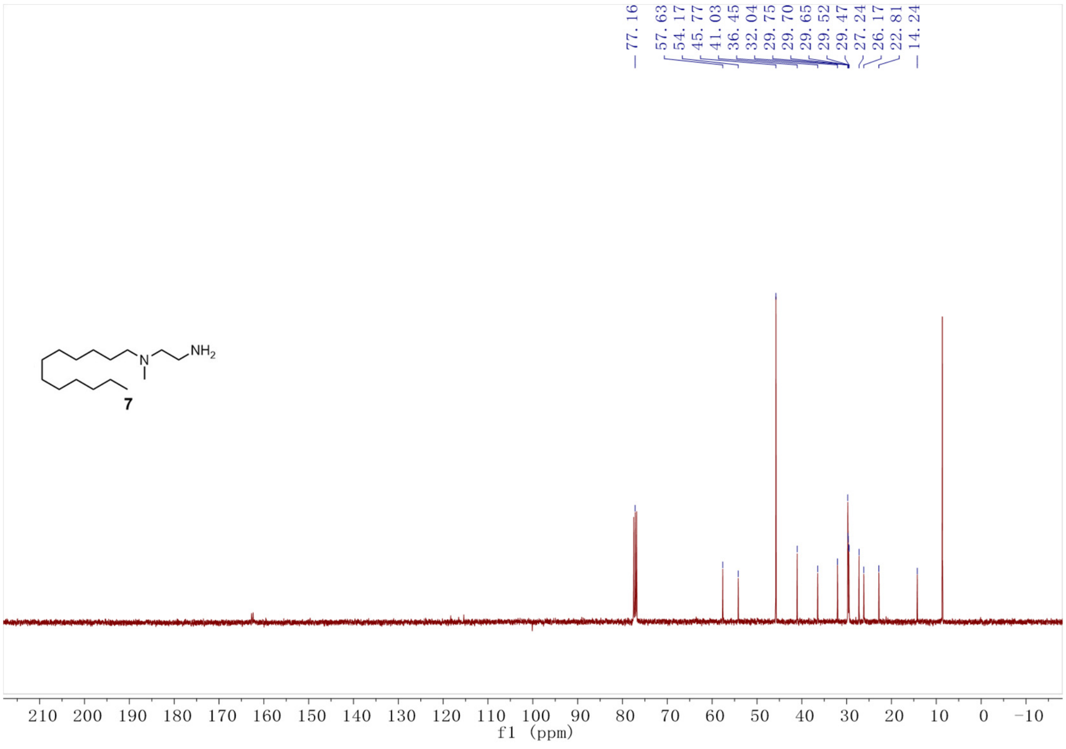
77.16
57.63
54.17
45.77
41.03
36.45
32.04
29.75
29.70
29.65
29.52
29.47
27.24
26.17
22.81
14.24
NH2
N
7
210 200 190 180 170 160 150 140 130 120 110 100 90 80 70 60 50 40 30 20 10 0 −10
f1 (ppm)

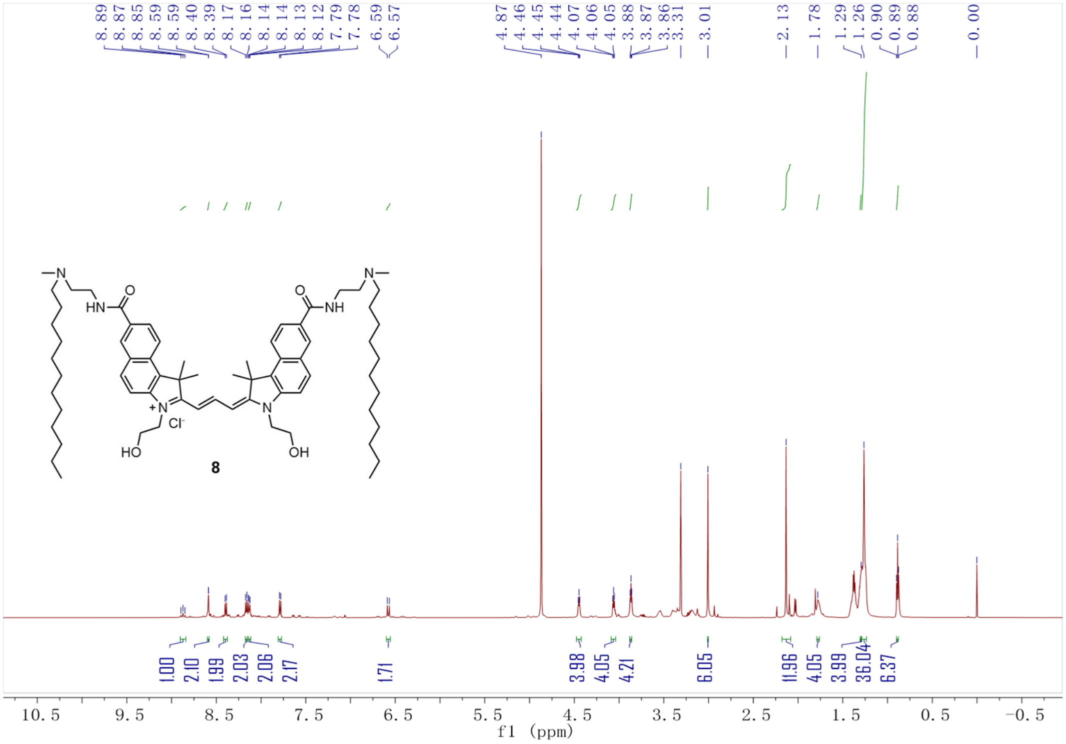

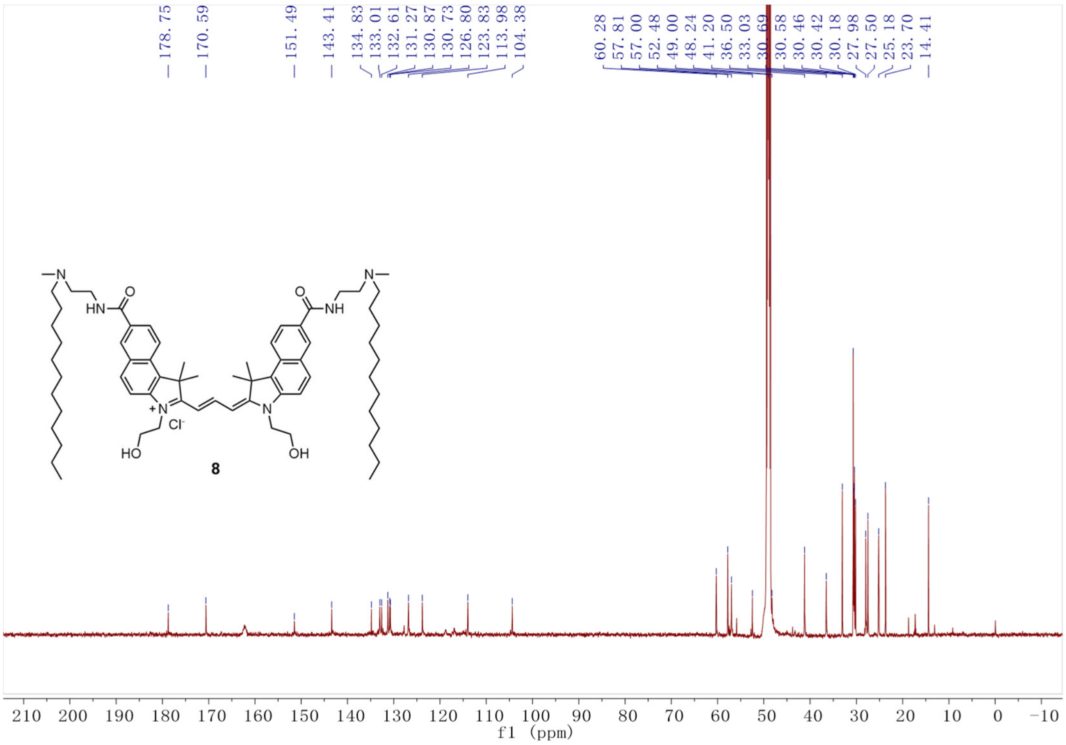

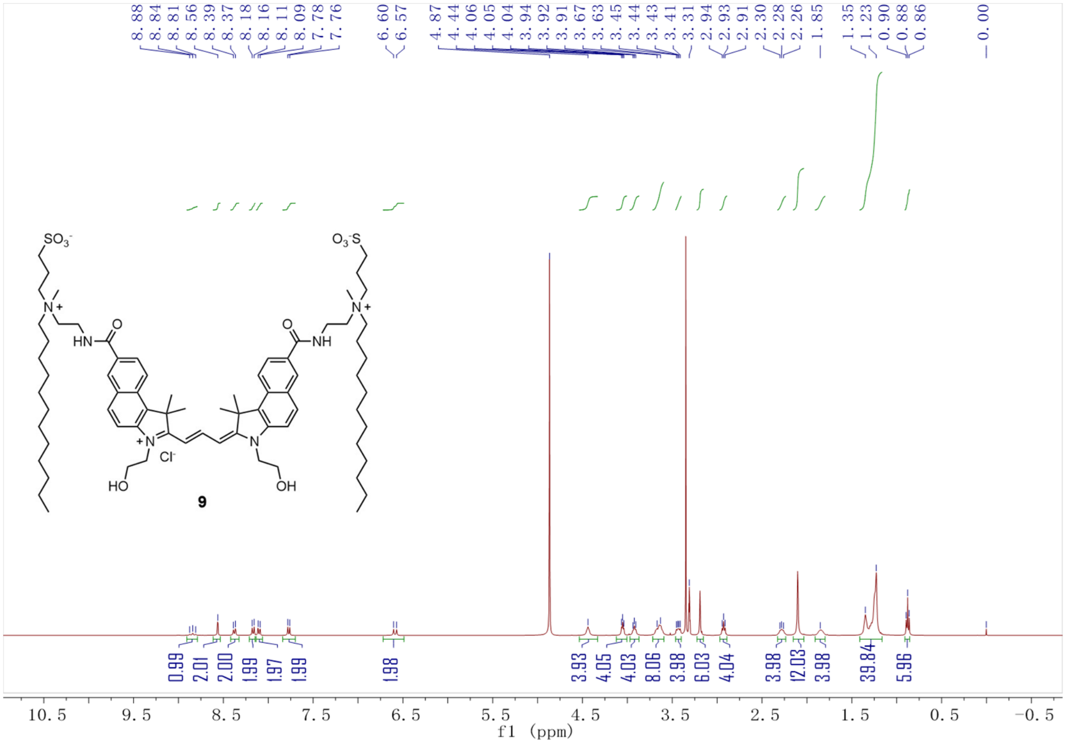
f1 (ppm)

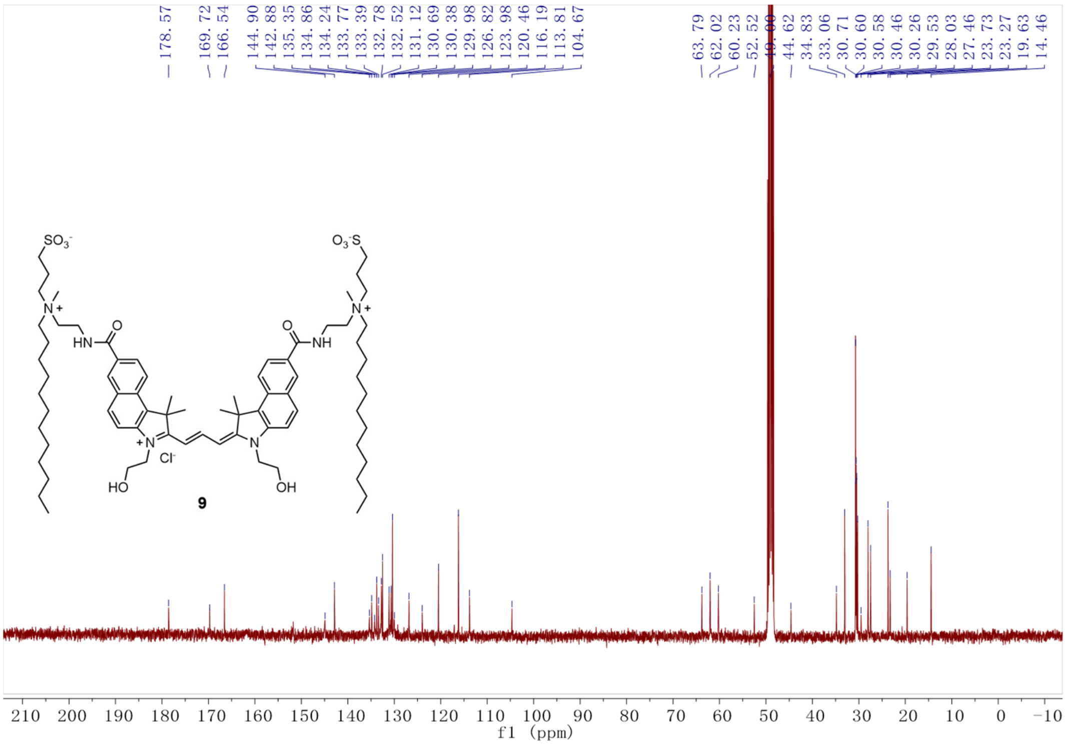
f1 (ppm)

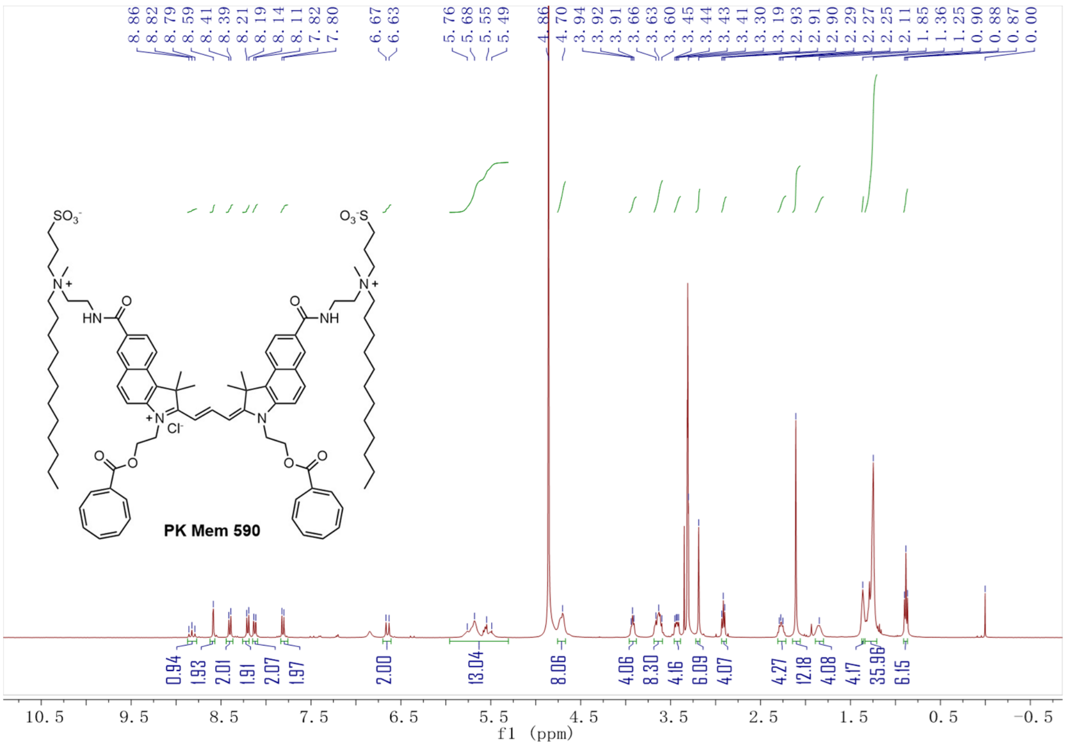

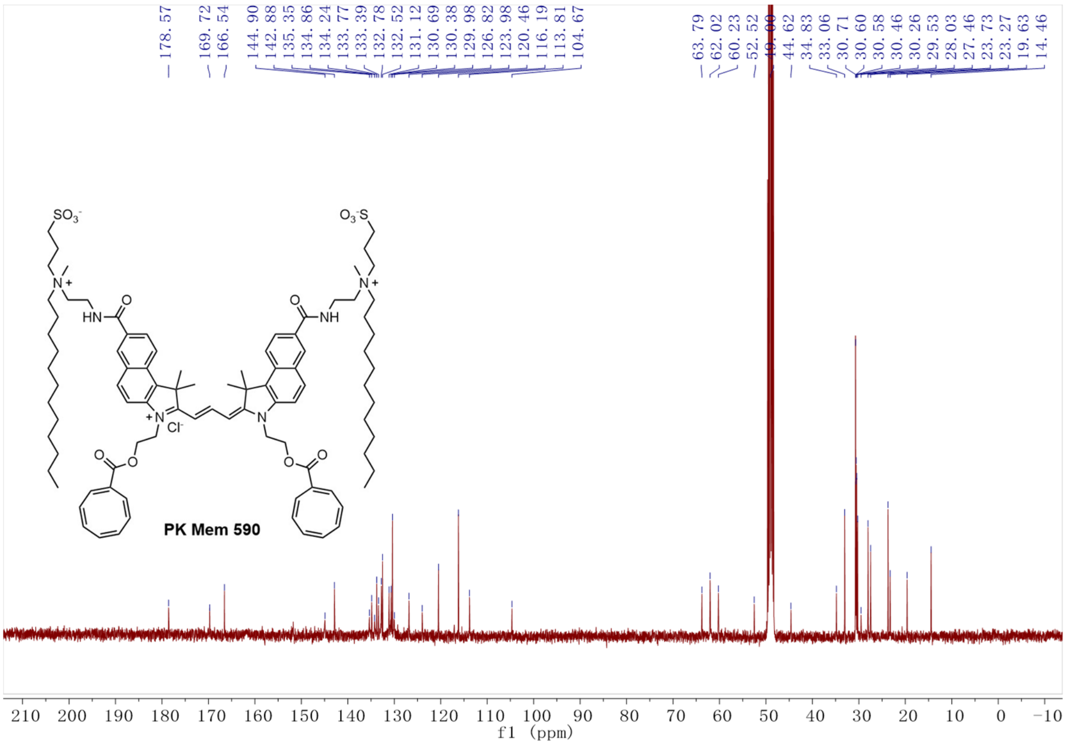

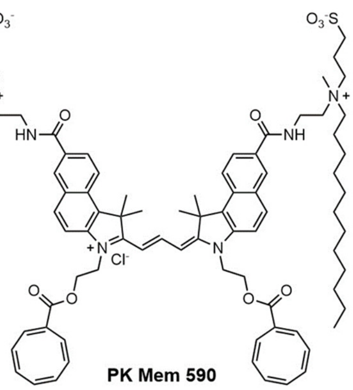
SO3-
O3-S
N+
HN
NH
N
Cl-
O

**PK Mem 590**

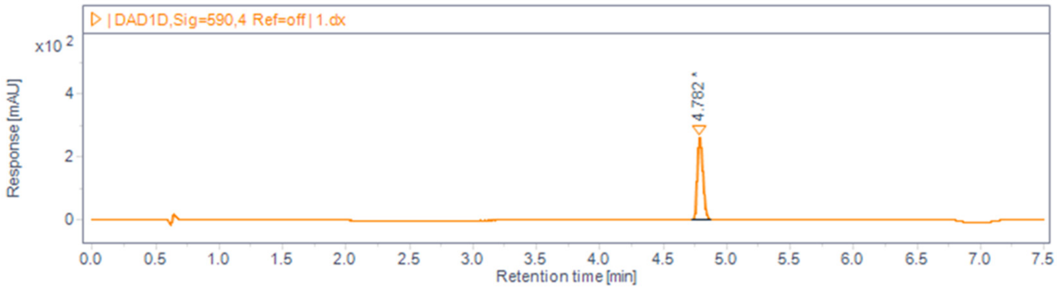
| DAD1D, Sig=590,4 Ref=off | 1.dx
x10 2
Response [mAU]
4.782
Retention time [min]

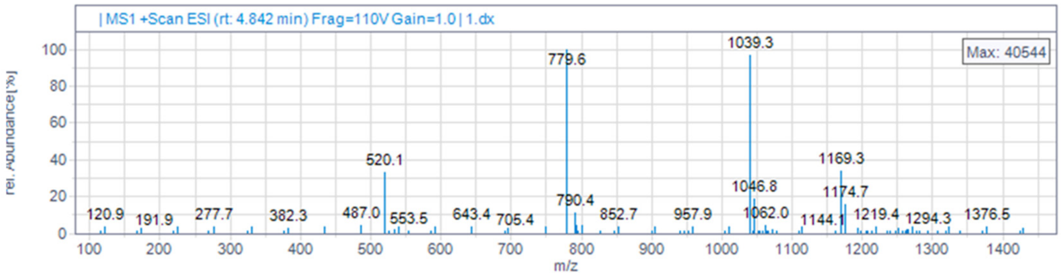
| MS1 +Scan ESI (rt: 4.842 min) Frag=110V Gain=1.0 | 1.dx
Max: 40544
rel. Abundance [%]
779.6
1039.3
520.1
1169.3
1046.8
1174.7
790.4
m/z

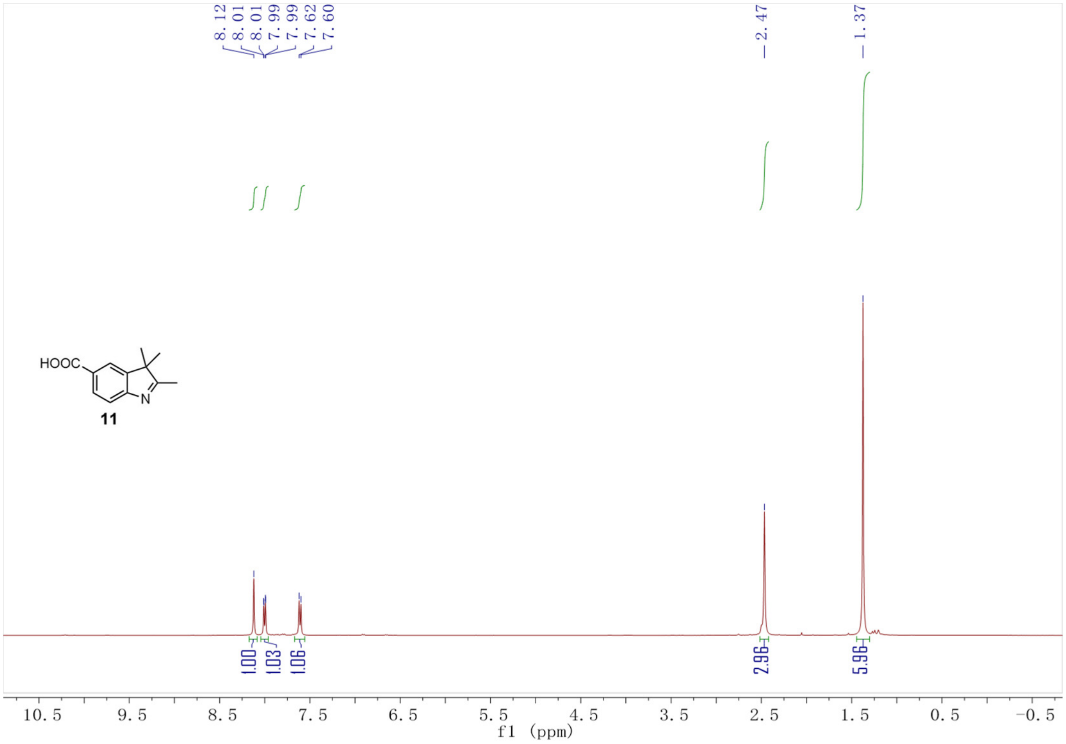
8.12
8.01
8.01
7.99
7.99
7.62
7.60
2.47
1.37
HOOC
N
11
1.00
1.03
1.06
2.96
5.96
10.5 9.5 8.5 7.5 6.5 5.5 4.5 3.5 2.5 1.5 0.5 -0.5
f1 (ppm)

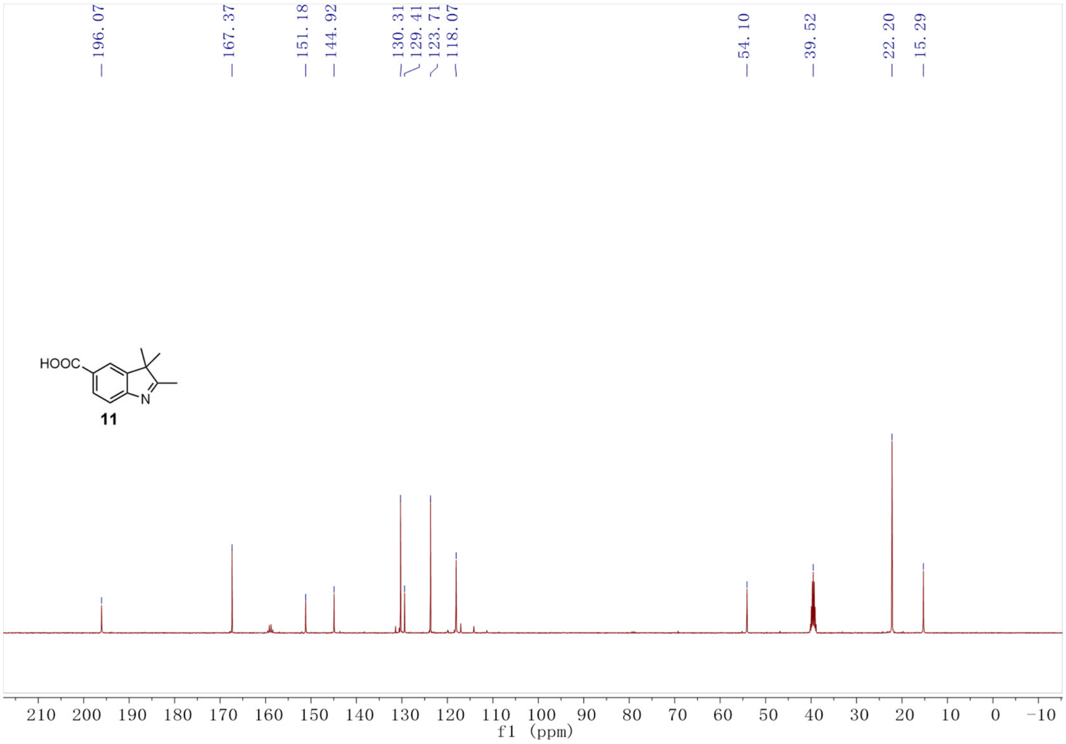
196.07
167.37
151.18
144.92
130.31
129.41
123.71
118.07
54.10
39.52
22.20
15.29
HOOC
N
11
210 200 190 180 170 160 150 140 130 120 110 100 90 80 70 60 50 40 30 20 10 0 -10
f1 (ppm)

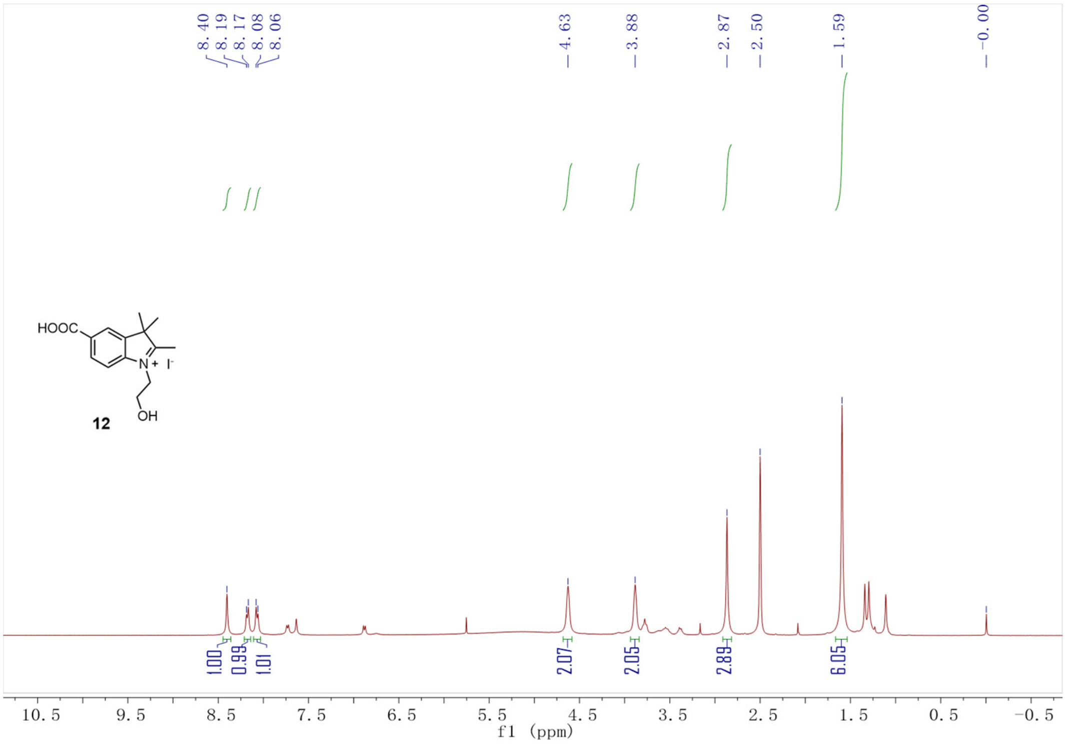
8.40
8.19
8.17
8.08
8.06
4.63
3.88
2.87
2.50
1.59
-0.00
HOOC
N+
I-
OH
12
1.00
0.99
1.01
2.07
2.05
2.89
6.05
10.5 9.5 8.5 7.5 6.5 5.5 4.5 3.5 2.5 1.5 0.5 -0.5
f1 (ppm)

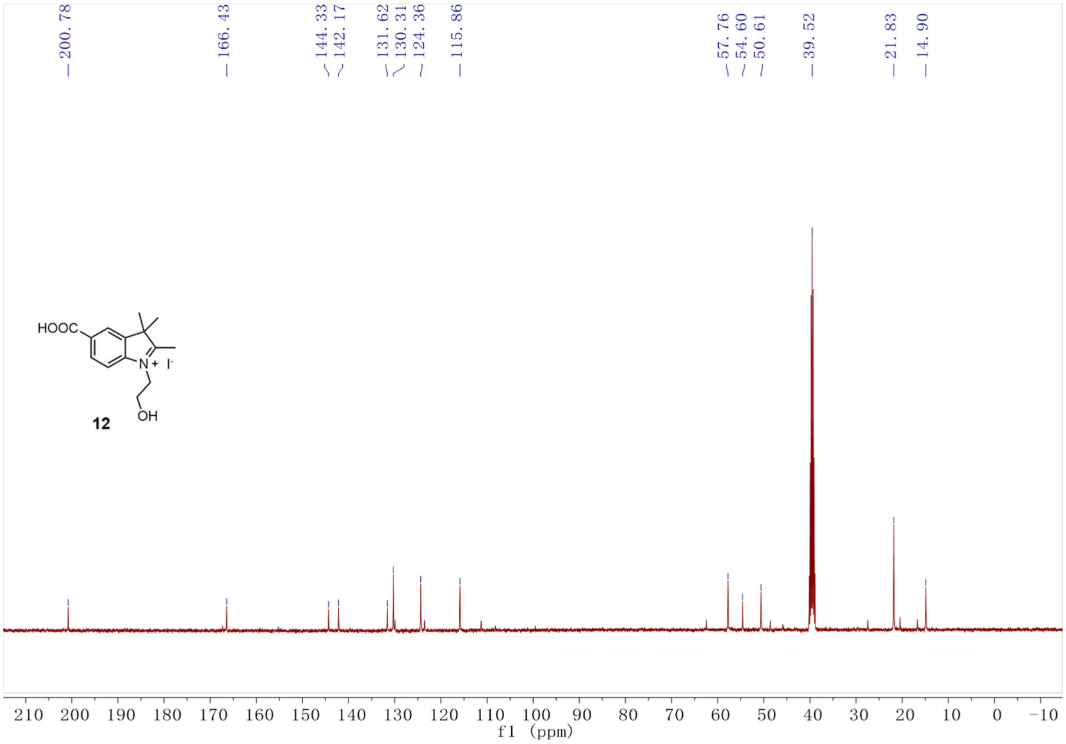
200.78
166.43
144.33
142.17
131.62
130.31
124.36
115.86
57.76
54.60
50.61
39.52
21.83
14.90
HOOC
N+
I-
OH
12
210 200 190 180 170 160 150 140 130 120 110 100 90 80 70 60 50 40 30 20 10 0 -10
f1 (ppm)

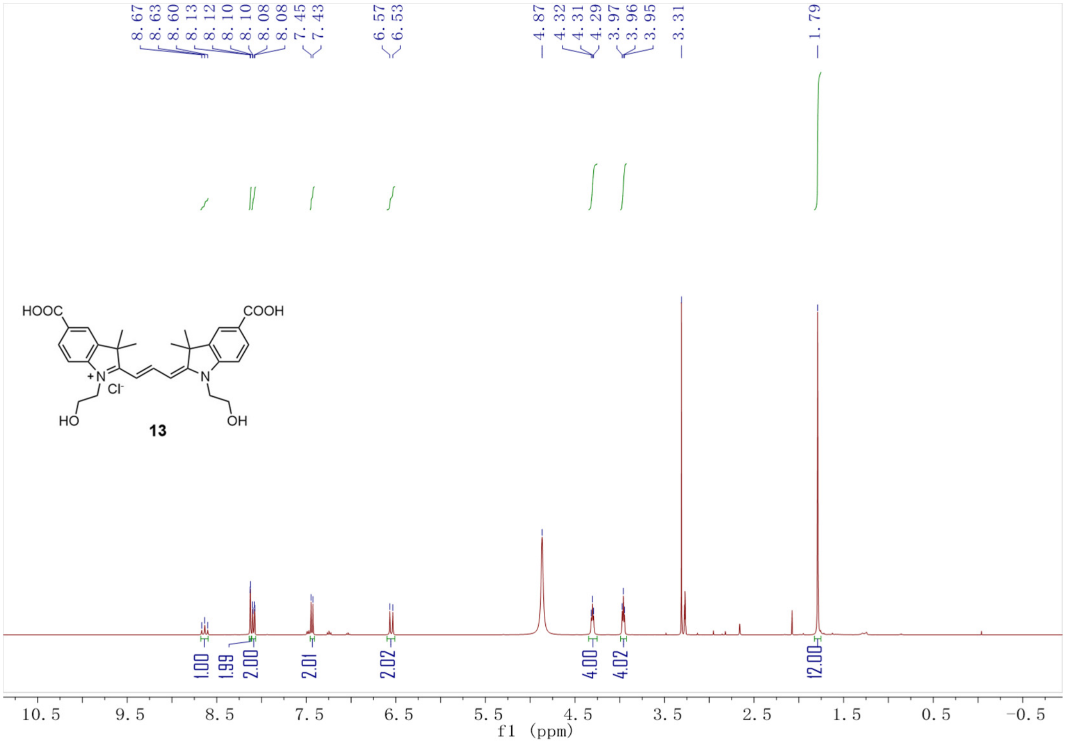
8.67
8.63
8.60
8.13
8.12
8.10
8.10
8.08
8.08
7.45
7.43
6.57
6.53
4.87
4.32
4.31
4.29
3.97
3.96
3.95
3.31
1.79
HOOC
COOH
Cl-
HO
OH
13
1.00
1.99
2.00
2.01
2.02
4.00
4.02
12.00
10.5
9.5
8.5
7.5
6.5
5.5
4.5
3.5
2.5
1.5
0.5
-0.5
f1 (ppm)

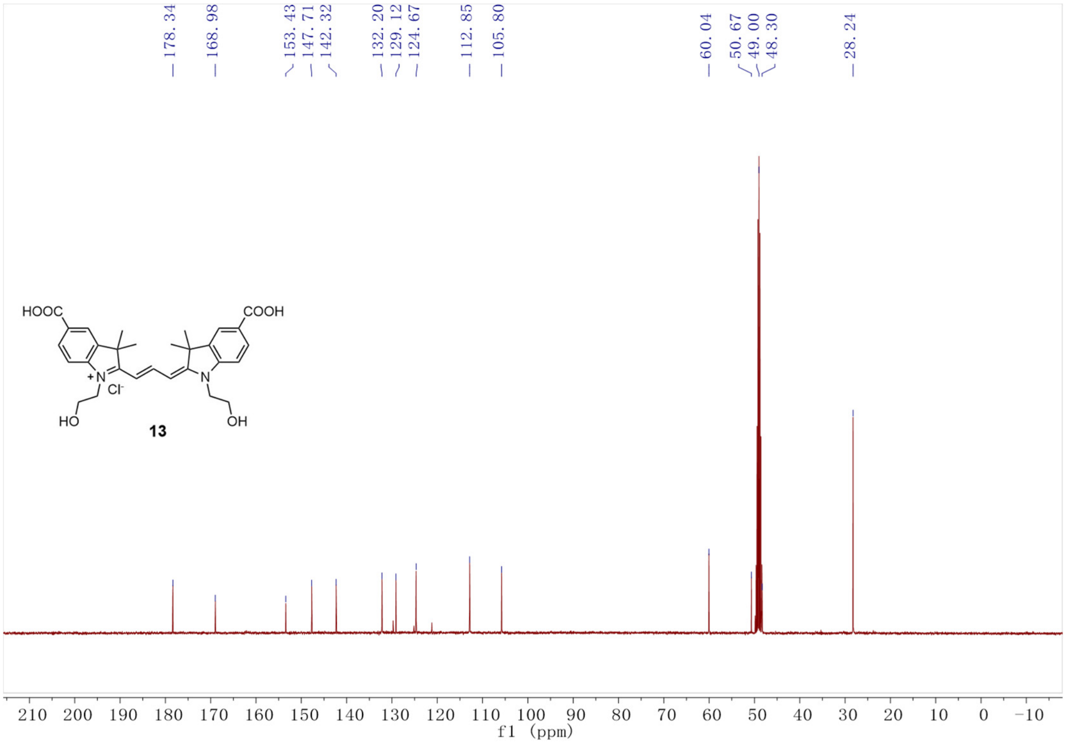
178.34
168.98
153.43
147.71
142.32
132.20
129.12
124.67
112.85
105.80
60.04
50.67
49.00
48.30
28.24
HOOC
COOH
Cl-
HO
OH
13
210
200
190
180
170
160
150
140
130
120
110
100
90
80
70
60
50
40
30
20
10
0
-10
f1 (ppm)

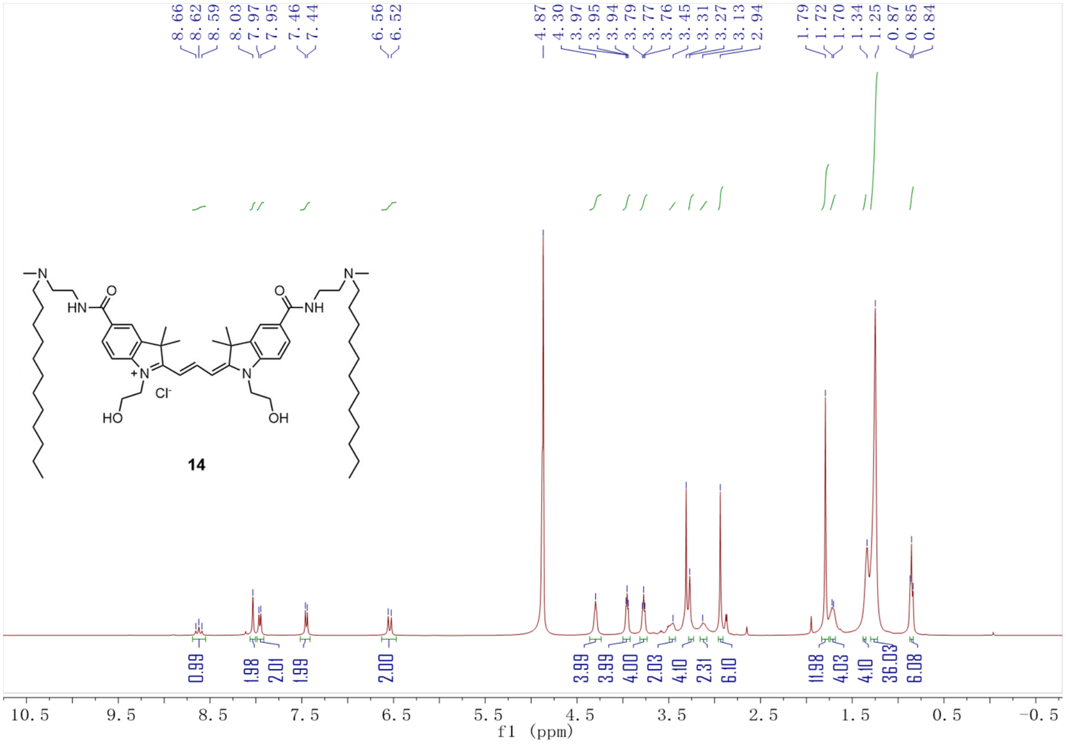
8.66
8.62
8.59
8.03
7.97
7.95
7.46
7.44
6.56
6.52
4.87
4.30
3.97
3.95
3.94
3.79
3.77
3.76
3.45
3.31
3.27
3.13
2.94
1.79
1.72
1.70
1.34
1.25
0.87
0.85
0.84
HN
NH
O
O
N
N
Cl⁻
HO
OH
14
0.99
1.98
2.01
1.99
2.00
3.99
3.99
4.00
2.03
4.10
2.31
6.10
11.98
4.03
4.10
36.03
6.08
10.5
9.5
8.5
7.5
6.5
5.5
4.5
3.5
2.5
1.5
0.5
-0.5
f1 (ppm)

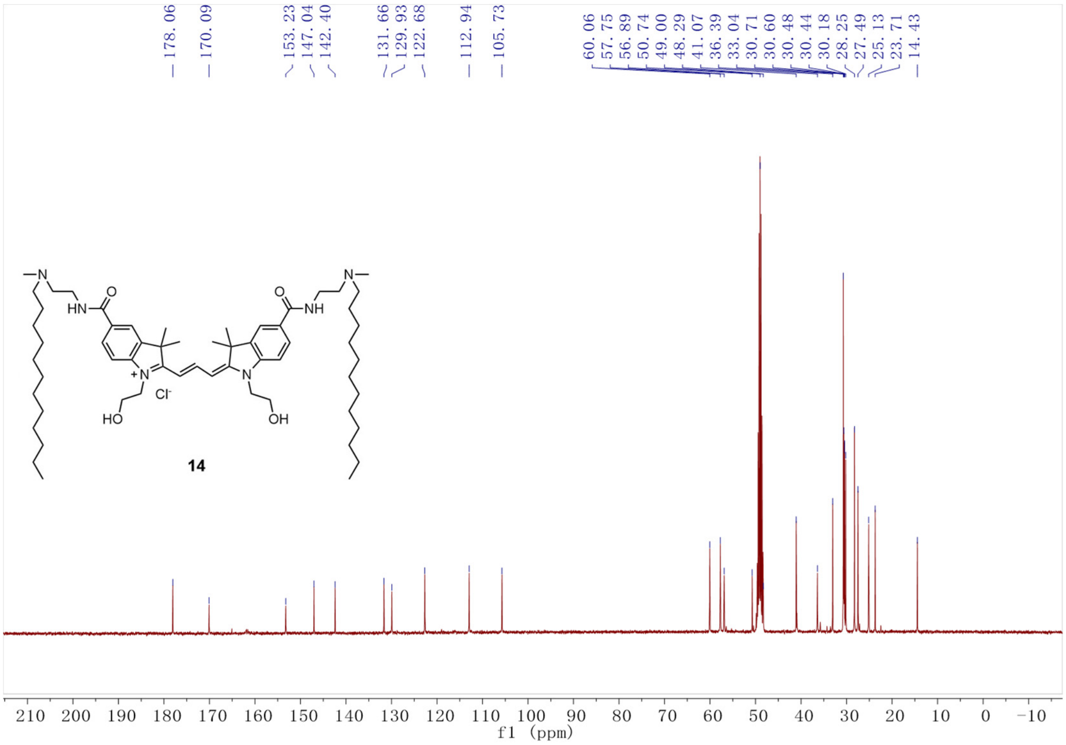
178.06
170.09
153.23
147.04
142.40
131.66
129.93
122.68
112.94
105.73
60.06
57.75
56.89
50.74
49.00
48.29
41.07
36.39
33.04
30.71
30.60
30.48
30.44
30.18
28.25
27.49
25.13
23.71
14.43
HN
NH
O
O
N
N
Cl⁻
HO
OH
14
210
200
190
180
170
160
150
140
130
120
110
100
90
80
70
60
50
40
30
20
10
0
-10
f1 (ppm)

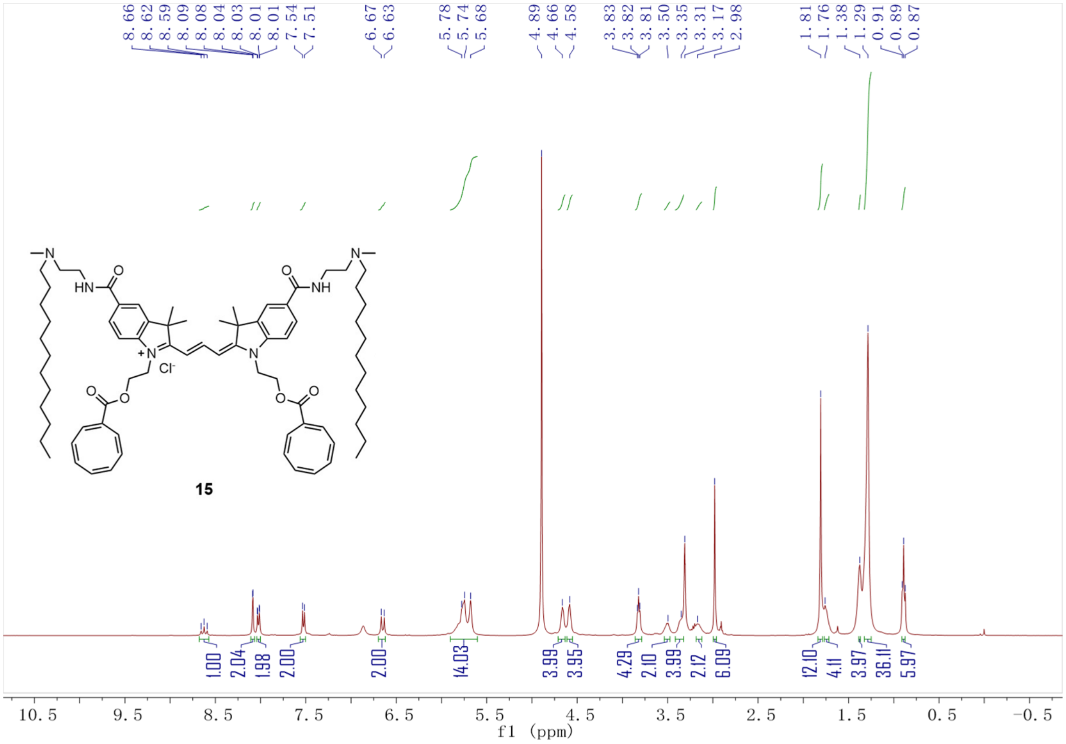
8.66 8.62 8.59 8.09 8.08 8.04 8.03 8.01 8.01 7.54 7.51
6.67 6.63
5.78 5.74 5.68
4.89 4.66 4.58
3.83 3.82 3.81 3.50 3.35 3.31 3.17 2.98
1.81 1.76 1.38 1.29 0.91 0.89 0.87
HN
O
NH
N
Cl⁻
O
15
1.00 2.04 1.98 2.00 2.00 14.03 3.99 3.95 4.29 2.10 3.99 2.12 6.09 12.10 4.11 3.97 36.11 5.97
10.5 9.5 8.5 7.5 6.5 5.5 4.5 3.5 2.5 1.5 0.5 -0.5
f1 (ppm)

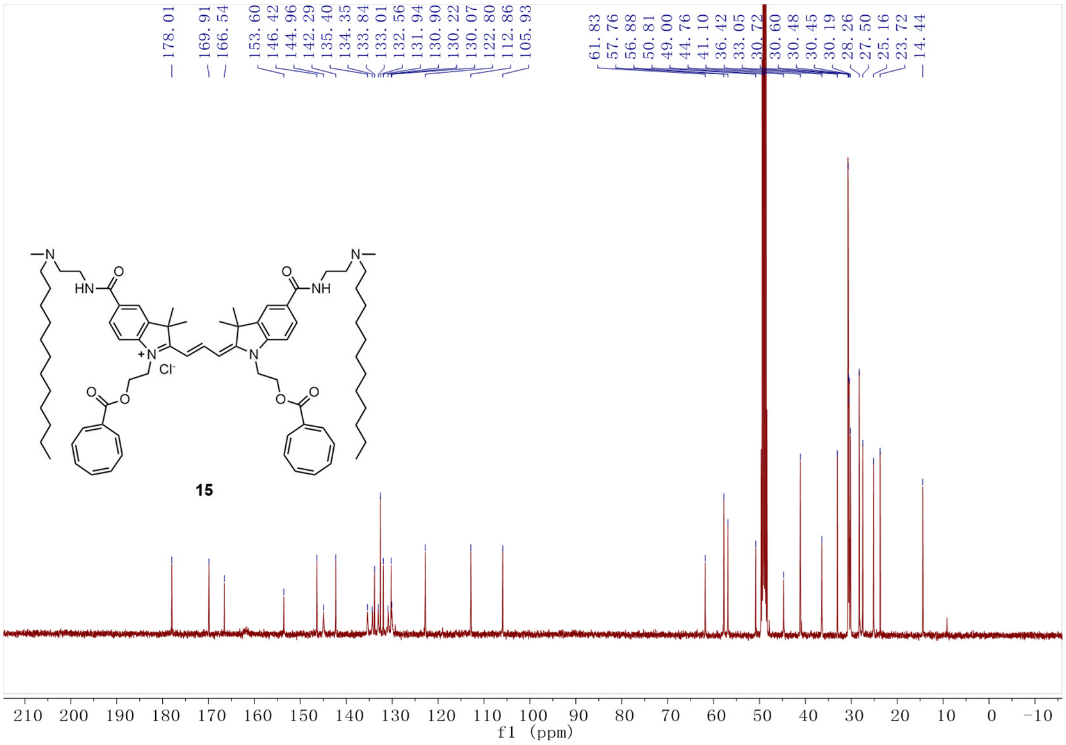
178.01
169.91 166.54
153.60 146.42 144.96 142.29 135.40 134.35 133.84 133.01 132.56 131.94 130.90 130.22 130.07 122.80 112.86 105.93
61.83 57.76 56.88 50.81 49.00 44.76 41.10 36.42 33.05 30.72 30.60 30.48 30.45 30.19 28.26 27.50 25.16 23.72 14.44
HN
O
NH
N
Cl⁻
O
15
210 200 190 180 170 160 150 140 130 120 110 100 90 80 70 60 50 40 30 20 10 0 -10
f1 (ppm)

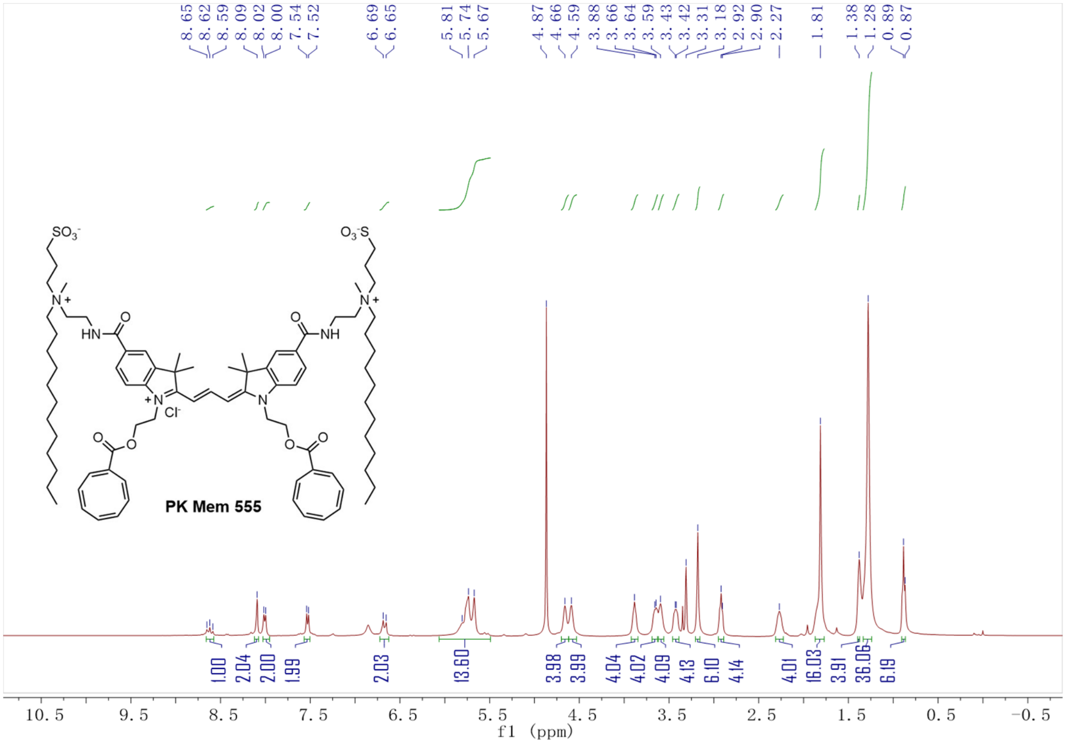

8.65 8.62 8.59 8.09 8.02 8.00 7.54 7.52 6.69 6.65 5.81 5.74 5.67 4.87 4.66 4.59 3.88 3.66 3.64 3.59 3.43 3.42 3.31 3.18 2.92 2.90 2.27 1.81 1.38 1.28 0.89 0.87
PK Mem 555
1.00 2.04 2.00 1.99 2.03 13.60 3.98 3.99 4.04 4.02 4.09 4.13 6.10 4.14 4.01 16.03 3.91 36.06 6.19
10.5 9.5 8.5 7.5 6.5 5.5 4.5 3.5 2.5 1.5 0.5 -0.5
f1 (ppm)

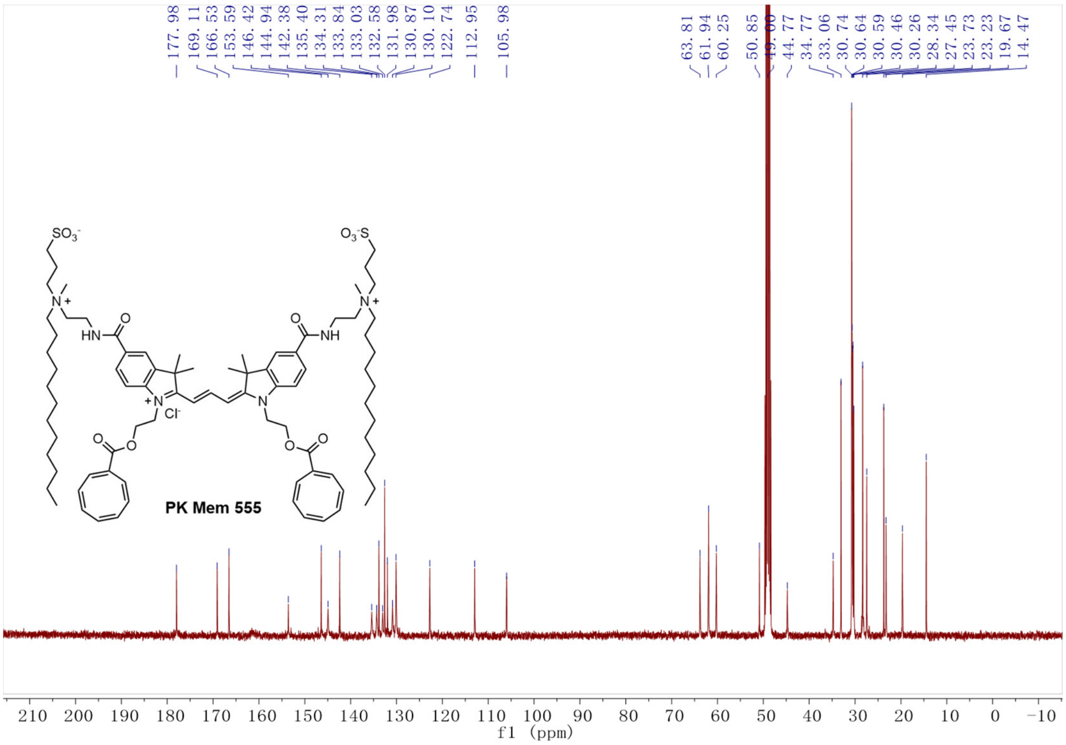

177.98 169.11 166.53 153.59 146.42 144.94 142.38 135.40 134.31 133.84 133.03 132.58 131.98 130.87 130.10 122.74 112.95 105.98
63.81 61.94 60.25 50.85 49.00 44.77 34.77 33.06 30.74 30.64 30.59 30.46 30.26 28.34 27.45 23.73 23.23 19.67 14.47
PK Mem 555
210 200 190 180 170 160 150 140 130 120 110 100 90 80 70 60 50 40 30 20 10 0 -10
f1 (ppm)

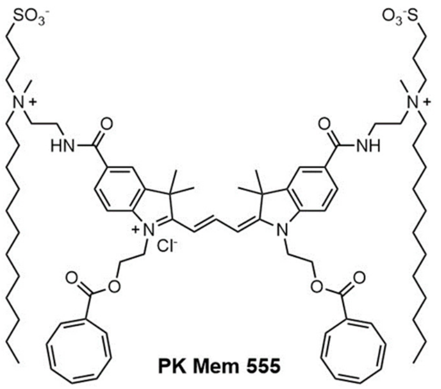

PK Mem 555

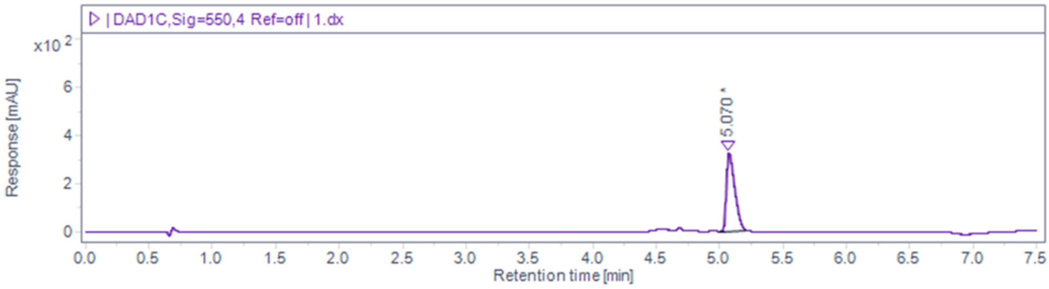

| DAD1C,Sig=550,4 Ref=off | 1.dx
x10 2
Response [mAU]
5.070
Retention time [min]

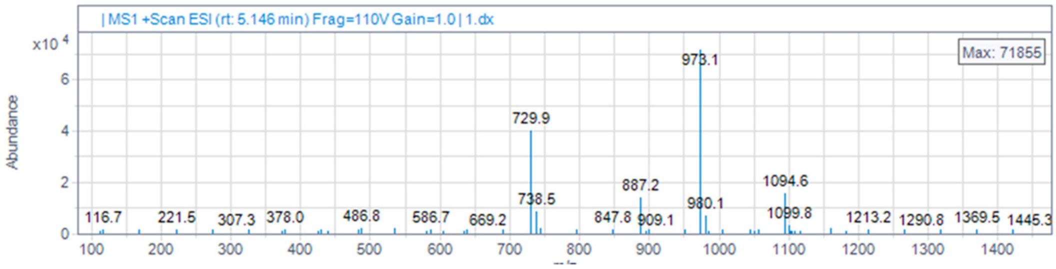

| MS1 +Scan ESI (rt: 5.146 min) Frag=110V Gain=1.0 | 1.dx
x10 4
Abundance
Max: 71855
973.1
729.9
887.2
1094.6
738.5
980.1
1099.8
116.7
221.5
307.3
378.0
486.8
586.7
669.2
847.8
909.1
1213.2
1290.8
1369.5
1445.3

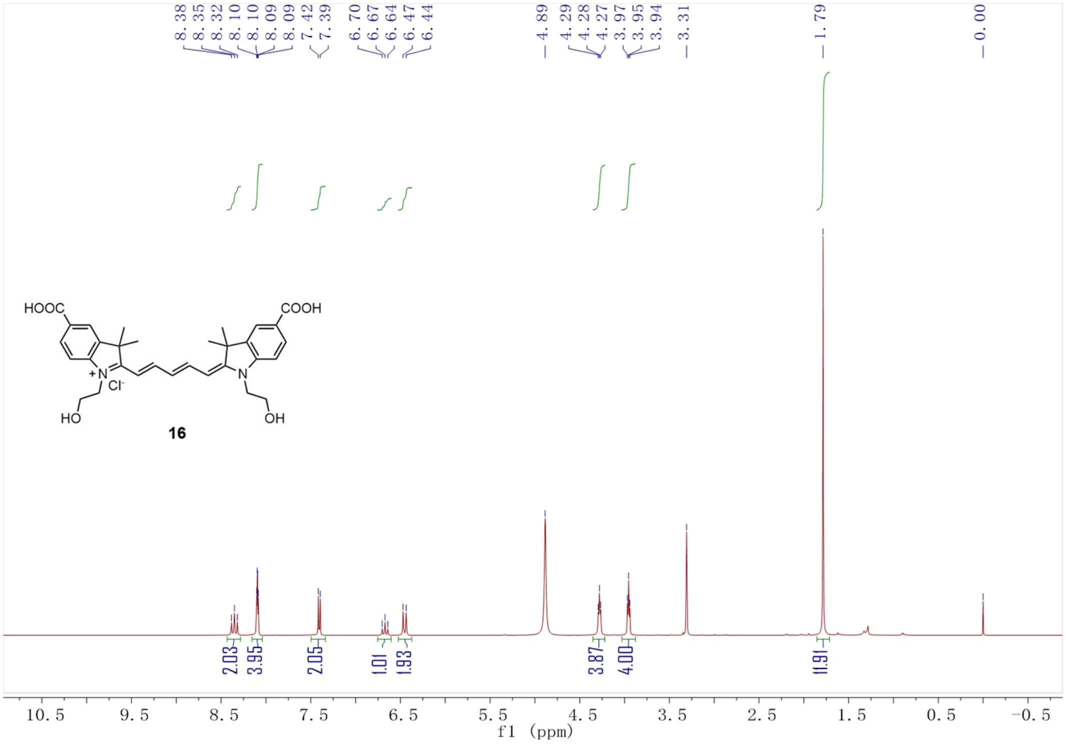
8.38
8.35
8.32
8.10
8.10
8.09
8.09
7.42
7.39
6.70
6.67
6.64
6.47
6.44
4.89
4.29
4.28
4.27
3.97
3.95
3.94
3.31
1.79
0.00
HOOC
COOH
Cl-
HO
OH
16
2.03
3.95
2.05
1.01
1.93
3.87
4.00
11.91
10.5 9.5 8.5 7.5 6.5 5.5 4.5 3.5 2.5 1.5 0.5 -0.5
f1 (ppm)

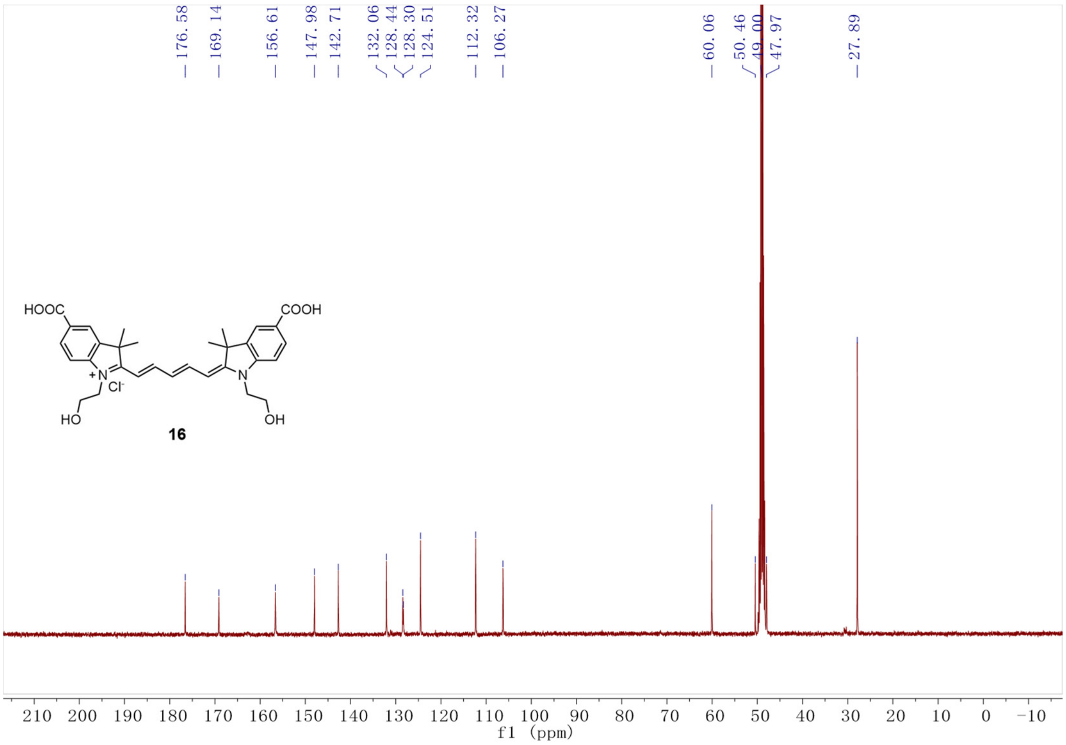
176.58
169.14
156.61
147.98
142.71
132.06
128.44
128.30
124.51
112.32
106.27
60.06
50.46
49.00
47.97
27.89
HOOC
COOH
Cl-
HO
OH
16
210 200 190 180 170 160 150 140 130 120 110 100 90 80 70 60 50 40 30 20 10 0 -10
f1 (ppm)

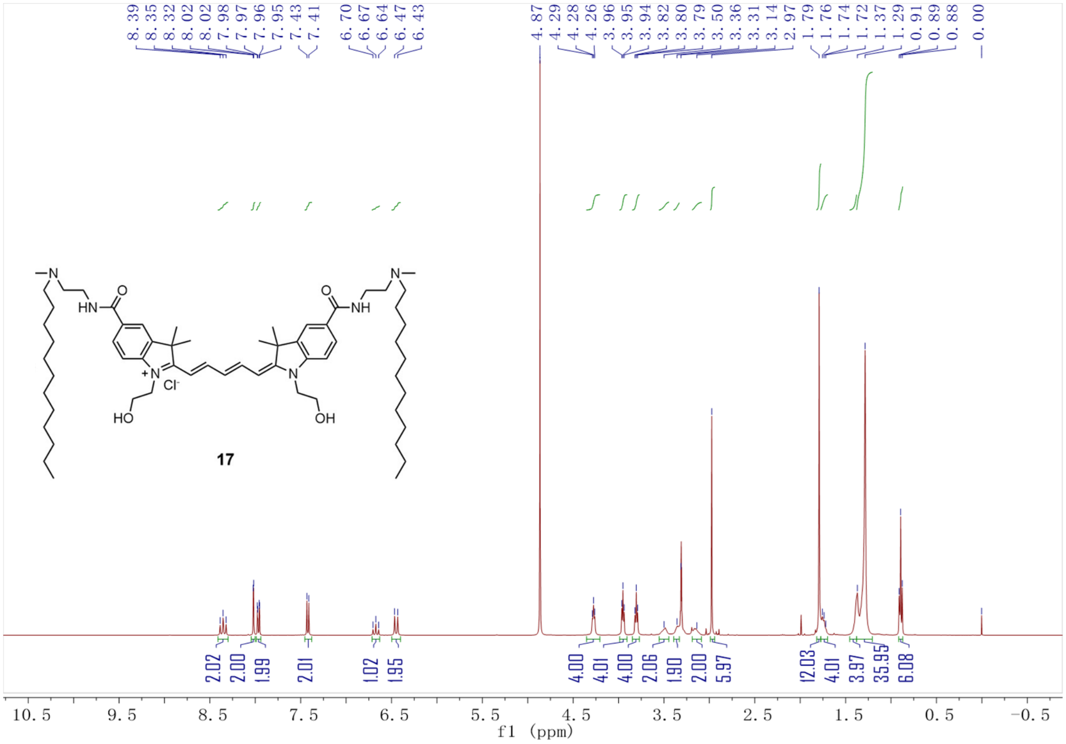

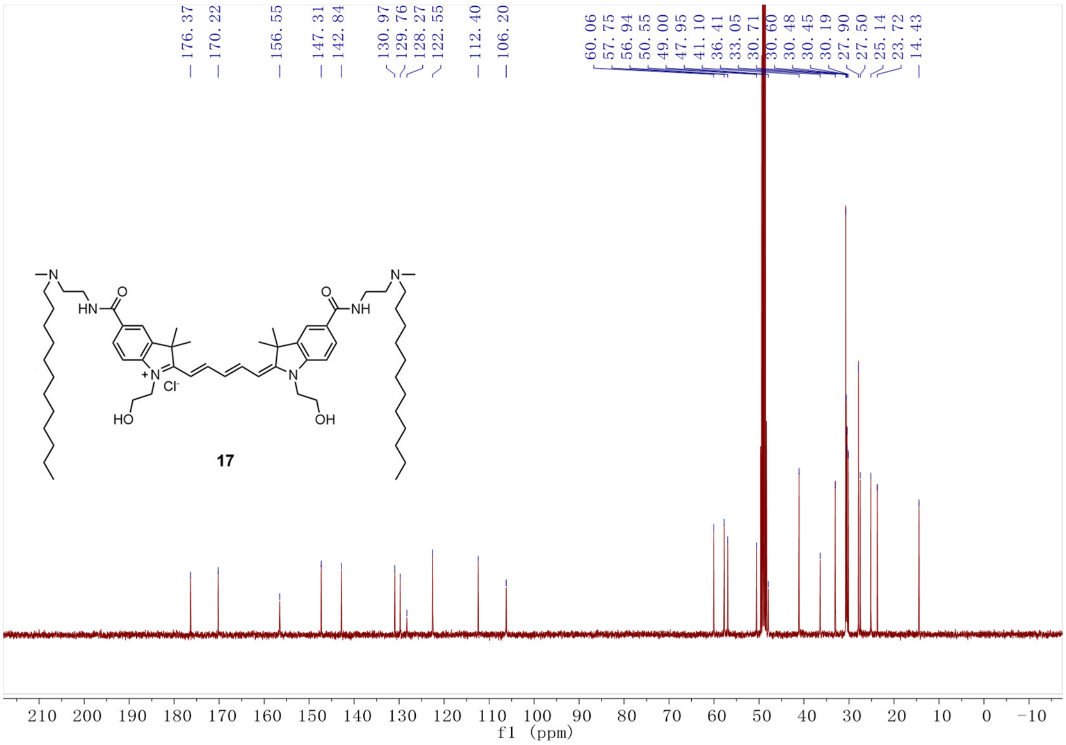

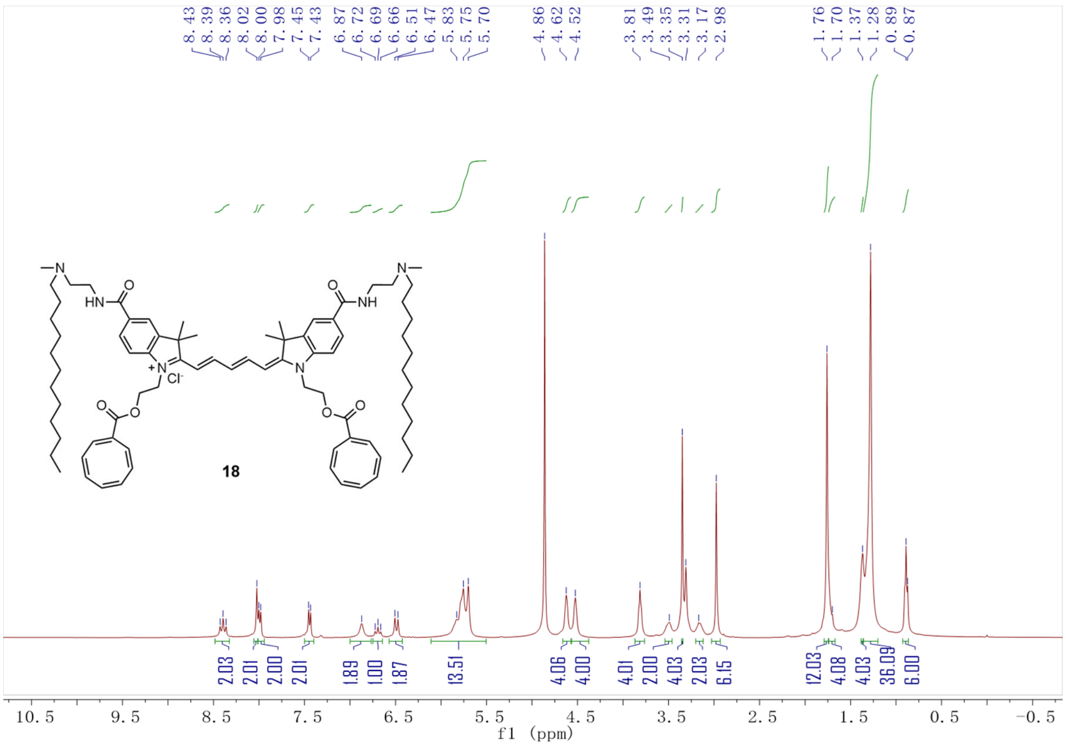
8.43 8.39 8.36 8.02 8.00 7.98 7.45 7.43 6.87 6.72 6.69 6.66 6.51 6.47 5.83 5.75 5.70 4.86 4.62 4.52 3.81 3.49 3.35 3.31 3.17 2.98 1.76 1.70 1.37 1.28 0.89 0.87
2.03 2.01 2.00 2.01 1.89 1.00 1.87 13.51 4.06 4.00 4.01 2.00 4.03 2.03 6.15 12.03 4.08 4.03 36.09 6.00
18
10.5 9.5 8.5 7.5 6.5 5.5 4.5 3.5 2.5 1.5 0.5 -0.5
f1 (ppm)

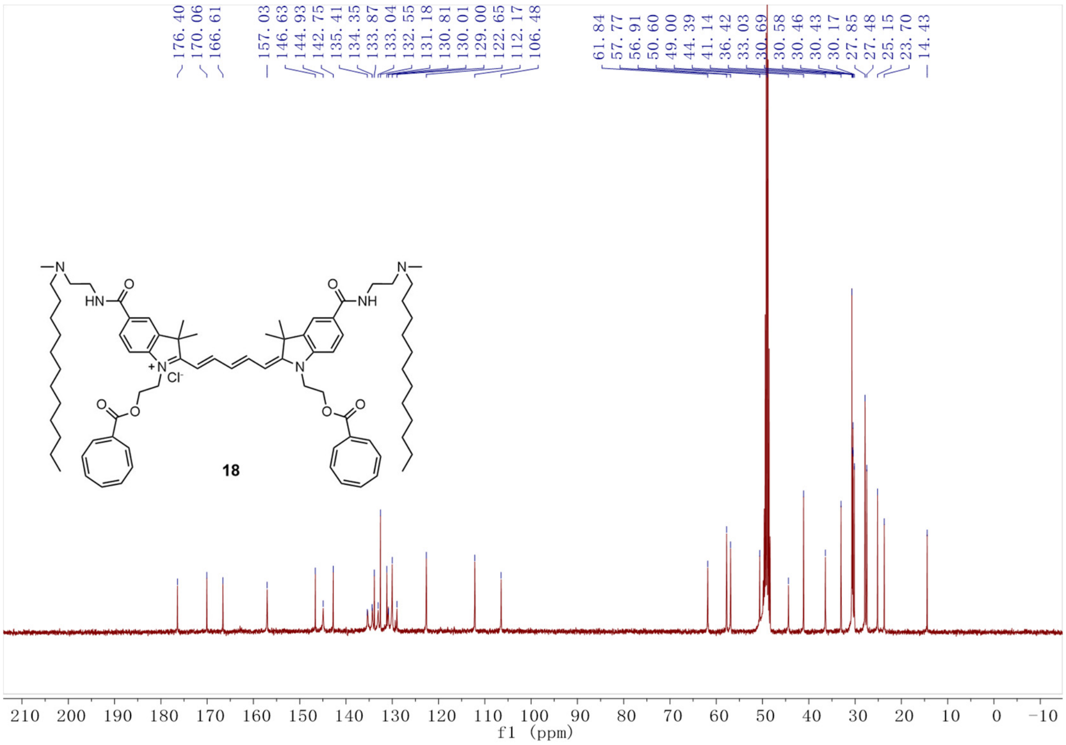
176.40 170.06 166.61 157.03 146.63 144.93 142.75 135.41 134.35 133.87 133.04 132.55 131.18 130.81 130.01 129.00 122.65 112.17 106.48
61.84 57.77 56.91 50.60 49.00 44.39 41.14 36.42 33.03 30.69 30.58 30.46 30.43 30.17 27.85 27.48 25.15 23.70 14.43
18
210 200 190 180 170 160 150 140 130 120 110 100 90 80 70 60 50 40 30 20 10 0 -10
f1 (ppm)

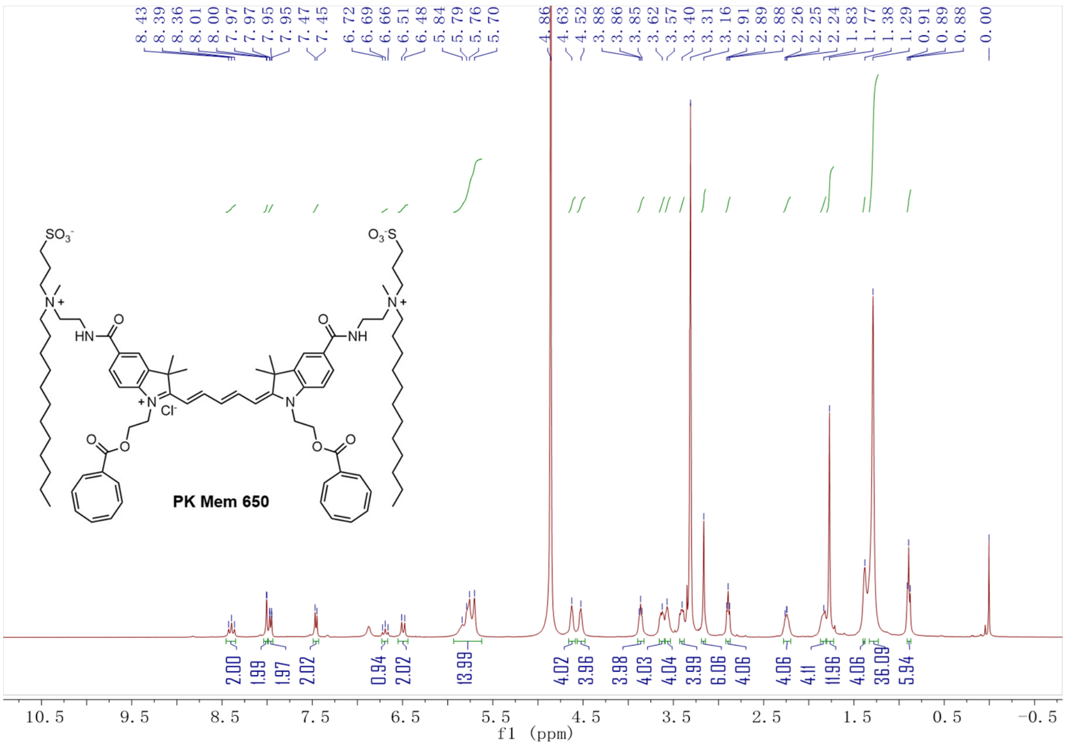
PK Mem 650
f1 (ppm)

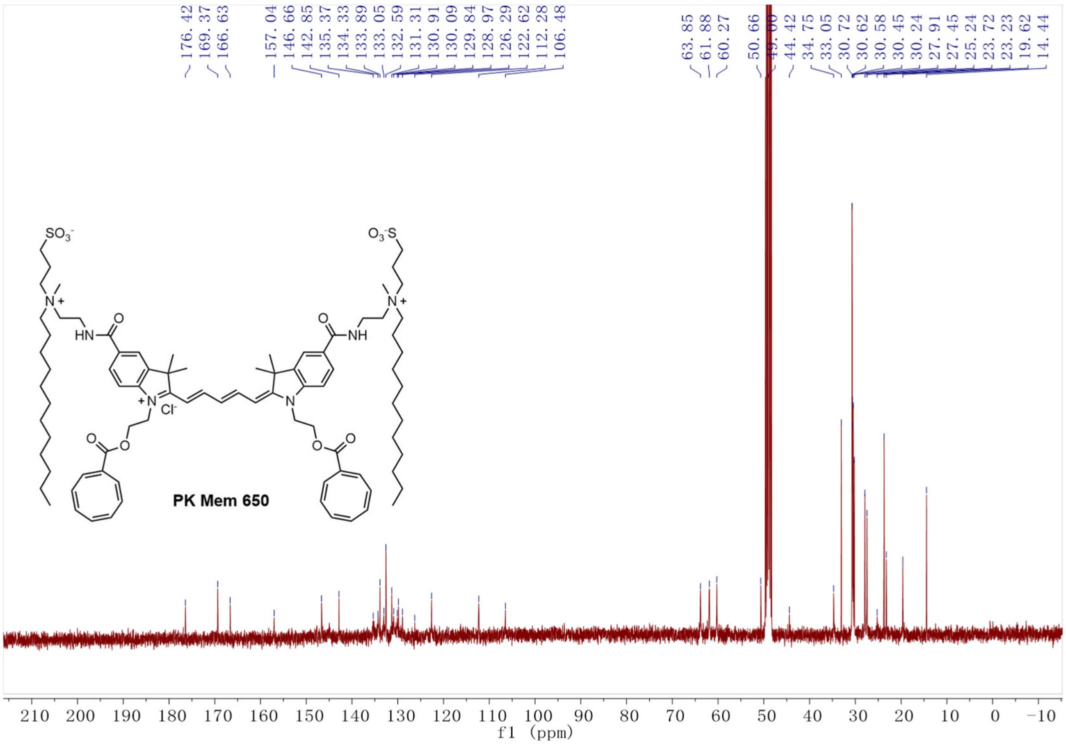
PK Mem 650
f1 (ppm)

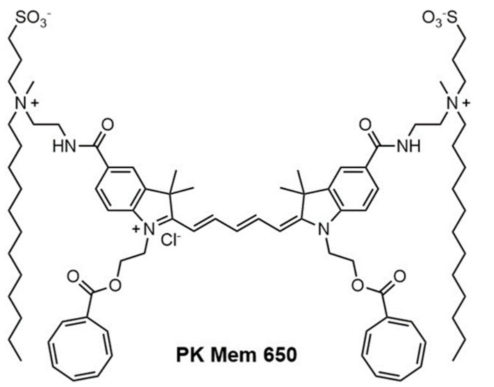

PK Mem 650

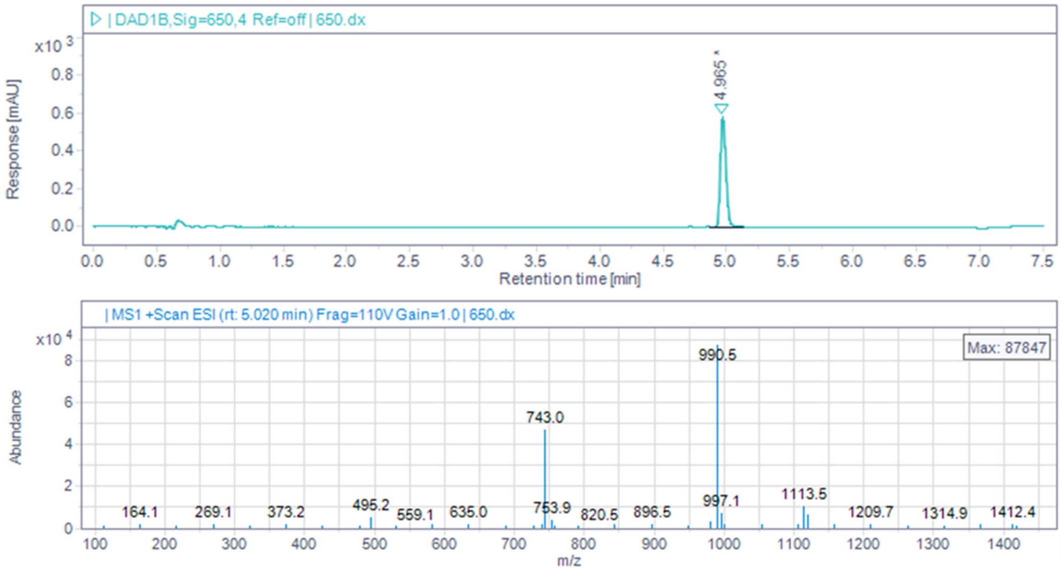

| DAD1B,Sig=650,4 Ref=off | 650.dx
x10 3
Response [mAU]
4.965
Retention time [min]
| MS1 +Scan ESI (rt: 5.020 min) Frag=110V Gain=1.0 | 650.dx
x10 4
Abundance
Max: 87847
990.5
743.0
164.1
269.1
373.2
495.2
559.1
635.0
753.9
820.5
896.5
997.1
1113.5
1209.7
1314.9
1412.4
m/z

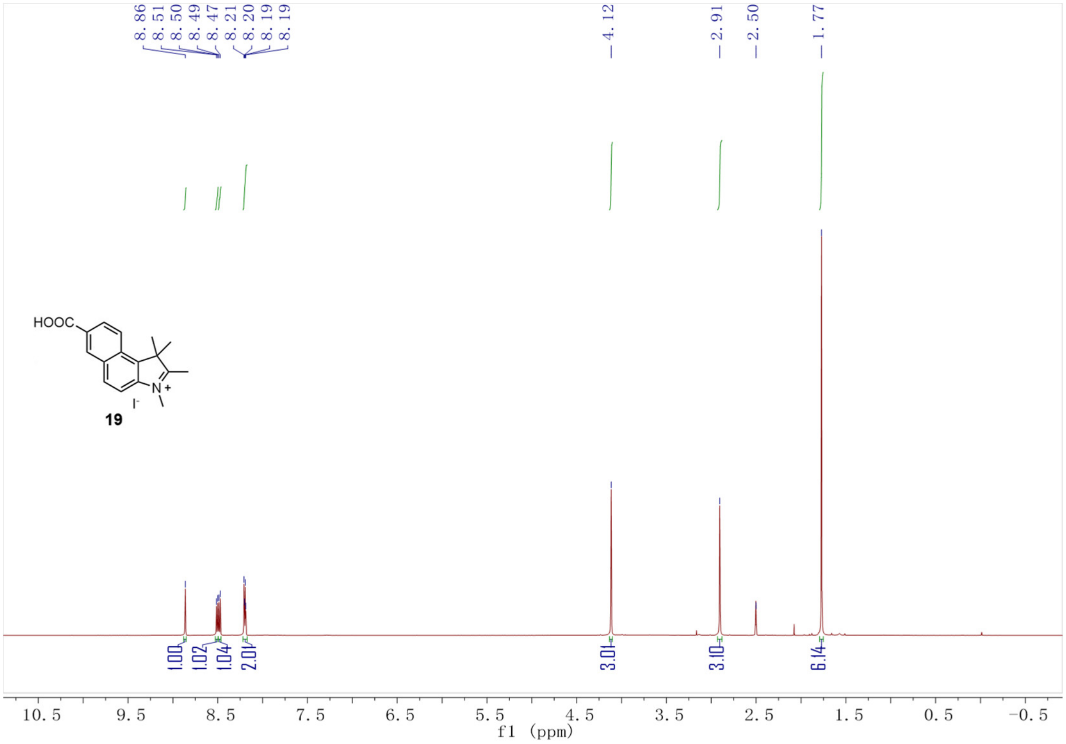
8.86
8.51
8.50
8.49
8.47
8.21
8.20
8.19
8.19
4.12
2.91
2.50
1.77
HOOC
N+
I-
19
1.00
1.02
1.04
2.01
3.01
3.10
6.14
10.5 9.5 8.5 7.5 6.5 5.5 4.5 3.5 2.5 1.5 0.5 -0.5
f1 (ppm)

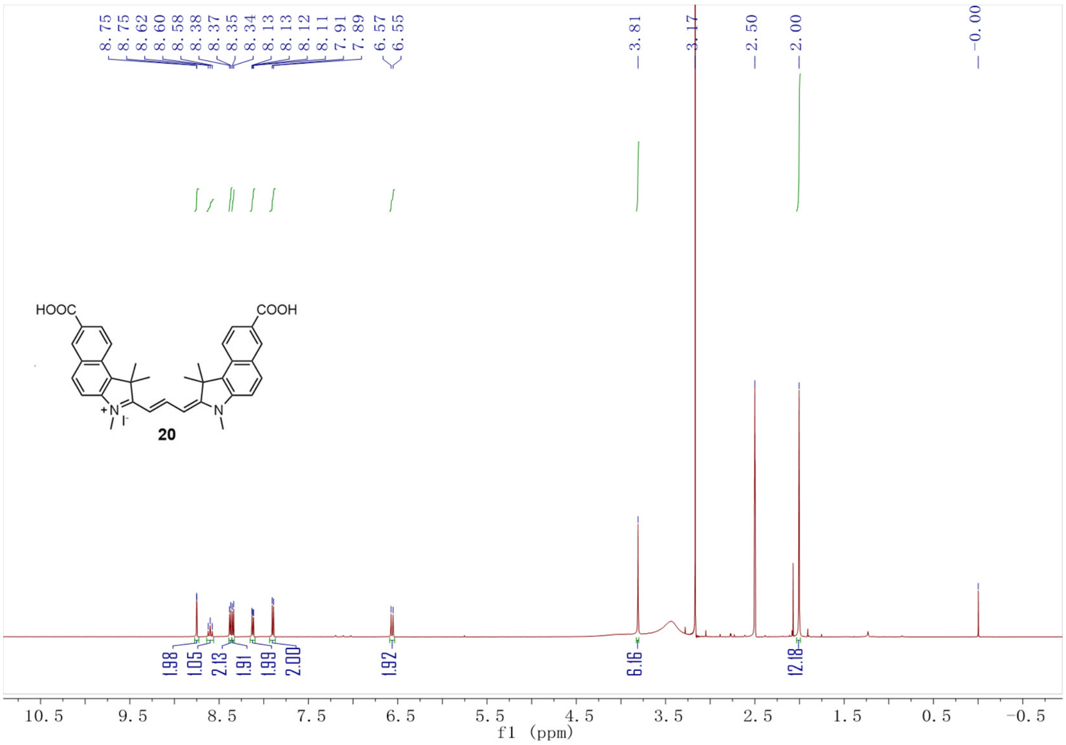
8.75
8.75
8.62
8.60
8.58
8.38
8.37
8.35
8.34
8.13
8.13
8.12
8.11
7.91
7.89
6.57
6.55
3.81
3.17
2.50
2.00
-0.00
HOOC
COOH
+N
I-
N
20
1.98
1.05
2.13
1.91
1.99
2.00
1.92
6.16
12.18
10.5 9.5 8.5 7.5 6.5 5.5 4.5 3.5 2.5 1.5 0.5 -0.5
f1 (ppm)

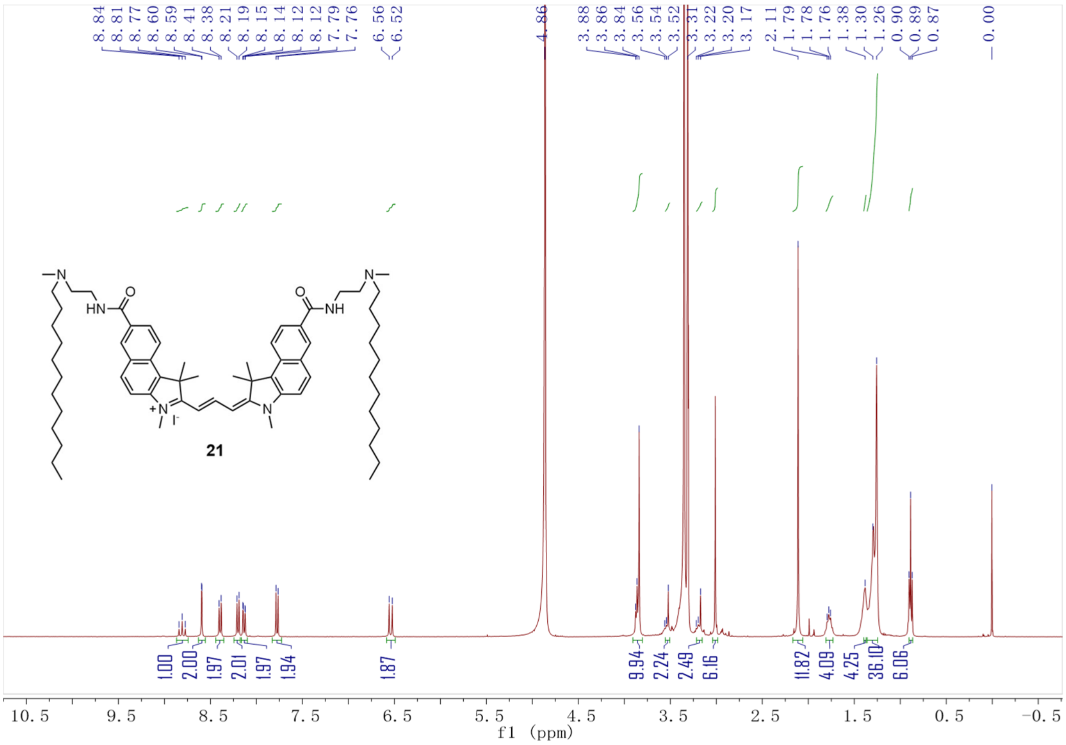
21
f1 (ppm)

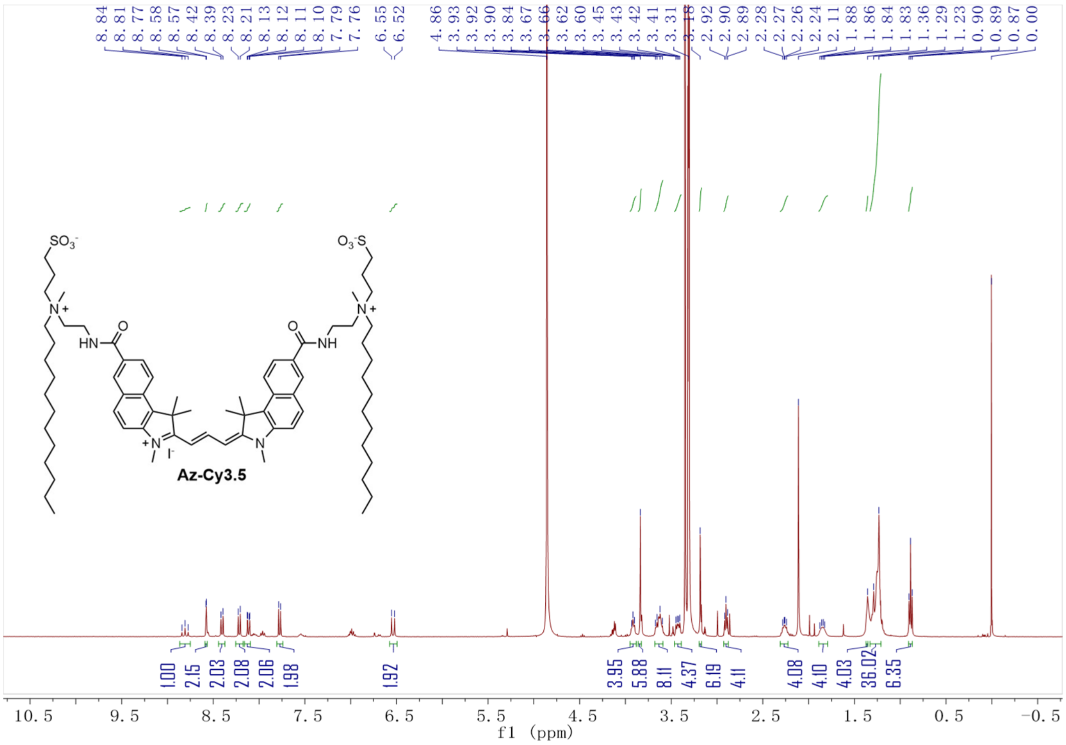
Az-Cy3.5
f1 (ppm)

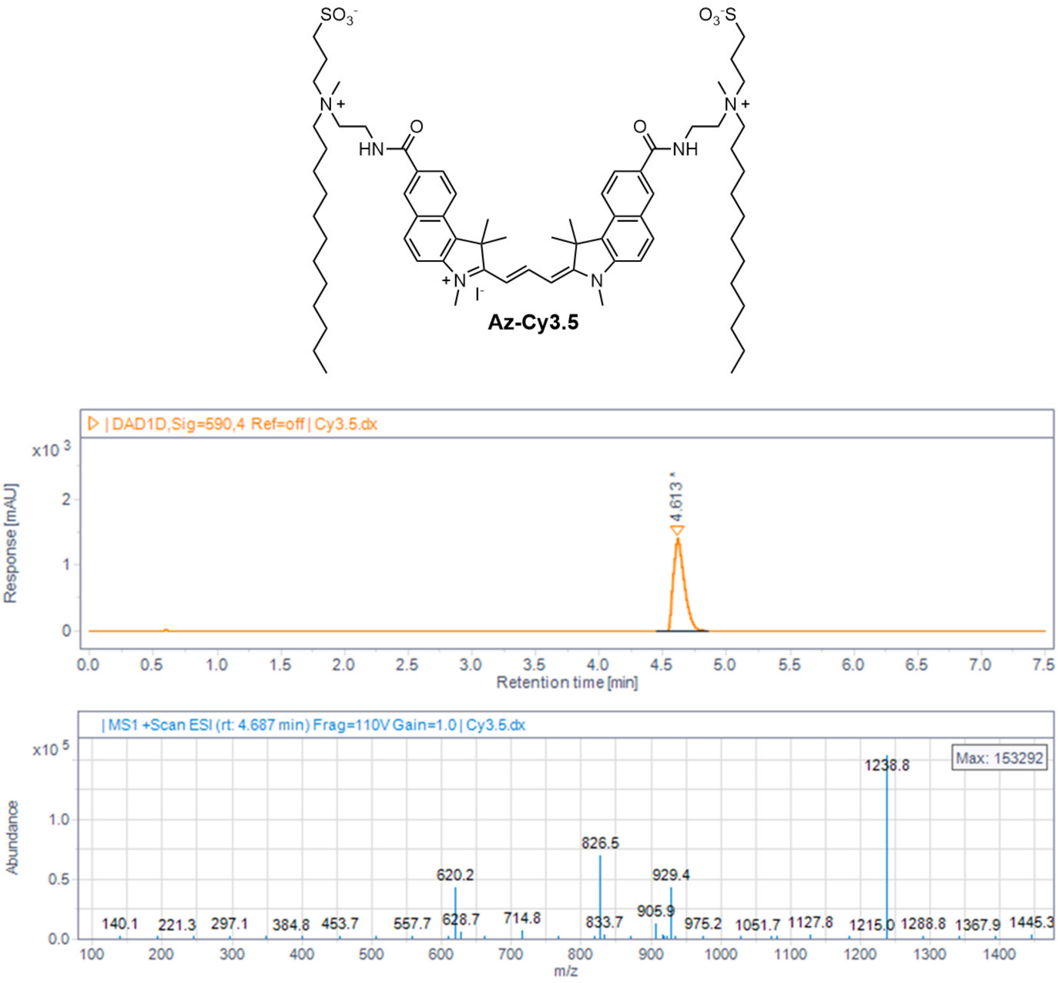
Az-Cy3.5
| DAD1D,Sig=590,4 Ref=off | Cy3.5.dx
4.613
Response [mAU]
Retention time [min]
| MS1 +Scan ESI (rt: 4.687 min) Frag=110V Gain=1.0 | Cy3.5.dx
Max: 153292
1238.8
826.5
620.2
929.4
Abundance
m/z

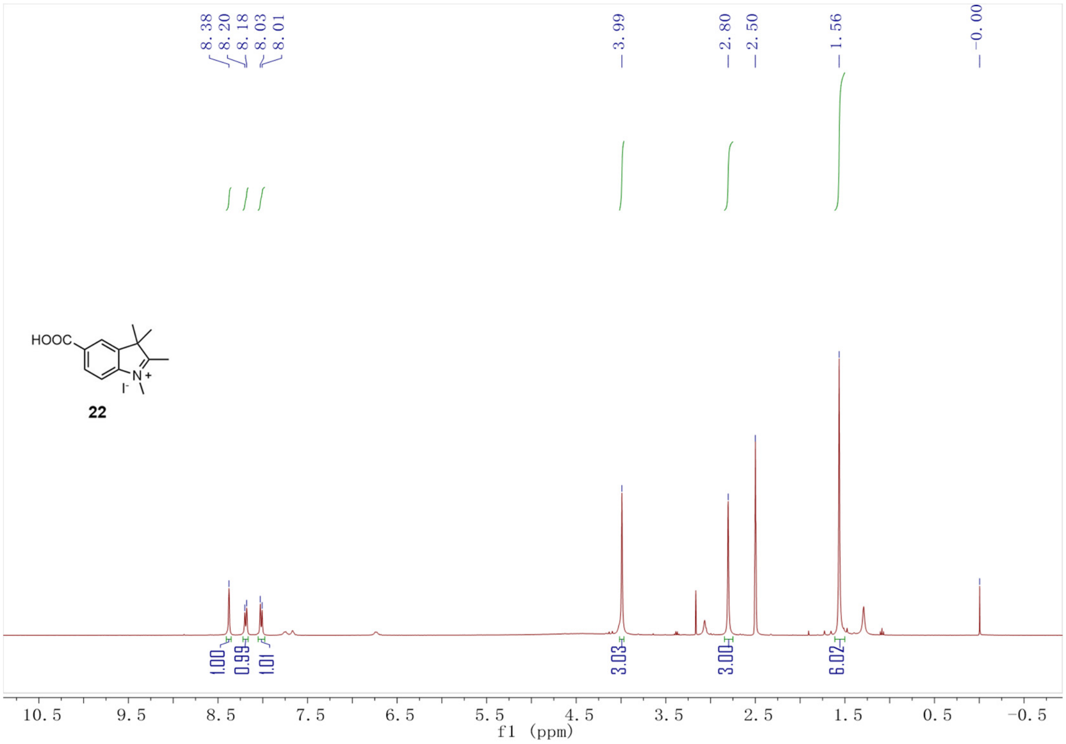
8.38
8.20
8.18
8.03
8.01
3.99
2.80
2.50
1.56
-0.00
HOOC
N+
I-
22
1.00
0.99
1.01
3.03
3.00
6.02
10.5
9.5
8.5
7.5
6.5
5.5
4.5
3.5
2.5
1.5
0.5
-0.5
f1 (ppm)

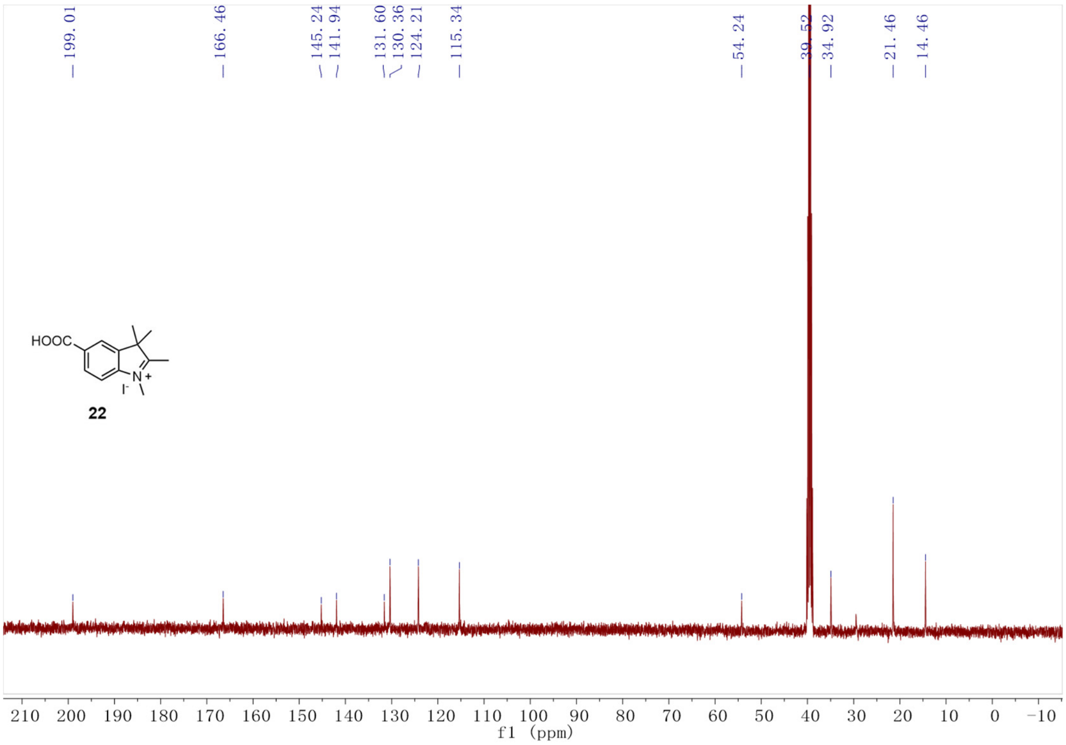
199.01
166.46
145.24
141.94
131.60
130.36
124.21
115.34
54.24
39.52
34.92
21.46
14.46
HOOC
N+
I-
22
210
200
190
180
170
160
150
140
130
120
110
100
90
80
70
60
50
40
30
20
10
0
-10
f1 (ppm)

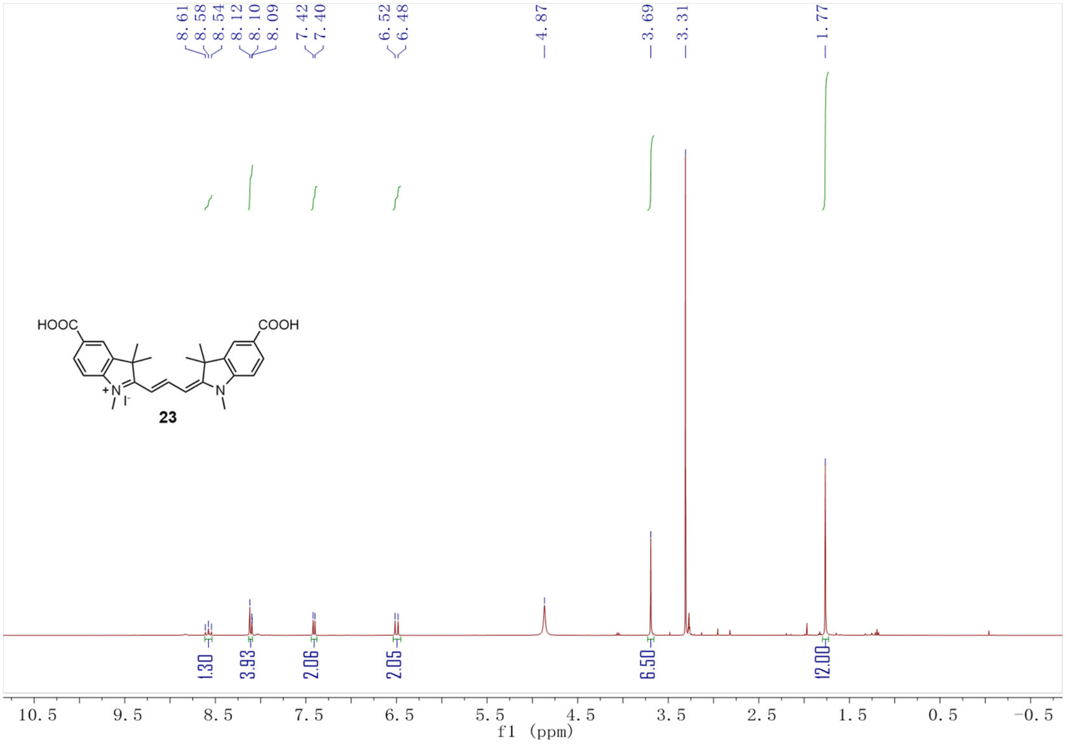
8.61
8.58
8.54
8.12
8.10
8.09
7.42
7.40
6.52
6.48
4.87
3.69
3.31
1.77
HOOC
COOH
N
I⁻
N
23
1.30
3.93
2.06
2.05
6.50
12.00
10.5
9.5
8.5
7.5
6.5
5.5
4.5
3.5
2.5
1.5
0.5
-0.5
f1 (ppm)

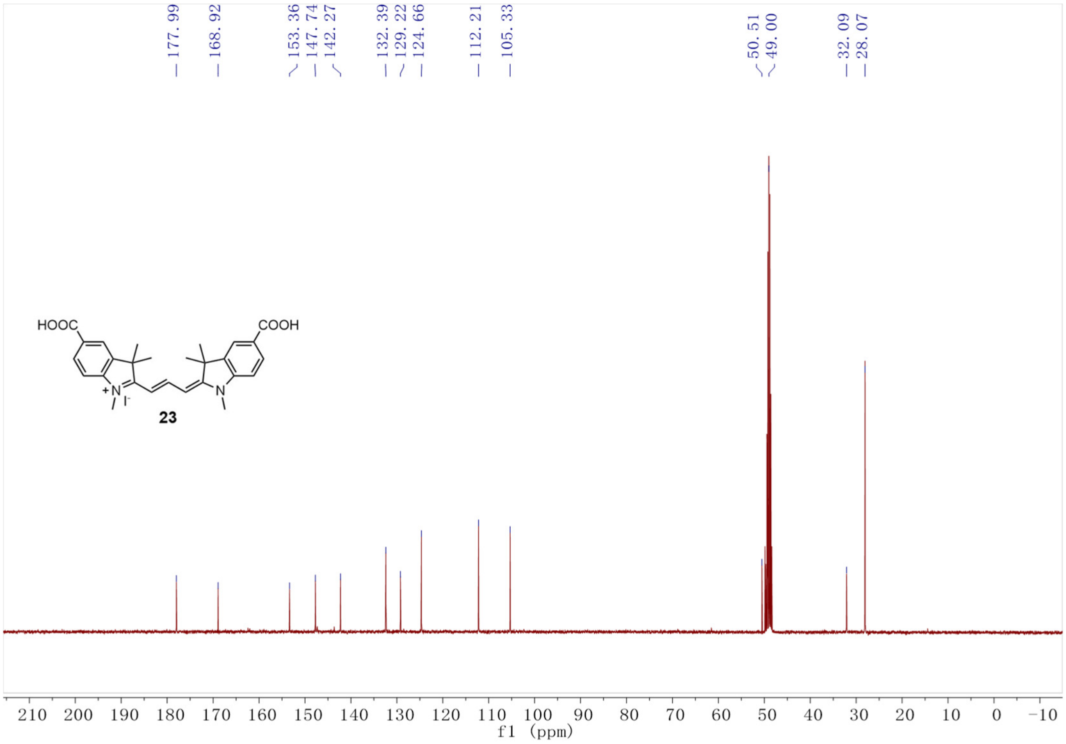
177.99
168.92
153.36
147.74
142.27
132.39
129.22
124.66
112.21
105.33
50.51
49.00
32.09
28.07
HOOC
COOH
N
I⁻
N
23
210
200
190
180
170
160
150
140
130
120
110
100
90
80
70
60
50
40
30
20
10
0
-10
f1 (ppm)

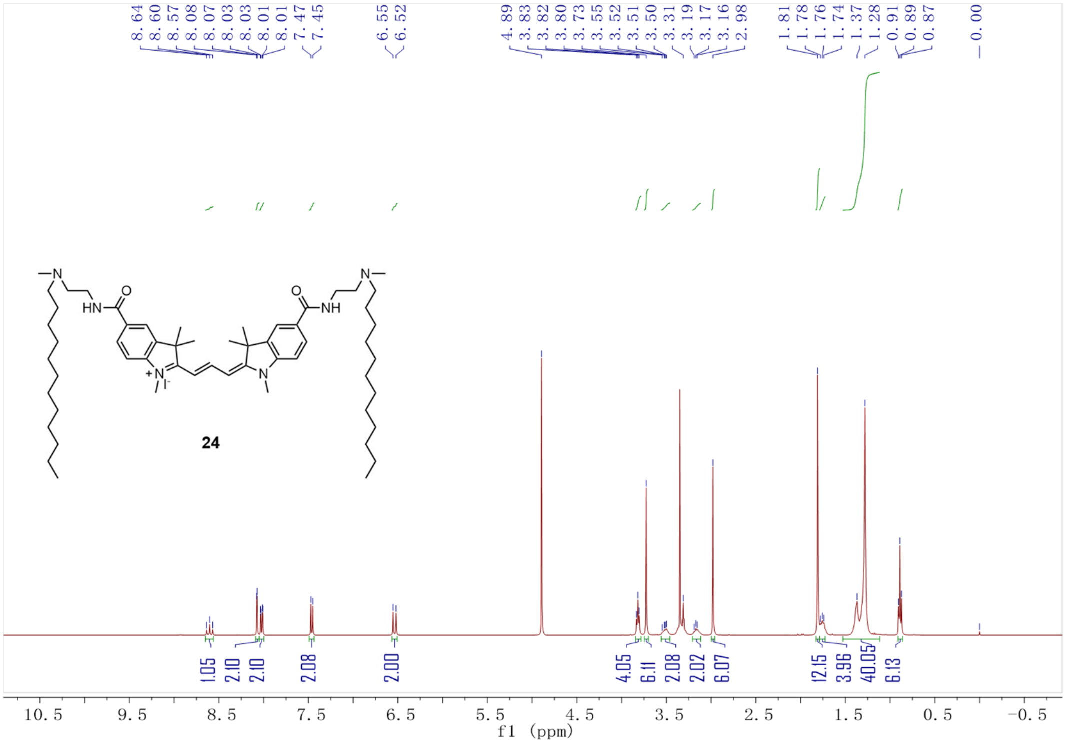
8.64 8.60 8.57 8.08 8.07 8.03 8.03 8.01 8.01 7.47 7.45 6.55 6.52 4.89 3.83 3.82 3.80 3.73 3.55 3.52 3.51 3.50 3.31 3.19 3.17 3.16 2.98 1.81 1.78 1.76 1.74 1.37 1.28 0.91 0.89 0.87 0.00
24
1.05 2.10 2.10 2.08 2.00 4.05 6.11 2.08 2.02 6.07 12.15 3.96 40.05 6.13
10.5 9.5 8.5 7.5 6.5 5.5 4.5 3.5 2.5 1.5 0.5 -0.5
f1 (ppm)

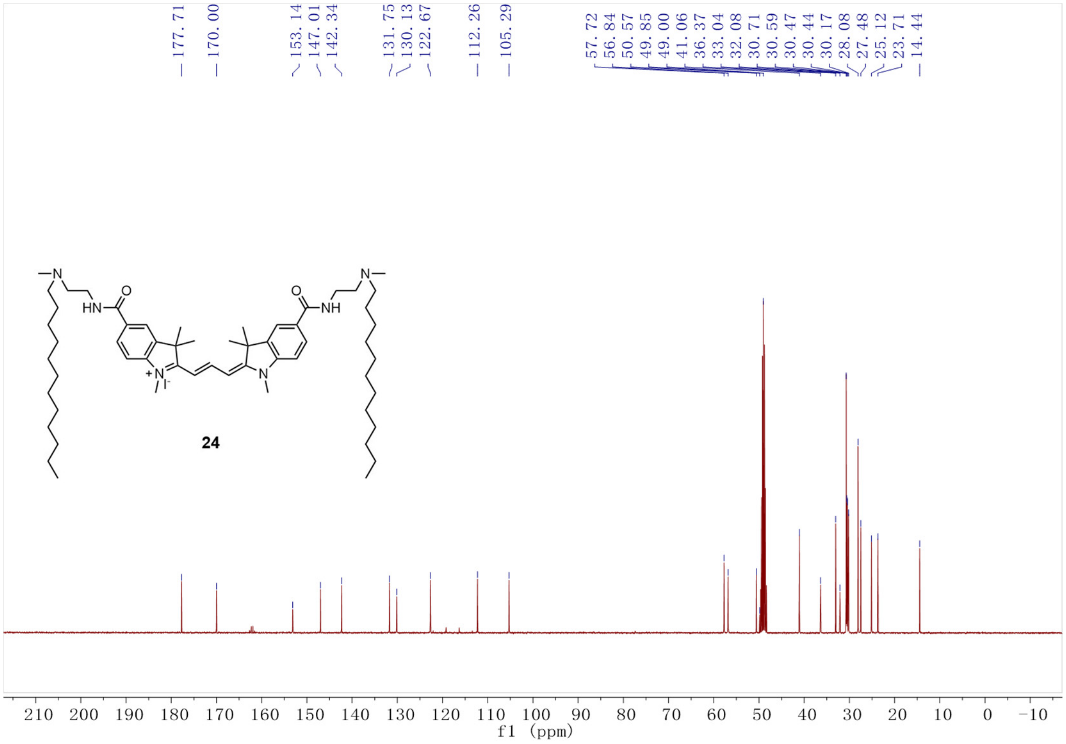
177.71 170.00 153.14 147.01 142.34 131.75 130.13 122.67 112.26 105.29 57.72 56.84 50.57 49.85 49.00 41.06 36.37 33.04 32.08 30.71 30.59 30.47 30.44 30.17 28.08 27.48 25.12 23.71 14.44
24
210 200 190 180 170 160 150 140 130 120 110 100 90 80 70 60 50 40 30 20 10 0 -10
f1 (ppm)

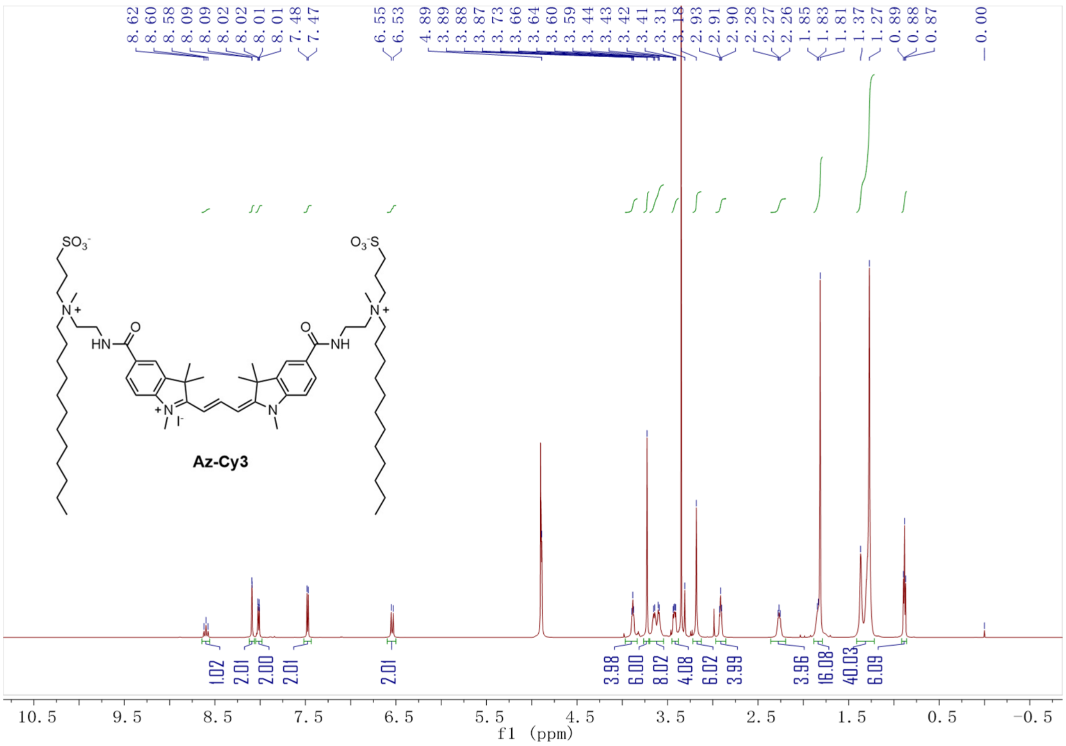
8.62 8.60 8.58 8.09 8.09 8.02 8.02 8.01 8.01 7.48 7.47 6.55 6.53 4.89 3.89 3.88 3.87 3.73 3.66 3.64 3.60 3.59 3.44 3.43 3.42 3.41 3.31 3.18 2.93 2.91 2.90 2.28 2.27 2.26 1.85 1.83 1.81 1.37 1.27 0.89 0.88 0.87 0.00
1.02 2.01 2.00 2.01 2.01 3.98 6.00 8.02 4.08 6.02 3.99 3.96 16.08 40.03 6.09
Az-Cy3
f1 (ppm)

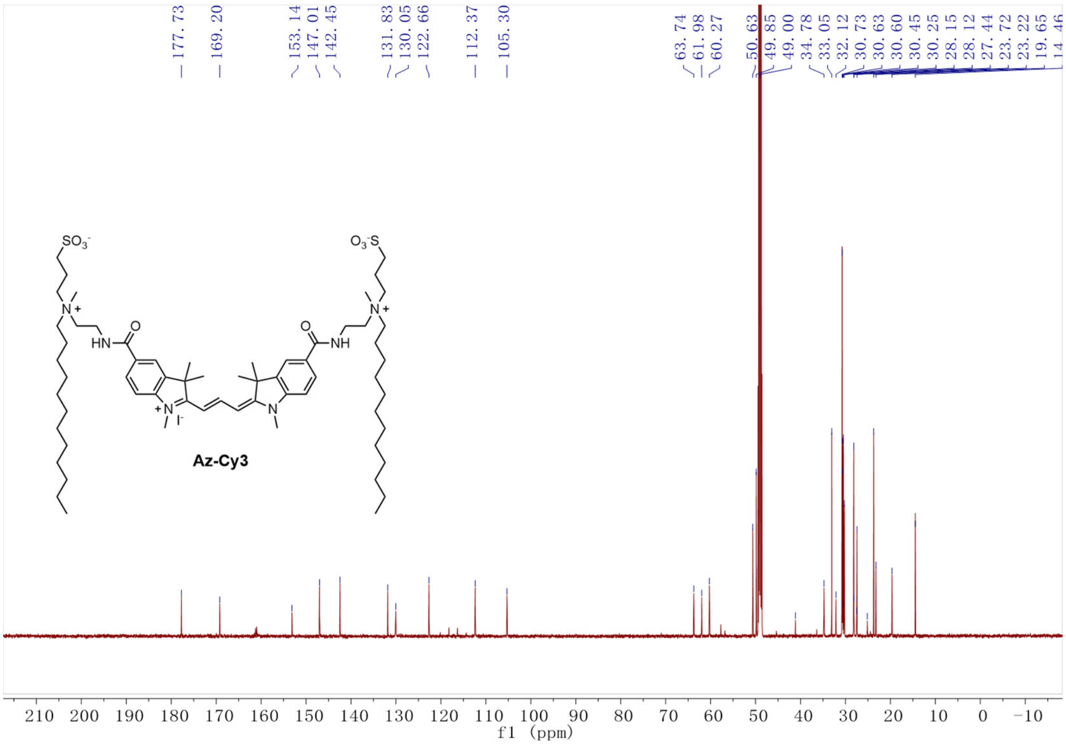
177.73 169.20 153.14 147.01 142.45 131.83 130.05 122.66 112.37 105.30 63.74 61.98 60.27 50.63 49.85 49.00 34.78 33.05 32.12 30.73 30.63 30.60 30.45 30.25 28.15 28.12 27.44 23.72 23.22 19.65 14.46
Az-Cy3
f1 (ppm)

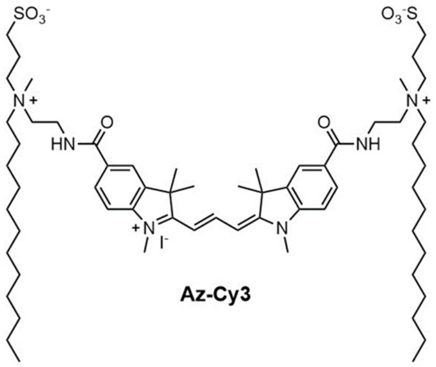
SO3-
O3-S
N+
HN
NH
O
I-
N

**Az-Cy3**

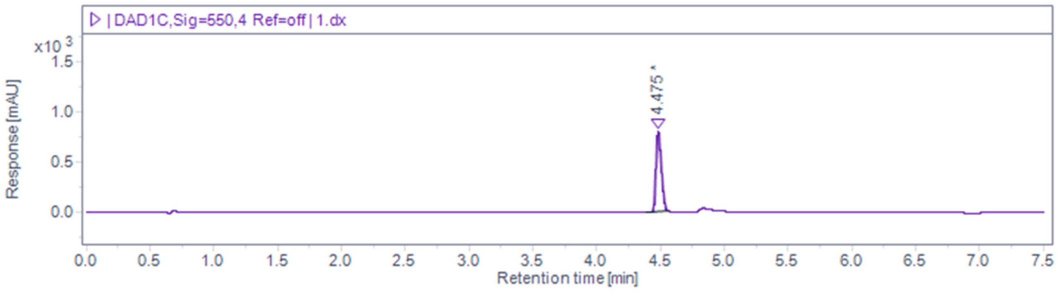
| DAD1C,Sig=550,4 Ref=off | 1.dx
x10 3
Response [mAU]
4.475
Retention time [min]

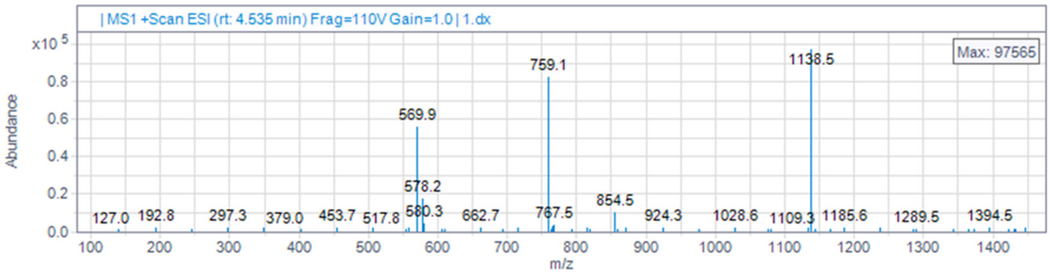
| MS1 +Scan ESI (rt: 4.535 min) Frag=110V Gain=1.0 | 1.dx
x10 5
Abundance
Max: 97565
1138.5
759.1
569.9
578.2
580.3
854.5
767.5
127.0
192.8
297.3
379.0
453.7
517.8
662.7
924.3
1028.6
1109.3
1185.6
1289.5
1394.5
m/z

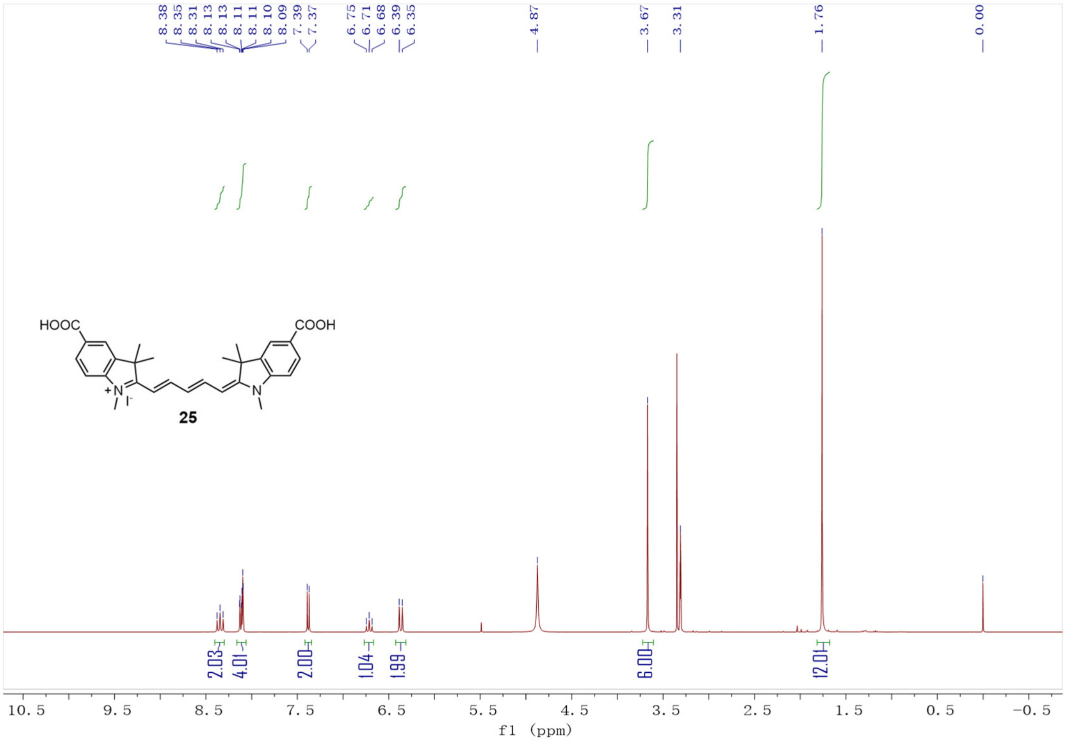
8.38
8.35
8.31
8.13
8.13
8.11
8.11
8.10
8.09
7.39
7.37
6.75
6.71
6.68
6.39
6.35
4.87
3.67
3.31
1.76
0.00
HOOC
COOH
25
2.03
4.01
2.00
1.04
1.99
6.00
12.01
10.5 9.5 8.5 7.5 6.5 5.5 4.5 3.5 2.5 1.5 0.5 -0.5
f1 (ppm)

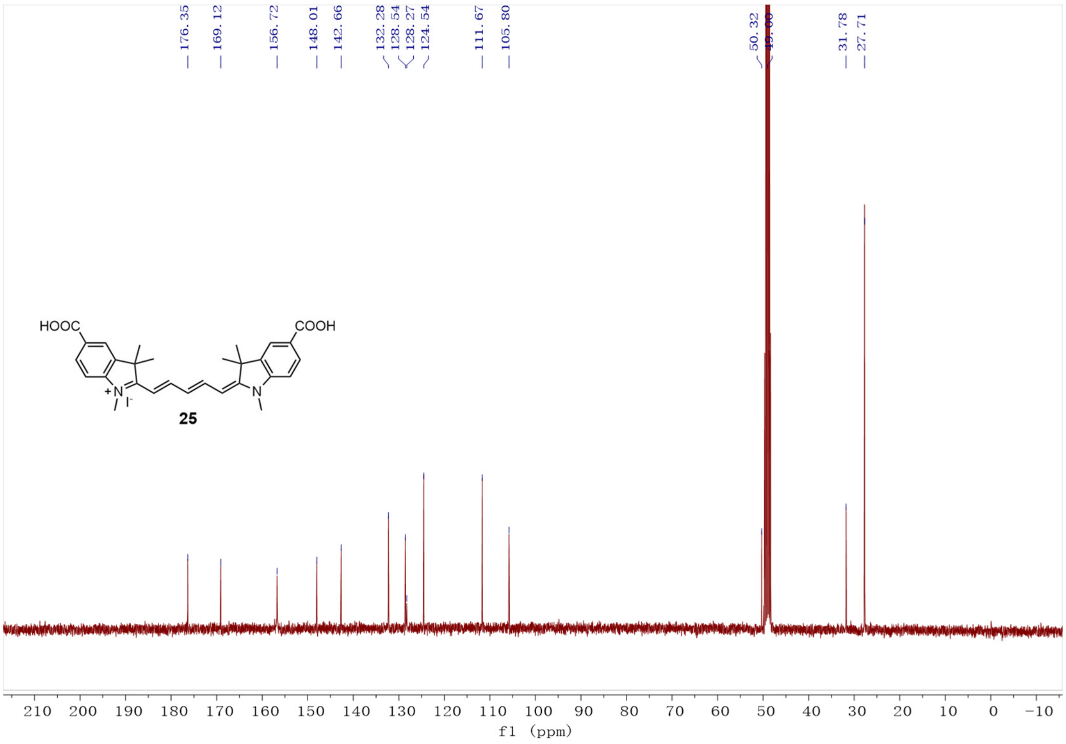
176.35
169.12
156.72
148.01
142.66
132.28
128.54
128.27
124.54
111.67
105.80
50.32
31.78
27.71
HOOC
COOH
25
210 200 190 180 170 160 150 140 130 120 110 100 90 80 70 60 50 40 30 20 10 0 -10
f1 (ppm)

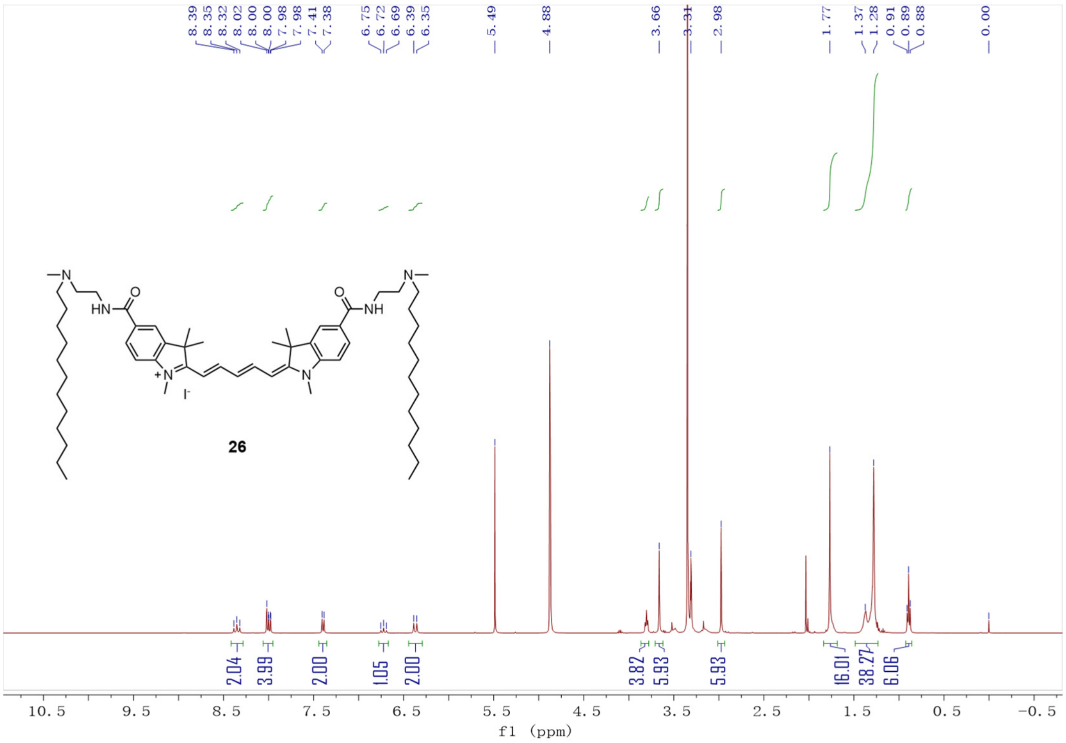
26
f1 (ppm)

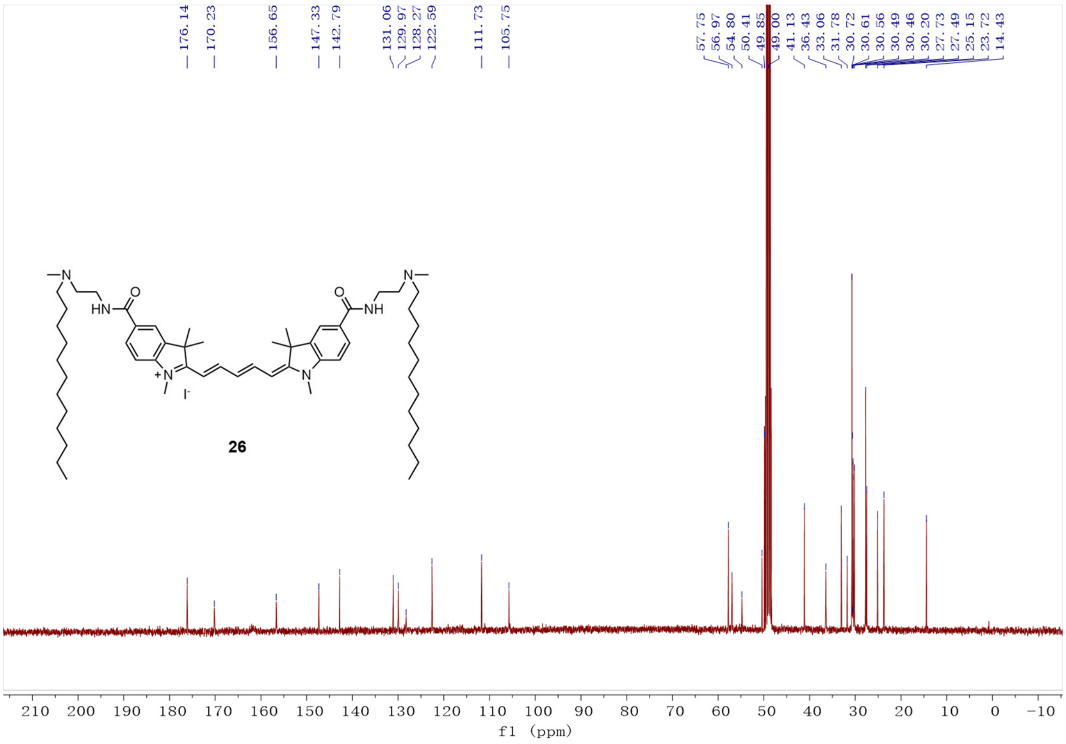
26
f1 (ppm)

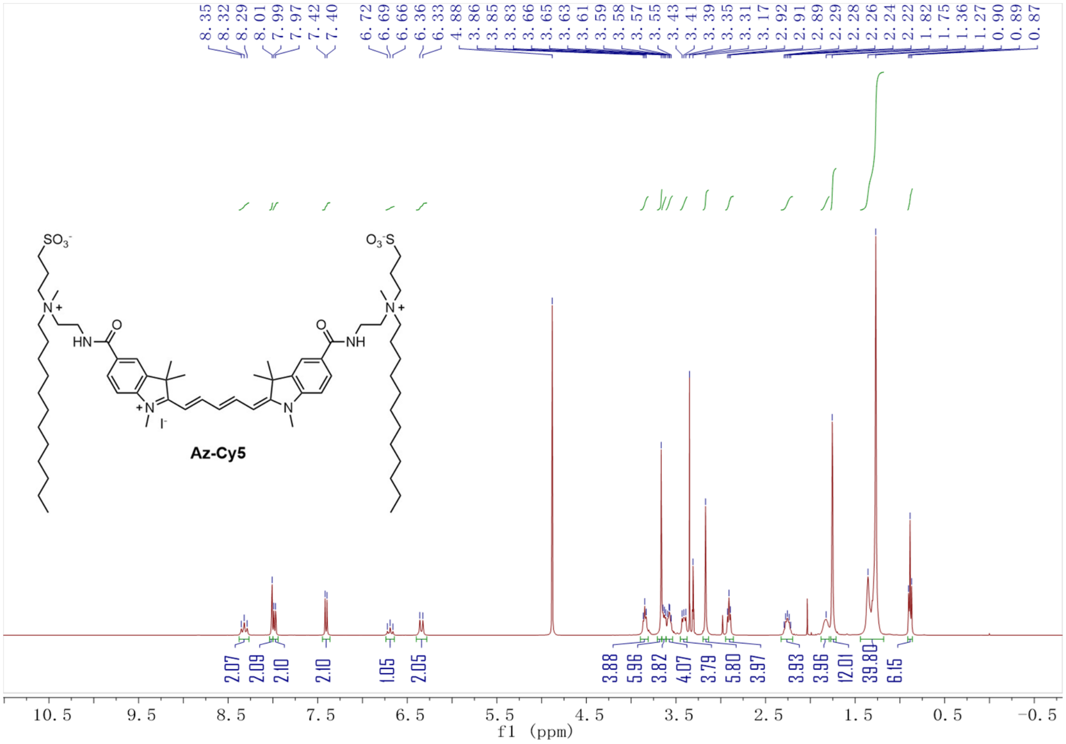
8.35
8.32
8.29
8.01
7.99
7.97
7.42
7.40
6.72
6.69
6.66
6.36
6.33
4.88
3.86
3.85
3.83
3.66
3.65
3.63
3.61
3.59
3.58
3.57
3.55
3.43
3.41
3.39
3.35
3.31
3.17
2.92
2.91
2.89
2.29
2.28
2.26
2.24
2.22
1.82
1.75
1.36
1.27
0.90
0.89
0.87
SO3-
O3-S
N+
HN
O
NH
N+
I-
N
Az-Cy5
2.07
2.09
2.10
2.10
1.05
2.05
3.88
5.96
3.82
4.07
3.79
5.80
3.97
3.93
3.96
12.01
39.80
6.15
10.5
9.5
8.5
7.5
6.5
5.5
4.5
3.5
2.5
1.5
0.5
-0.5
f1 (ppm)

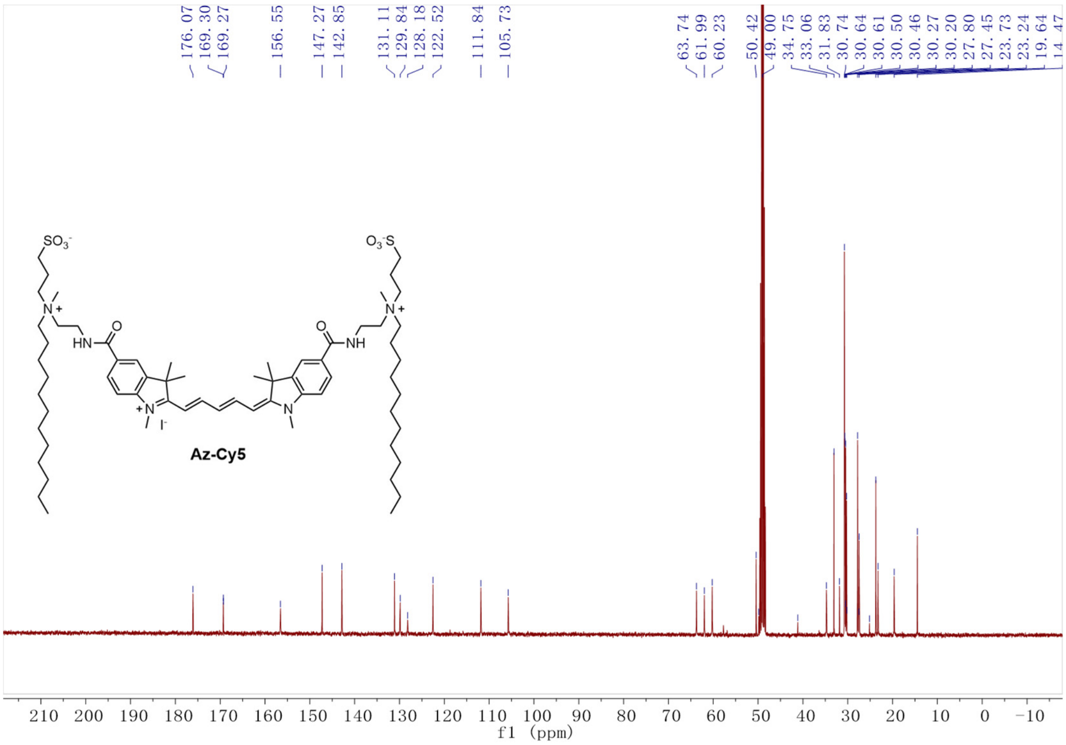
176.07
169.30
169.27
156.55
147.27
142.85
131.11
129.84
128.18
122.52
111.84
105.73
63.74
61.99
60.23
50.42
49.00
34.75
33.06
31.83
30.74
30.64
30.61
30.50
30.46
30.27
30.20
27.80
27.45
23.73
23.24
19.64
14.47
SO3-
O3-S
N+
HN
O
NH
N+
I-
N
Az-Cy5
210
200
190
180
170
160
150
140
130
120
110
100
90
80
70
60
50
40
30
20
10
0
-10
f1 (ppm)

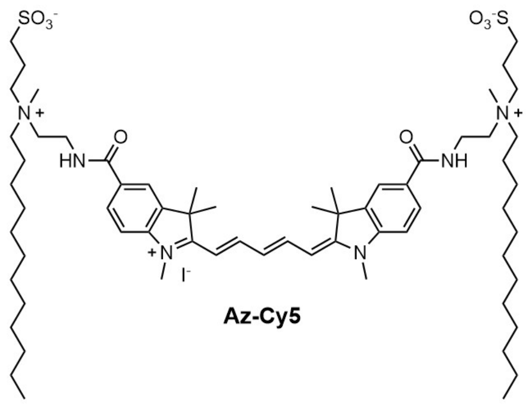
SO3-
O3-S
N+
HN
NH
O
+N
I-
N

**Az-Cy5**

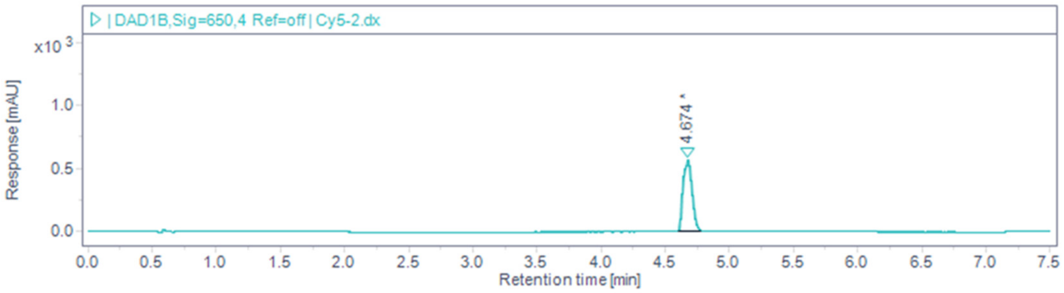
| DAD1B,Sig=650,4 Ref=off | Cy5-2.dx
x10 3
Response [mAU]
4.674
Retention time [min]

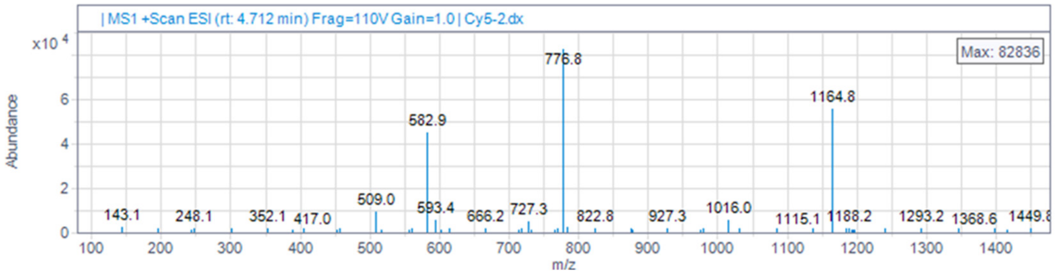
| MS1 +Scan ESI (rt: 4.712 min) Frag=110V Gain=1.0 | Cy5-2.dx
x10 4
Abundance
Max: 82836
776.8
582.9
1164.8
509.0
593.4
727.3
1016.0
143.1
248.1
352.1
417.0
666.2
822.8
927.3
1115.1
1188.2
1293.2
1368.6
m/z
